# Characterization of ovine follicular fluid and granulosa cell-derived extracellular vesicles and their miRNA cargo following in vitro exposure to bisphenols A and S

**DOI:** 10.64898/2026.03.27.713654

**Authors:** Alice Desmarchais, Svetlana Uzbekova, Virginie Maillard, Pascal Papillier, Cécile Douet, Thomas Duret, Rustem Uzbekov, Quentin Meffre, Benoît Piégu, Gaëlle Lefort, Noémie Teixido, Anaïs Vitorino Carvalho, Sébastien Roger, Sébastien Elis

**Author notes:** Corresponding author: Alice Desmarchais.

## Abstract

Bisphenol A (BPA) and Bisphenol S (BPS) exposure disrupt ovarian function and granulosa cell (GC) steroidogenesis. Extracellular vesicles (EVs) and their miRNA cargo, as mediators of cellular response to environmental stimuli, might be involved in fertility and folliculogenesis. This study explored modulation of microRNA expression after 48h BPA or BPS exposure (10 µM) in ovine primary GC and EVs from corresponding conditioned medium (CM EVs). Small RNA sequencing of control (0h) and 48h treated GC, CM EVs as well as follicular fluid EVs allowed identification of 533 ovine miRNAs, including 129 new sequences. BPA did not alter miRNA expression in GC, while BPS decreased cellular oar-24b miR. In contrast, BPA modified expression of 4 miRNAs in CM-EVs, including 3 new sequences, and two miRNAs were modified by BPS. Both compounds reduced expression of sequence homologous to miR-1306. Further studies are required to decipher their roles in bisphenol toxicity in GC.

## 1. Background

During folliculogenesis, the oocyte develops within the follicle, which provides the necessary environment for its growth and acquisition of competence for fertilization and early embryo development. This optimal environment is provided particularly by the surrounding follicular granulosa cells (GC), which enable follicular growth and steroidogenesis (synthesis and secretion of estradiol and progesterone), but also by the follicular fluid (FF). FF fills the antrum and is composed of secretion products from follicular cells and molecules supplied by blood vessels such as ions, hormones, growth factors, metabolic precursors (Monniaux et al., 2019).

Ovarian follicle growth, along with oocyte development and maturation, also requires constant, tight communications between the oocyte and follicular cells such as GC and cumulus cells (Dalbies-Tran et al., 2020; Monniaux et al., 2019). This communication includes the exchange of biomolecules like nucleic acids or proteins and occurs through direct transfer between adjacent follicular cells (Collado et al., 2018) mediated by extracellular vesicles (EVs) (Marchais et al., 2022; Raposo and Stoorvogel, 2013; Smith and Russell, 2022). These EVs carry a variety of biomolecules, including proteins, lipids, messenger RNAs (mRNAs) and non-coding RNAs such as microRNAs (miRNAs) (Valadi et al., 2007).

EVs are membranous lipid bilayer vesicles secreted by most cells to deliver signals to target cells. According to the latest instructions in the Minimal information for studies of extracellular vesicles (MISEV) 2023 guidelines, EVs are classified as small EVs (diameter < 200 nm) or large EVs (diameter > 200 nm) (Welsh et al., 2024). They originate either from the endosomal system, where they are released as exosomes (30 to 200 nm) after maturation in multivesicular bodies, or from the shedding of the plasma membrane as microvesicles or ectosomes (100 - 1000 nm). Follicular fluid EVs (FF EVs) (< 200 nm) or cumulus cell EVs can be taken up by granulosa cells, by blastocyst or oocytes through the zona pellucida (Da Silveira et al., 2017; Fiorentino et al., 2024; Matsuno et al., 2017; Pavani et al., 2018; Uzbekova et al., 2020). This uptake has been associated with improved oocyte maturation and embryo early development (Azari-Dolatabad et al., 2025; Da Silveira et al., 2017, 2015; Lipinska et al., 2025; Schneberger et al., 2026). FF EVs can modulate apoptosis, proliferation and steroidogenesis in bovine GC and human KGN cell line (Hung et al., 2017; Varik et al., 2025; Ying et al., 2023; Yuan et al., 2021) and have been shown to promote or facilitate cumulus expansion in cattle (Hung et al., 2015).They have been related to modification of gene expression in bovine (Sohel et al., 2013) and equine granulosa cells (Da Silveira et al., 2015, 2014). The authors suggest this effect may be linked to miRNA content within the EVs (Da Silveira et al., 2012; Sohel et al., 2013).

MiRNAs are small (19-25 bp) non coding RNA sequences which are crucial post transcriptional epigenetic regulators as they regulate a large proportion of genes (Ha and Kim, 2014) including those involved in follicle growth and in the acquisition of oocyte developmental competence (Souza et al., 2025; Alexandri et al., 2020; Machtinger et al., 2017; Tesfaye et al., 2018). Extracellular vesicles containing miRNA have been isolated from several biological fluids including follicular fluid in equine (Da Silveira et al., 2012), bovine (Capra et al., 2022; Sohel et al., 2013), human (Santonocito et al., 2014), porcine (Matsuno et al., 2017), and rhinocerotid (Gad et al., 2024) species.

Moreover, in humans, FF EV - associated miRNA were considered as a marker of IVF outcomes (Machtinger et al., 2017; Martinez et al., 2018). Therefore, EVs and their miRNA cargo could have important implication for fertility, follicular development, and oocyte competence (Fiorentino et al., 2024; Santonocito et al., 2014), especially as they could mediate cellular response to environmental stimuli. In human FF, small EVs are more likely to contain miRNAs than large EVs (Varik et al., 2025) or non-exosomal EV subpopulations (Sohel et al., 2022).

Female fertility is vulnerable to endocrine-disrupting chemicals such as bisphenol A (BPA) and bisphenol S (BPS) (Nawaz et al., 2025; Peters et al., 2024; Trela-Kobędza and Ajduk, 2025). Bisphenols (BPs) are synthetic chemicals used in the production of numerous plastic materials, including polycarbonates and epoxy resins, found in food containers, cosmetics, medical devices, paper receipts, and others (Almeida et al., 2018; Bousoumah et al., 2021; H. Zhang et al., 2019). BPs are known to alter female reproductive function (Kawa et al., 2021; Ma et al., 2019; You and Song, 2021). BPA and BPS have been detected in human biological fluids, including urine (Calafat et al., 2005; Mendy et al., 2020; Sol et al., 2021), follicular fluid (Hoffmann-Dishon et al., 2024; Lebachelier De La Riviere et al., 2023), and serum (Gao et al., 2021; Khmiri et al., 2020; Paoli et al., 2020; Raimondo et al., 2024), raising concerns about their potential impact on ovarian function and female reproduction.

Moreover, BPA and BPS impair steroidogenesis in rodent (Shi et al., 2019, 2017), porcine (Berni et al., 2019; Bujnakova Mlynarcikova and Scsukova, 2021, 2018), bovine (Campen et al., 2018), human (Amar et al., 2020; Mansur et al., 2016), and ovine GC (Téteau et al., 2023, 2020).

Environmental stimuli can cause cellular stress response and can modify EV miRNA cargo (Harischandra et al., 2017). For example, BPA can deregulate the expression of cellular and EV-associated miRNA in GC from human (Rodosthenous et al., 2019), rat (Lite et al., 2019) and bovine (Sabry et al., 2023, 2022) species.

The ovine model, as a monoovulaory species, presents several similarities with human follicular growth and is therefore relevant for studying the disruptive effects of bisphenols on reproductive physiology (Gingrich et al., 2019; Scaramuzzi et al., 2011). In the present study, we investigated whether a 48-hour exposure to BPA or BPS can alter EV secretion and the expression of cellular and EV-enriched miRNAs from ovine granulosa cells. As little is known about ovine ovarian miRNA, we also qualitatively compared the EV-enriched miRNAs found in follicular fluid to those secreted by GC in conditioned media.

## 2. Methods

### 2.1. Chemicals and Antibodies

BPA (#42088) and BPS (#43034) were purchased from Sigma-Aldrich (Saint Quentin Fallavier, France). All other chemicals were obtained from Sigma-Aldrich, unless otherwise stated in the text. The primary anti-bodies used are indicated in Additional Table 1. Horseradish peroxidase (HRP)-conjugated anti- rabbit, anti-mouse, anti-rat and anti-goat were purchased from Perkin Elmer (Courtaboeuf, France).

**Table 1.**
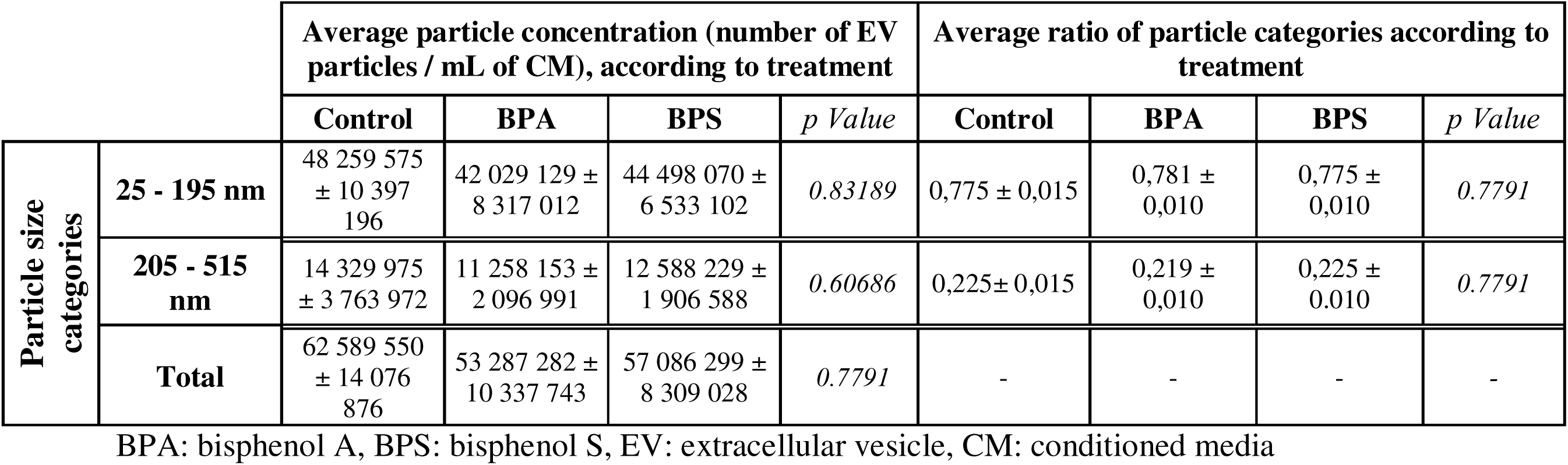
Average number of particles per mL of 48h conditioned media according to size categories. Results are presented as mean ± standard error of mean of 10 different experiments. Statistical analysis was determined with parametric analysis of variance (ANOVA) by permutation. EV = extracellular vesicle, CM = conditioned media.

### 2.2. Granulosa cell samples

Experimental design summary is presented in Figure 1. Ovaries of ewes (adult and lambs) were collected at a local slaughterhouse. Follicles (2 -6mm) were punctured with a 18G needle connected to a vacuum pumping system with a collection tube receiving follicular fluid (FF). FF was centrifuged for 5 min at 600g (Centrifuge 5702, Eppendorf, Germany). Follicular fluid supernatants were then processed as described in the « EV isolation » section. Pellets containing granulosa cells (GC) were resuspended in ACK buffer containing ammonium chloride (155 mM), potassium bicarbonate (10 mM), ethylenediaminetetraacetic acid (0.10 mM), for 3 min to lyse red blood cells. After centrifugation for 5 min at 600g, supernatants were discarded and cell pellets were washed in serum free modified McCoy’s5A medium containing L-Glutamine (3mM), bovine serum albumin (0.1%), penicillin/streptomycin (120.103 UI/L and 120 mg/L respectively), HEPES (20 mM, pH 7.6), 11βOH-4-androstenedione (96 nM), bovine apo-transferrin (5 mg/L), selenium (0.12 µM) and insulin (1.74 nM). After dropped off on a Percoll density medium (50% Percoll, 50% medium), cells were purified using centrifugation (30 min, 700 g). After washing in medium, the cells were filtered through a 100 µm cell strainer (Falcon, Corning®, USA), and suspensions were centrifuged at 600 g for 5 min. This washing and centrifugation step was repeated once. Finally, the pellets were resuspended in modified McCoy’s 5A and cells were counted using a Thoma hemocytometer with trypan blue staining. For culture, cells were plated at a density of 5 x 10^5^ viable cells per well in 1 mL modified McCoy’s 5A medium in 24-well plates (BioLite, Thermo Fisher Scientific). Cells from each batch were distributed across 36 wells and incubated overnight. The next day, the cells were treated for 48h (12 wells per condition) with either 10 µM BPA, or 10 µM BPS, or a vehicle control containing only 0.01% ethanol (171 mM). Cell cultures were maintained at 37°C in a humidified incubator with an atmosphere of 5% CO2 in air.

**Figure 1.**
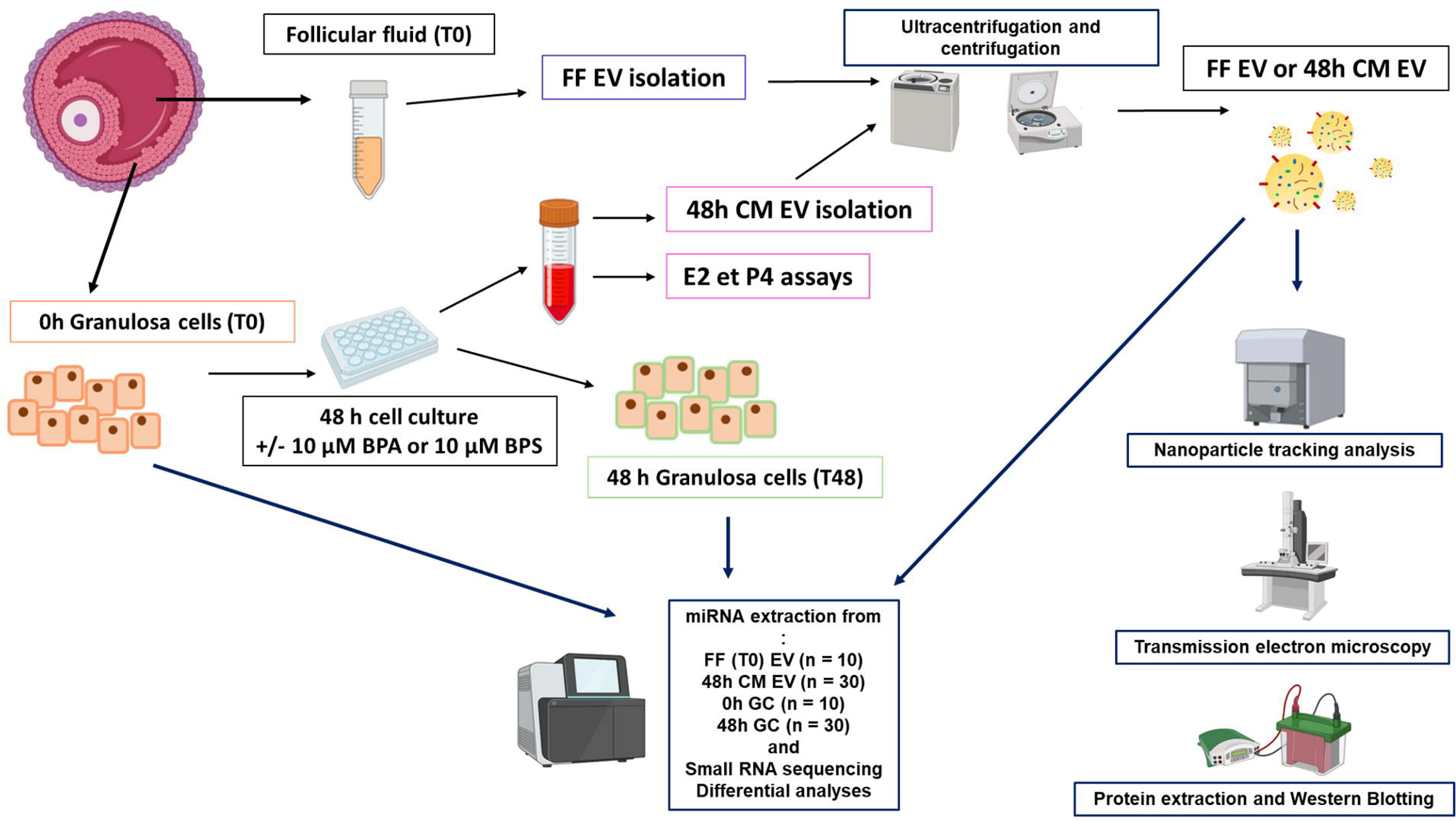
Summary of the design study. Ovine granulosa cells were isolated from ovine ovarian follicles and cultured for 48h with or without 10 µM BPA or BPS. Extracellular vesicles (EVs) were isolated from ovine follicular fluid and from the condioned media of the 48-hour granulosa cell culture. EVs were characterized using several methods: nanoparicle tracking analysis, transmission electron microscopy, western blotting and small RNA sequencing. Desmarchais A. (2026). Figure created in part with Biorender.

Aliquots of GC batches collected at the time of seeding (0h GC) and after 48h of culture (48h GC) were prepared for subsequent analyses. For RNA isolation, cells were resuspended in NucleoZOL (Macherey Nagel, Germany).

### 2.3. Steroidogenesis analysis: progesterone and estradiol assays

Aliquots (200 µL) of pooled 48-hour conditioned media were stored at -20°C until analysis. Estradiol concentration in undiluted CM samples was assessed using commercial enzyme immunoassay (Estradiol kit, DRG Products, DRG instruments GmbH, Germany), according to the manufacturer’s instructions. Row estradiol concentrations were normalized to control values and ranged from 7.7 to 159.8 pg/mL. These data represent 13 independent cell culture batches, with each treatment measured once.

Progesterone concentration in 48h CM was determined by ELISA (Canepa et al., 2008) using 10 μL of diluted (1/5) sample. Progesterone concentrations were expressed as the ratio of progesterone (ng/mL) in treated condition normalized to that of control condition and ranged from 1.6 to 28.9 ng/mL. Data represent 21 cell culture batches, with each treatment measured once.

### 2.4. EVs isolation

EVs were isolated by differential ultracentrifugation, a common easy operating method for extracting particles based on their size and density differences but with medium purity (Yu et al., 2025). First, follicular fluids (4 mL) or 48-hour cell culture-conditioned media (48h CM, 12mL, pooled from all 12 treated wells for one condition) were centrifuged for 15 min at 3000g (Centrifuge 5702, Eppendorf, Germany). The resulting supernatants were centrifuged at 12 000 g (Centrifuge SIGMA 2-16KL) at 4°C for 30 min. Follicular fluid supernatants were stored separately overnight at -80°C. Subsequently, the FF or 48h CM supernatants were transferred to ultracentrifugation tubes (Ultra-ClearTM, Beckman, USA) and centrifuged at 100 000 g at 4 °C for 1,5 h using 70.1 Ti rotor and Optima XE 90 centrifuge (Beckman, USA). The pellets were resuspended in 7 mL of PBS (#28374, BupHTM ThermoScientific, USA) with Trehalose 25 mM (T9531, Sigma Aldrich, USA), which was pre-filtered through 0.1µm filter (Millex® Millipore, Merck), and this suspension was centrifuged again at 100 000 g at 4°C for 1.5h. The final EV-enriched pellets were resuspended in PBS-Trehalose: 130 to 350 µL for FF-EVs, and 60 to 130 µL for conditioned media EVs (48h CM EVs). From these suspensions, 100 µL of FF EVs and 50 µL 48h CM EVs were used for miRNA extraction with NucleoZOL (Macherey nagel, Germany) as described in following section. For particle characterization, 8 µL of FF EVs or 48h CM EVs suspensions were fixed with 8 µL of 2% glutaraldehyde in PBS. For the western blotting analysis, EVs pellets were resuspended in 50 µL PSB buffer containing 50mM TCEP (Macherey Nagel, Germany), and stored at -20°C for further analysis.

### 2.5. EVs particle size detection and concentration

#### 2.5.1. Nanoparticle Tracking Analysis

The size and concentration of the fixed FF EVs (n = 10) or 48h CM EVs (n = 10) were analyzed by Nanoparticle tracking assay (NTA, Zeta View PMX-120, Particle Metrix, Meerbusch, Germany) According manufacturer’s recommendations, fixed EVs suspension samples were diluted in pre-filtered PBS (filter 0.1µm) at a ratio of 1 :250 or 1 :500 for FF EVs and at 1 :4000 or 1 :8000 for 48h CM EVs. Preparations were injected in the sample chamber with sterile syringes. Measurements were performed at 25°C using 488 nm lazer on 11 positions with Nanoparticle Tracking Analysis ZetaView software (version 8.05.14 SP7). Concentrations were expressed as mean ± standard error of mean (SEM) particles/mL.

#### 2.5.2. Microscopic characterization

EV-enriched preparations were visualized using transmission electron microscopy. Microscopic imaging of EVs was performed by the Microscopy Department of the University of Tours on JEOL JEM-1011 instrument (Tokyo, Japan) equipped with a Gatan digital camera driven by Digital Micrograph software (Gatan, Pleasanton, CA) for image acquisition and analysis.

### 2.6. Western blotting

Protein concentration in 0h GC, 48h GC, FF EVs and CM EVs samples was assessed using the Protein Quantification Assay (Macherey Nagel, Germany) according to manufacturer’s instructions.

According MISEV’s recommendation (Welsh et al., 2024) EV samples were analyzed for tetraspanins CD63, CD81, and CD9, cytosolic proteins Alix and HSPA1, as well as for tubulin and secretory associated protein calnexin. Protein samples were incubated 5 min at 95°C. Protein lysates (10 μg FF EVs, 5 µg GC or 2 μg CM EVs were subjected to electrophoresis on 4–12 % acrylamide gel (Life technologies, Saint-Aubin,France) and transferred onto 0.2 μm mini Trans-Blot Turbo nitrocellulose membranes (Biorad, France). After blocking with 5 % non-fat dry milk powder (NFDMP) in TBS-0.1 % (BupH Tris Buffered Saline Packs, ThermoScientific, France) Tween-20 (TBST) for 90 min at room temperature, blots were incubated overnight with appropriate primary antibodies (for final dilutions, see Additional Table 1) in TBST with 5% NFDMP at 4°C The membranes were then washed in TBST and incubated with the appropriate secondary HRP-conjugated antibody.

### 2.7. MiRNA extraction

FF EVs, CM EVs or CG samples were collected as previously described in the text. Total RNA was extracted using NucleoZOL and filter columns from Nucleospin® miRNA kit (Macherey Nagel, Germany) according manufacturer’s instructions. The purity of RNA isolated from CG and EVs samples was evaluated using Nanodrop (Thermo Fisher) at the absorbance of 230 and 260 nm. Sample RNA purity, measured by 260/280 and 260/230 ratios is shown in Additional Table 2.

**Table 2.**
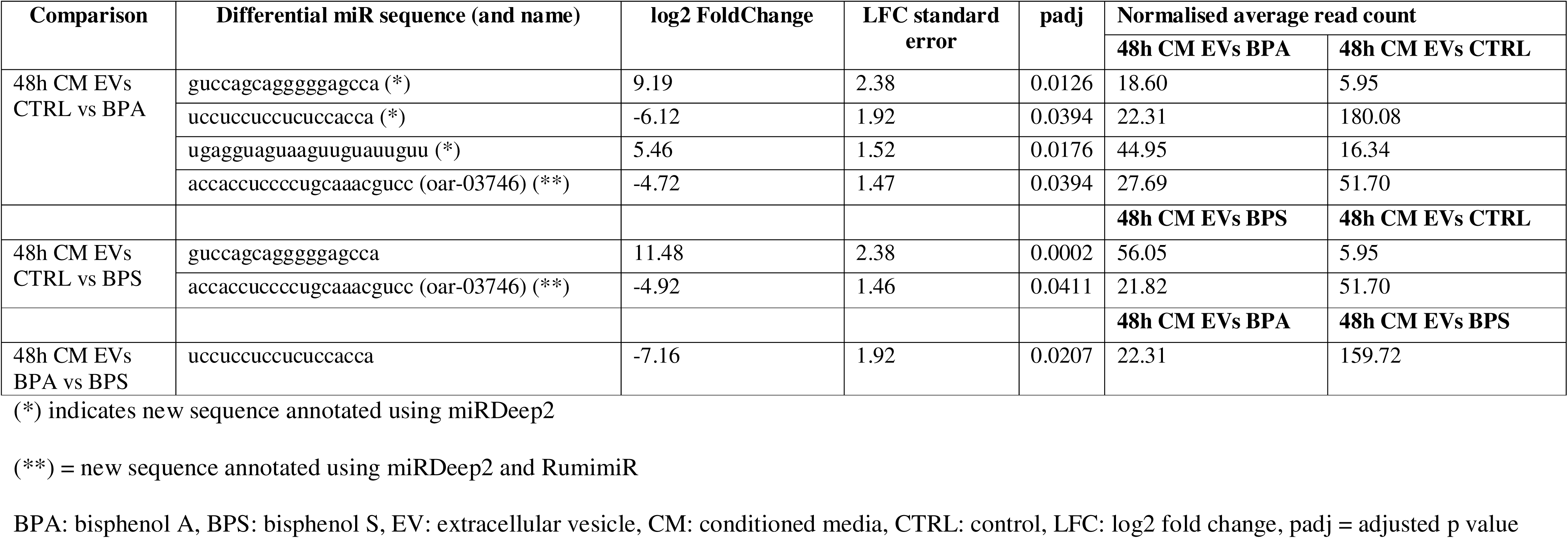
List of differentially expressed miRNAs in 48h CM EVs conditions. All p values were adjusted for multiple tests by the Benjamini and Hochberg method and had false discovery rates < 0.05.

### 2.8. Small RNAseq Gene Expression Studies

After verifying sample RNA purity using DropSense16 (Trinean) (Additional Table 2), next generation sequencing was performed at GeT Genopole Toulouse Midi-Pyrénées facility (GET, 2018). Eight condition groups were analyzed in 10 biological replicates each. For each of 80 samples, RNAseq libraries were constructed using 100 ng of total RNA for FF EV and CG samples or 4 μl of RNA preparations from MC-EV samples (approximately 80ng RNA) at the GeTLTRiX facility (GénoToul, Génopole Toulouse Midi-Pyrénées). NEXTFLEX® Small RNA-Seq Kit v4 with UDI was used according to manufacturer’s instructions. A bead cleanup was applied in place of the final sizing step. Briefly, 3’ and 5’ adapters were ligated to RNA fragments, converted to cDNAs and an amplification with nineteen cycles of PCR for FF and CG samples or with twenty-four cycles for CM samples was performed along with samples specific barcodes.

The yield and the quality of the libraries were assessed using D1000 ScreenTapes on TapeStation 4200 (Agilent Technologies, Santa Clara). The libraries were then pooled to equimolar concentrations and transferred to INRAE PGTB sequencing facility (PGTB, 2018). The library pool loaded into one P3 Flow Cell on Illumina NextSeq 2000 using 50 bp single read sequencing mode with NextSeq 1000/2000 P3 Reagent kit (50 Cycles). Quality control (QC) was performed on sequenced data both before and after mapping for all biological replicates. The raw sequence data for all replicates had a Q-score above 30. Sequencing data and experimental details are available in NCBI’s Gene Expression Omnibus (Edgar, 2002) and are accessible on European Nucleotide Archive database repository from EMBL’s European Bioinformatics Institute through accession number PRJEB109058 (https://www.ebi.ac.uk/ena/browser/view/PRJEB109058).

### 2.9. Bioinformatic treatments of sequencing data

Bioinformatic treatments of sequencing data were performed by the ISLANDe facility (INRAE Centre Val de Loire, UMR Physiologie de la Reproduction et des Comportements) using the computing resources of ISLANDe bioinformatics platform.

The miRNA sequences were analyzed using the nf-core/smrnaseq pipeline (version 2.4.0) (Alexander Peltzer et al., 2024). The read sequences were mapped using the ARS-UI_Ramb_v3.0 genome version (NCBI_Assembly: GCF_016772045.2). From the existing genome annotation, 104 precursor loci were included and complemented by two additional annotation sources using miRmachine (Umu et al., 2023) and using RumimiR database (Bourdon et al., 2019). FeatureCounts program from Subread v2.0.4 software (Liao et al., 2014) was used with strandness information to count the number of reads mapping to exons and summarized to the gene level to generate the count table for reads mapped on sheep genome, including information of transcripts using associated GCF_016772045.2 ARS-UI_Ramb_v3.0 gtf annotation file. The miRNA counting matrix from the pipeline was used directly in the differential analysis. A counting matrix was also created from the results of mirDeep2. This matrix only retained loci found in more than 10% of the samples.

### 2.10. Statistical analysis

All quantitative experiments were repeated with at least 10 independent biological replicates.

All statistical analyses were performed using R version 4.5 (R Core Team, 2025). For small RNAseq data, since amplification in 48h CM EV samples differed from the other conditions, standardization and analysis were carried out separately. Analyses were performed using the R/Bioconductor DESeq2 package (Love et al., 2014). Briefly, raw count table was filtered to eliminate undetected or poorly detected miRNAs (less than 10 median reads in all conditions). Normalization factors were calculated using RLE method. A negative binomial generalized log-linear model was fitted to count data. Pair-wise comparisons between biological conditions were extracted using specific contrasts to identify differential expression. A correction for multiple testing was applied using Benjamini-Hochberg procedure (BH) to control the False Discovery Rate (FDR) (Benjamini and Hochberg, 1995). miRNAs with FDR ≤ 0.05 were considered to be differentially expressed between conditions. The concentration of EV particles and the ratio according to EV size category were analyzed with a one-way analysis of variance (AnovaModel.1) and Tukey post hoc test (multcomp package). Steroid hormone concentration data were analyzed using non-parametric ANOVA by permutation (lmperm package), assessing the effect of treatment, cell culture batch, and their interactions. Pairwise comparisons between conditions were performed using Tukey post hoc test (nparcomp package).

### 2.11. MiRNA target gene function prediction

mRNA targets of each of miRNA sequences were predicted by the intaRNA-3.4.1 software with a minimum interaction length of 7-mers using 3’ untranslated regions (3’UTR) from GCA_016772045.2_ARS-UI_Ramb_v3.0 genome version. RNA-target duplexes were selected based on an interaction energy between -30 and -8 kcal/mol and 2-3 mismatches around the seed region. Descriptive functions were annotated using the biomaRt v2.62.1 package.

## 3. Results

### 3.1. Conditioned media progesterone and estradiol assays

Progesterone and estradiol mean concentrations ± standard error of mean in control spent media were 13,05 ± 1.96 ng/mL and 24.31 ± 4.39 pg/mL, respectively. After 48h of treatment, 10 µM BPA significantly decreased GC progesterone secretion compared to control (- 19%, p < 0.0001) whereas no effect was observed with 10 µM BPS (Figure 2A). Both 10 µM BPA and 10 µM BPS increased GC estradiol secretion by 511% (p < 0.0001) and 46%, respectively (p = 0.0027, Figure 2B).

**Figure 2.**
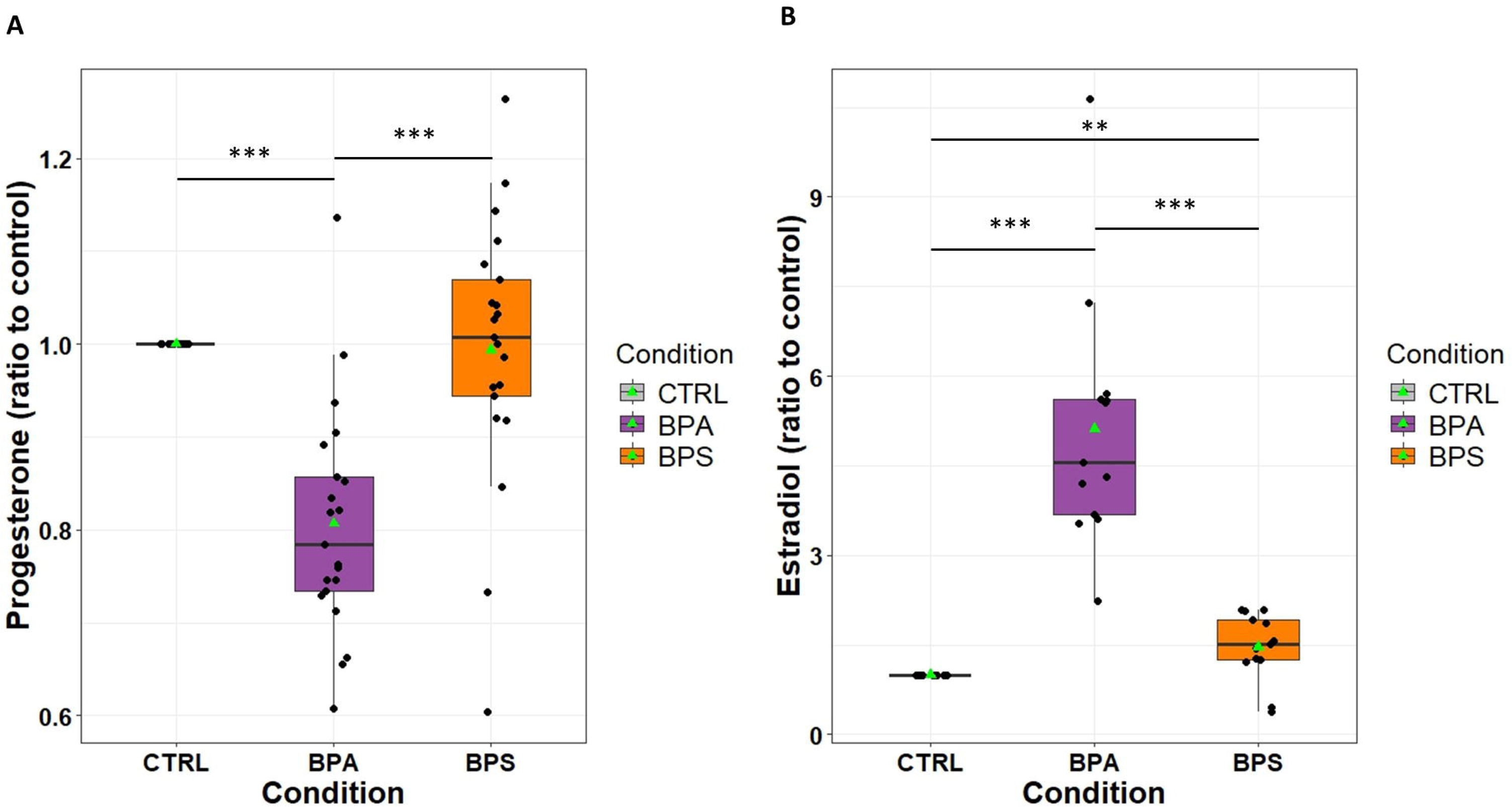
Effect of bisphenol A (BPA) and bisphenol (BPS) on ovine granulosa cells (GC) steroidogenesis. The progesterone (A) and estradiol (B) concentrations were measured in the culture medium after 48 hours. Cells were cultured in Modified McCoy’s 5A media in the presence or absence of 10 µM BPA or BPS at 10 ; the control condition (CTRL) contained 1/2000 DMSO. Results are from 21 independent cultures experiments, all batches were used for progesterone and 13 for estradiol assays. The mean concentrations in the control conditions were 13.05 ± 1.96 ng/mL for progesterone and 24.3 ± 4.4 pg/mL for estradiol. Hormone levels for each treatment condition were normalized to the control within the same experiment. The data are expressed as the mean ± SEM. Statistical significance was determined using non-parametric analysis of variance (ANOVA) by permutation (lmperm package). The treatment effect, culture effect and treatment by-culture interactions were assessed. Where a significant treatment effect was found, Tukey post hoc test (nparcomp package) was performed to determine differences between conditions (** p ≤ 0.01, *** p ≤ 0.001). Mean is indicated by green triangle and median is indicated with black bar in the box.

### 3.2. Characterization of ovine follicular fluid and 48h conditioned media EVs

The transmission electron microscopy results showed that both FF and CM samples contained EV like particles with some typical spherical or cup shaped structures (Figure 3A). Western blotting analysis confirmed that EV preparations were positive for specific makers including the tetraspanin proteins CD63 and CD81 and programmed cell death 6-interacting protein Alix (Figure 3B). Bands were also detected for proteins known to be promiscuously incorporated into EVs and non-vesicular extracellular particles (VNEPs), such as HSPA1A (both constitutive HSP73 and inducible HSP72 isoforms), which are also described in smaller nanoparticles like exomeres or supermeres (< 50 nm) (Yu et al., 2025). Additionally, a band corresponding to calnexin, an endoplasmic reticulum membrane protein, were detected in EV samples.

**Figure 3.**
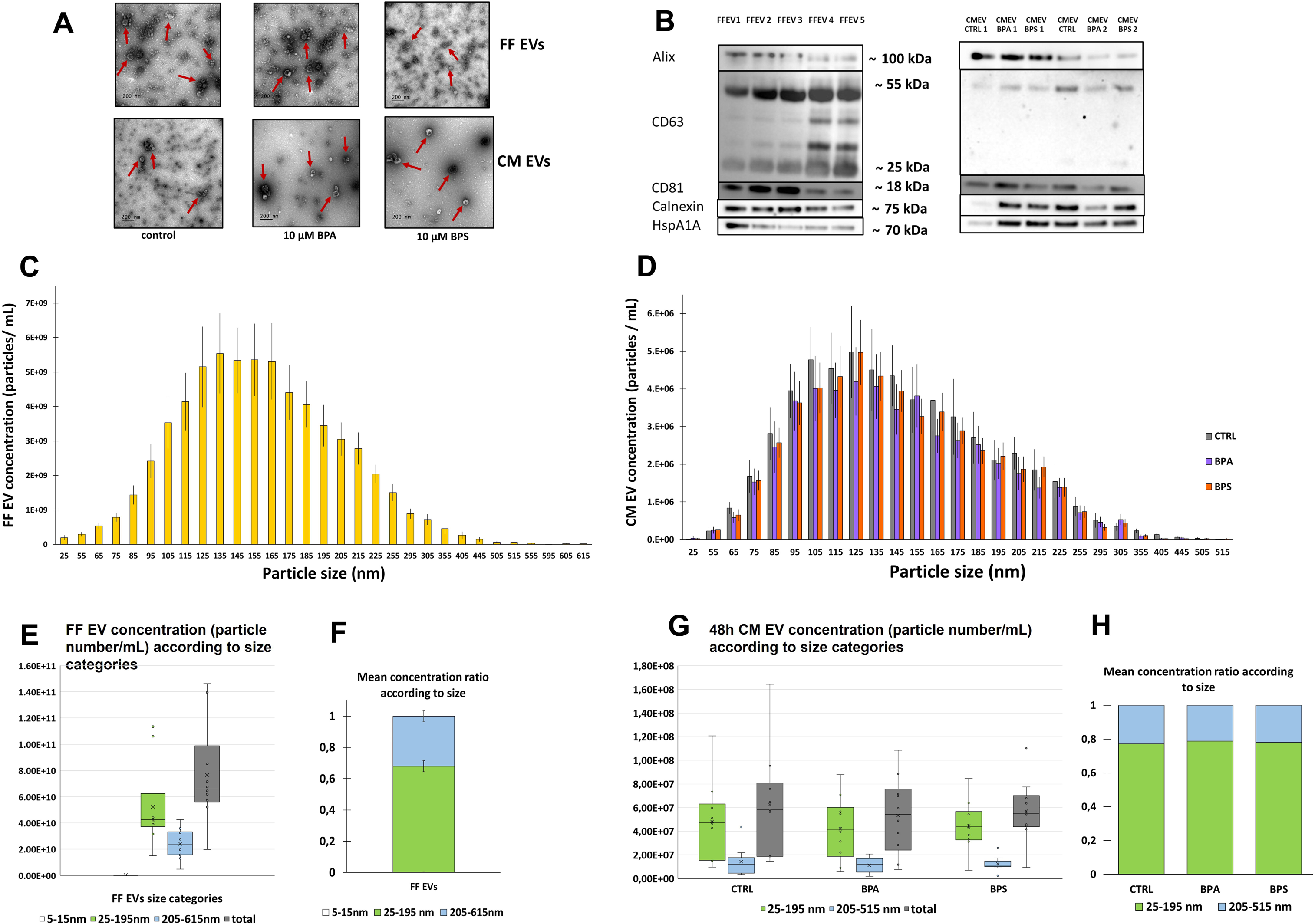
Morphological and molecular characterization of extracellular vesicles (EVs) in ovine follicular fluid and 48 h granulosa cell conditioned media (CM). A) Representative transmission electron microscopy images of EVs in FF and CM samples. EVs are indicated by arrows, scale bar = 200 nm. B) Immunoblotting analysis for EV specific protein marker (CD63, CD81, Alix) and non-vesicular extracellular particles (Calnexin, HSPA1A). Five FF EVs different samples (10 µg protein / lane) and six CM EVs samples (as 2 batches per condition, 2 µg protein / lane) were tested. No quantification was made. Results are presented as the mean ± SEM from 10 different experiments. Cand D) Representative of EVs size distribution assessed by nanoparticle tracking analysis in: FF (C) and 48h CM (D). E and G) EVs concentration according size categories in FF (E) and 48h CM (G). F and H) Mean EV concentration ratio according size categories in FF (F) and in 48h CM (H).

### 3.3. Nanoparticle tracking analysis

Particle size was measured in increments of 10 nm from 5 to 1005 nm. The average concentration of particles per mL of follicular fluid in FF EV samples (n = 10) is presented in Figure 3C. The total average concentration (±SEM) in FF EV samples was 76.5 x 10 ^9^ ± 12.3 x 10 ^9^ particles/mL of FF. Particles were divided into three size categories: 1) 5 to 15 nm, 2) 25 to 195 nm (small EVs) and 3) 205 to 615 nm (large EVs) (Figure 3E), given that for each increment size ≥ 625 nm, no FF EV samples or only few (≤ 2 samples) contained particles. These categories represented 0.1 %, 67.8 % and 32.1 % of the total particle count, respectively (Figure 3F). The peak particle size was 135 nm, with a mean concentration ± SEM of 5.54 x 10 ^9^ ± 1.16 x 10 ^9^ particles/mL of FF.

For conditioned medium EV samples (n = 10), the average concentration of EVs per mL of CM is presented in Figure 3D. The total average concentration of particles (±SEM) in CM control was 62.6 x 10 ^6^ ± 14. 08 x 10 ^6^/mL of CM. Given that, for each condition, few (≤ 2) or no samples contained particles ≤ 15 nm or ≥ 525 nm, particles were divided into two categories: 1) 25 to 195 nm (small EVs), and 2) 205 to 515 nm (large EVs) (Figure 4G). Under control condition, these categories represented 77.5 % and 22.5 % of the total, respectively (Figure 4H). Neither BPA nor BPS had a significant effect on the total particle concentration (Table 2) or on the proportional distribution of particles between these two size categories (Figure 4H). The peak size was 125 nm, with a mean concentration ± SEM of 4.98 x 10 ^6^ ± 1.22 x 10 ^6^ particles/mL of CM under control condition.

**Figure 4.**
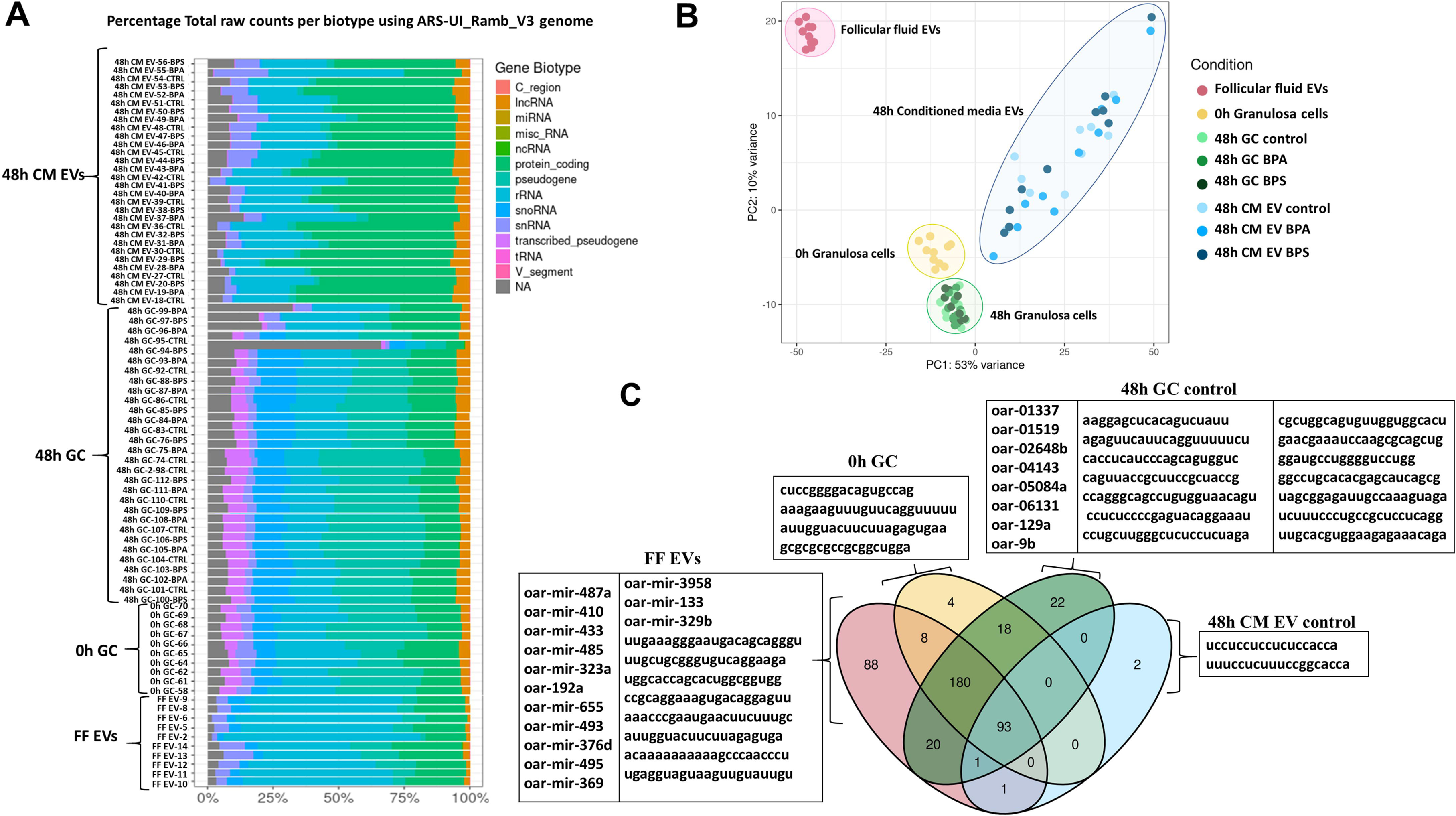
Characterization and identification of miRNA sequences in FF EVs, 0h GC, 48h GC, and 48h CM EVs conditions. A) Percentage distribution of total read counts by biotype using the ARS-UI_Ramb_V3 genome annotation versus the completed ovine genome annotation across all 80 samples. B) Principal component analysis (PCA) computed on all miRNAs. PCA was performed on a final set of 445 sequences. For the PCA, normalized counts were obtained using DESeq, which included the estimation of size factors and dispersions. A variance-stabilizing transformation was subsequently applied to the data for visualization. A miRNA sequence was retained for analysis if the median of its reads in at least one condition was greater than 10. For the FF EV category, the 22 miRNA sequences with highest median count are presented. FF = Follicular fluid, EVs = Extracellular vesicles, GC = Granulosa cells, CM = Conditioned media C) Venn diagram showing unique and shared differentially expressed miRNAs detected before normalization across four conditions: FF EVs (n = 10), 0h GC (n = 10), 48h GC CTRL (n = 10) and 48h CM EVs CTRL (n = 10).

### 3.4. Extension of the miRNA annotation in Ovis aries

Two complementary approaches were used to extend the miRNA annotation in Ovis aries. The first approach relied on miRMachine, which enables the identification of conserved miRNA orthologs previously reported in mammals through sequence homology searches. This strategy allows the detection of miRNA loci present in Ovis aries but not included in the original genome annotation. A total of 114 candidate precursors identified by miRMachine and absent from the original annotation were retained, identified with the prefix MIRMA and designated below as homology identified miRNA sequences. The second approach combined the results obtained from miRDeep2, which predicts putative precursor loci based on small RNA sequencing data, with ovine information from the RumimiR database. Only miRDeep2 candidate precursors producing identical mature miRNAs to those reported in sheep in the RumimiR database were retained, identified with the prefix RUM and designated below as ovine already reported miRNA sequences. By combining these two strategies, a total of 303 additional miRNA precursor loci were identified, extending the 104 precursors already included from the official genome annotation. This full set of annotated miRNAs was then used as input for the nf-core/smrnaseq pipeline. In addition, selection of miRDeep2 predicted miRNA precursors present in at least nine independent samples (>10% of the total samples) was considered to ensure robustness. These additional miRNAs were not incorporated into the reference genome annotation but were included exclusively in the downstream differential expression analyses.

### 3.5. Sequence quality and mapping of small RNAseq data

First, biotype-counting profiles (BCPs) for the different sample matrices were generated using the RambV3.0 ovine genome annotation (ARS-UI_Ramb_v3.0 (NCBI_Assembly: GCF_016772045.2) (Figure 4A). FF EV samples contained mainly rRNA, followed by protein-coding RNA, then snoRNA, snRNA, and lncRNA. GC samples showed a more balanced representation of each biotype, while CM EV consisted largely of protein-coding RNA, then rRNA, snRNA, and lncRNA. Very few reads were mapped to miRNA, but it can be explained by the limited number of annotated miRNA (104) in the chosen Ramb_v3.0 annotation. While this representation provided useful additional information on other non-coding RNAs (snoRNA, snRNA, lncRNA), it was not suitable for miRNA analyses.

Using this completed genome annotation version, biotype-counting profiles confirmed differences between the sample matrices (Additional Figure 2). The profiles indicated that in FF EV samples, miRNA represented 38.5% of all biotype counts (Additional Table 3; for detailed miRTrace biotype counts per sample, see Additional Table 4). In all 48h GC samples, miRNA represented about 22% of biotype count – a higher proportion than in 0h GC samples (14.5%). In all CM EV samples, miRNAs represented an average of only 0.30% of the total counts. Unknown RNA category comprising snoRNA, snRNA, and lncRNA, represented 99% of the total biotype counts in CM EVs while it represented only 61% counts in FF EVs and about 67% in GC samples. The read length distribution (Additional Figure 3) showed an enrichment of sequences 20-23 base pair (bp) long in FF EVs and GC samples, consistent with the expected size of mature miRNAs. In GC and CM EVs samples, the majority of sequences were up to 48 bp in length, while there is another peak around 20 bp. Following this, the sequencing data were mapped against the completed annotated reference genome. Small RNA sequencing generated an average of 14,900,394 ± 377,793 reads per sample for FF EVs, 18,346,728 ± 631,718 reads per sample for 0h GC, 16,069,898 ± 891,891 reads for 48h GC samples, and 14,828,006 ± 494,797 for CM EV samples (Additional Table 5). The average filtered read rate (mean ± SEM) was 81 ± 0.9 % for FF EVs, 90 ± 1 % for 0h GC samples, 93 ± 0.3 % for 48h GC, and 66 ± 1.7% for CM EVs samples. Mapping rates are detailed in Additional Table 6.

In total, our current study identified 533 miRNA sequences. Among these, 103 (19.3 %) sequences were exact matches of known ovine miRNAs. The 430 other sequences (80.7 %) were located outside of any annotated feature, among which, 301 (99.3 %) of the 303 identified additional miRNA precursor loci. Indeed, 112 (22.3 %) and 189 (35.5 %) were respectively mapped by homology using miRmachine, or miRDeep2 associated with RumimiR database. Finally, 129 (24.2%) were mapped using miRDeep2 predicted miRNA precursors.

### 3.6. Global detection of ovine miRNAs

Following filtering (retaining miRNA sequence with a median read count > 10 in at least one condition) and normalization, a total of 445 miRNA and all samples from the 8 groups were used for the principal component analysis (PCA) (Figure 4B). The PCA showed clear separation between the four sample matrices: FF EVs, 0h GC 48h GC and 48h CM EVs with a discrimination between 48h CM EVs and GC on component 1. Component 2 discriminated the FF EVs and the GC and also showed a large dispersion of 48h CM EVs samples. No distinct clustering based on BPA or BPS treatment was observed within 48h GC and 48h CM EVs groups.

Sequencing results showed 437 miRNA sequences across the four following conditions: FF EVs, 0h GC, 48h GC control, and CM EVs control. Overall, a total of 93 miRNA (21.3%) were identified commonly expressed among the four following groups FF EVs, 0h GC, 48h GC control and 48h CM EV control while 88 (20.1%), 4 (0.91%), 22 (5.0%), and 2 (0.46%) miRNA were exclusively expressed by these groups, respectively (Figure 4C).

### 3.7. Differences between ovine FF EVs and 0h GC miRNAs

After filtering, a total of 413 miRNA sequences were detected and analyzed in FF EVs and 0h GC samples, including 281 common sequences, 110 miRNAs specific to FF EV, and 22 miRNAs specific to 0h GC (Figure 4C). Differential expression analysis (DEA) revealed that 312 miRNA sequences were significantly differently expressed between FF EVs and 0h GC, including 79 known ovine miRNAs, 113 reported ovine sequences, 80 homology identified sequences, and 40 unidentified miRNA sequences. Of the total differential expressed (DE) sequences, 129 were upregulated in 0h GC (Additional Table 7), with 65 sequences exhibiting a log2 fold change (LFC) value between 2 and 38.05 (Additional Table 8). Particularly, 31 sequences were preferentially expressed by 0h GC (mean read < 10 in FF EVs), including 7 unknown predicted sequences (LFC > 6,5 and mean read = 0 in FF EVs), with « GCGCGCGCCGCGGCUGGA » (highest LFC = 38.05), the following unknown sequence « UGAGGUAGUAAGUUGUAUUGUU » (LFC = 16.64) and known ovine miRNAs such as miR-181a-1, miR-150, miR107, miR-103, miR-374, miR362, miR-21, miR-23b, miR-181d, miR-10b, miR-146, miR-26, and members of let-7 family.

On the other hand, 183 miRNAs were upregulated in FF EVs of the DE miRNAs, including 27 predicted miRNAs (Additional Table 6) and 76 miRNAs with a LFC ranging from - 2.16 to - 44.48 (Additional Table 9), with the following sequence « CAGGAUUAACCAGAGGACAGUGU » which had the highest LFC (= - 44.48). Of note, 51 sequences were preferentially expressed by FF EVs (mean read < 10 in 0h GC and > 10 in FF EVs), including 14 unidentified sequences predicted with miRDeep2 (LFC > - 4.55 and mean read < 10 in 0h GC). Key miRNA in this set also included known ovine miRNAs such as miR-376 family members, miR-410, miR-329, miR-485, miR-379, miR-655, miR-382, miR1185, miR-495, miR-143, miR-380, miR-494, miR-22, and miR-487b.

### 3.8. Differences between ovine 0h GC and 48h GC control miRNAs

After filtering, a total of 346 miRNA sequences in the 0h GC and 48h GC were counted and kept for analysis, including 291 common sequences, 12 0h GC specific miRNAs and 43 specific to 48h GC control (Figure 4C). DEA revealed 220 miRNAs that were differentially expressed between 0h GC and 48h GC control. These comprised 36 known ovine miRNAs, 157 homologous miRNA sequences and 27 newly identified sequences (Additional Table 7). Among the 111 miRNAs upregulated in 0h GC, 20 showed the most significant changes (LFC < - 2), including 2 known ovine miRNAs (oar-mir-30c and oar-mir-19b), 4 reported ovine sequences, 6 homology identified miRNA sequences and 8 unannotated predicted sequences. In this last group, the sequence « GCGCGCGCCGCGGCUGGA » is the most differential (LFC = - 12.87) (Additional Table 10). Of the 109 miRNAs upregulated in 48h GC control, 29 were the most significant (LFC > 2), including 2 known ovine miRNAs (oar-mir-27a and oar-mir-221), 6 ovine reported miRNA sequences, 8 homologous miRNA sequences and 12 unidentified sequences, including the most differential as « AGAGUUCAUUCAGGUUUUUCU » (LFC = 8.18) (Additional Table 11).

### 3.9. Qualitative comparisons between FF EV or 48h GC control with 48h CM EV control miRNAs

Due to differences in amplification efficiency, 48h CM EV samples were analyzed qualitatively rather than quantitatively. Comparisons were made between (i) 48h GC control and 48h CM EVs control, and (ii) FF EV and 48h CM EV conditions. First, of 336 miRNA sequences detected, 2 novel sequences were only identified in 48h CM EV control (UCCUCCUCCUCUCCACCA and UUUCCUCUUUCCGGCACCA, Figure 4C), and 245 sequences specific to 48h GC control (including 26 known annotated ovine miRNAs, 115 ovine reported sequences, 67 homologous miRNAs and 37 newly identified miRNAs). Eighty-nine sequences were common to both conditions (32 known ovine miRNAs, 25 ovine reported sequences, 28 homologous miRNAs and 4 unknown miRNAs).

From the second comparison, 394 miRNA sequences were detected in FF EV and 48h CM EV conditions. Of them, three novel sequences were specifically identified in 48h CM EV control, 303 sequences were only present in FF, and 88 miRNAs were common to both. These 88 common sequences represented 97% of miRNAs detected in 48h CM EV, 22.5% of those in FF EV and 26.4% of those in 48h GC.

### 3.10. Effect of 10 µM BPA or 10 µM BPS on 48h GC miRNAs expression

After filtering, 354 miRNA sequences across the 48h GC control, BPA-treated and BPS-treated GC samples were kept for analysis, with 313 sequences common to all three conditions. As shown in Figure 5A, 7, 6 and 9 miRNAs were specifically detected in 48h GC control, BPA or BPS conditions, respectively. DEA showed no differentially expressed miRNA between the 48h GC control and BPA-treated 48h GC samples. However, one miRNA identified by homology (oar-24b) was downregulated by 10 µM BPS compared to the control (LFC = - 0.277 ± 0.071, padj = 0.033). Moreover, another ovine already reported miRNA sequence (oar-01635) was upregulated in 48h BPA-treated GC compared to BPS-treated (LFC = 0.222 ± 0.046 and padj = 0.00063) (Figure 6A).

**Figure 5.**
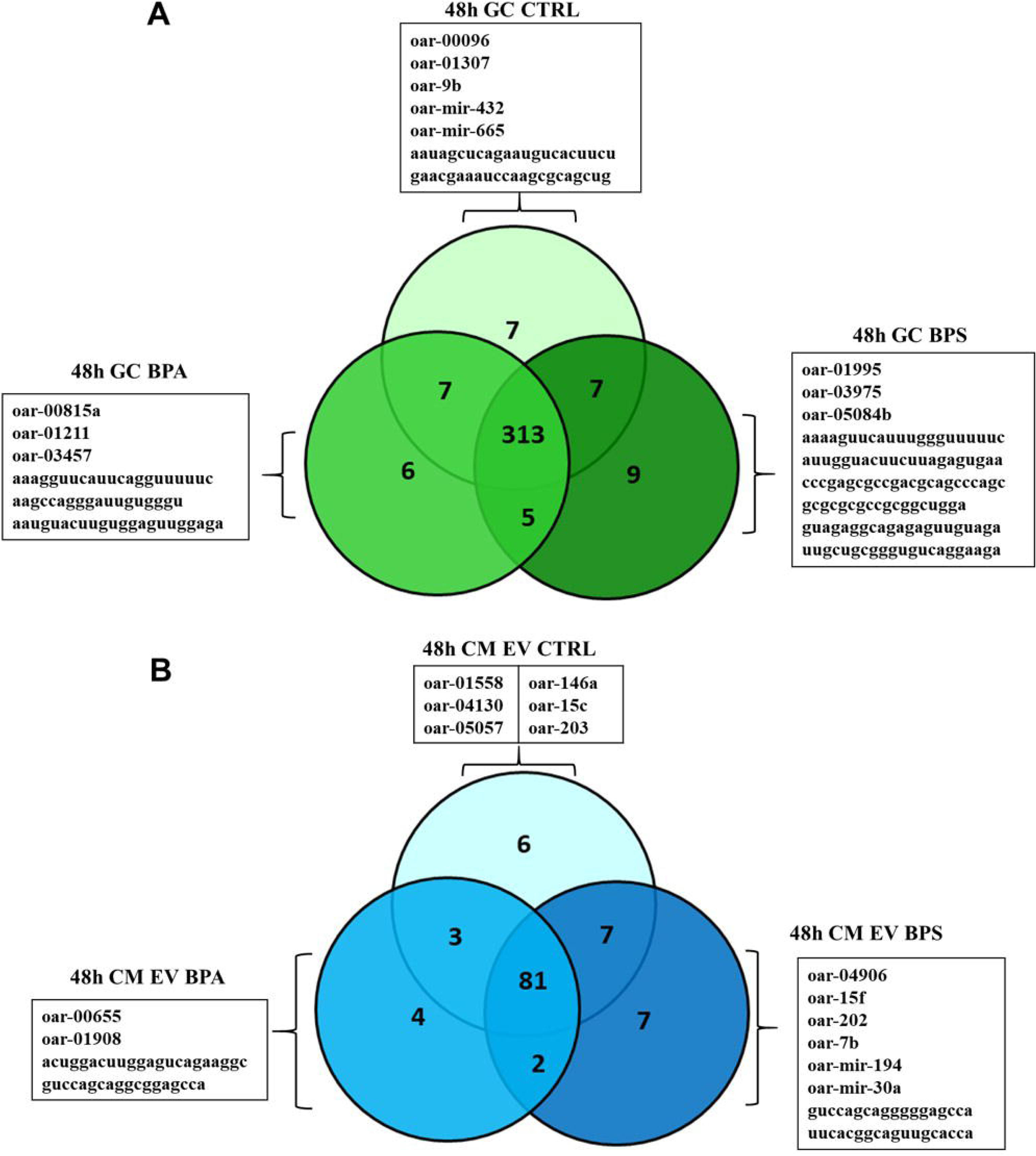
Venn diagram showing unique and shared detection of miRNAs in 48h GC and 48h CM EVs condition groups. A) This diagram shows miRNAs detected in 48h GC control, 48h GC BPA and 48h GC BPS conditions, after filtering and before normalization (n= 10: condition). B) This diagram shows miRNAs detected in 48h CM EVs CTRL, 48h CM EVs BPA and 48h CM EVs BPS conditions, after filtering and before normalization (n= 10: condition). CTRL = control A miRNA sequence was retained if the median of its reads in at least one condition must was greater than 10.

**Figure 6.**
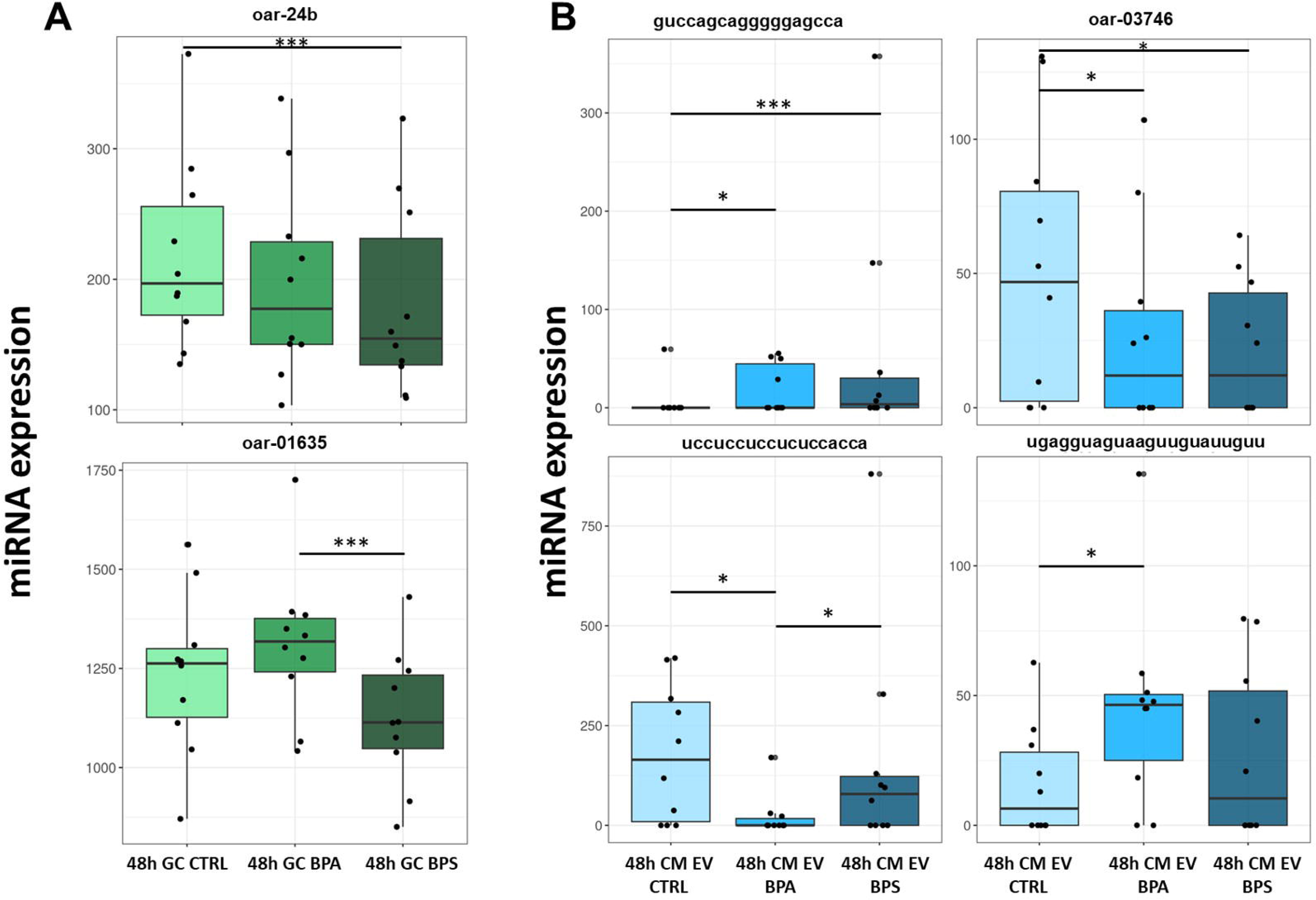
Differential analysis of candidate miRNAs in 48h GC and 48h CM EVs condition groups. (A) Expression of Oar-24b and oar 01635 in 48-hour GC in the presence of absence of 10 µM BPA or BPS compared to control condition. *** indicates adjusted p-value ≤ 0.001. (C) Expression of “guccagcagggggagcca”, Oar-03746, “uccuccuccucuccacca” and “ugagguaguaaguuguauuguu” in EVs from 48-hour conditioned medium (48h CM EVs) with or without 10µM BPA or BPS. *** indicates adjusted p-value ≤ 0.001, * indicates adjusted p-value ≤ 0.01

### 3.11. Effect of 10 µM BPA or 10 µM BPS on 48h CM EVs miRNAs expression

Following filtering, a total of 110 miRNA sequences in 48h CM EVs control, in 48h CM EVs BPA, and in 48h CM EVs BPS conditions. These comprised 35 annotated ovine miRNAs, 25 already reported ovine sequences, 34 homologous miRNAs, and 10 novel sequences. Specific condition expressed miRNAs were also identified: 6 in 48h CM EVs control, 4 in 48h CM EVs BPA and 7 miRNAs in 48h CM EVs BPS condition (Figure 5B).

DEA identified four differentially expressed miRNAs between 48h CM EVs control and 48h CM EVs BPA, including 2 upregulated (« GUCCAGCAGGGGGAGCCA » and « UGAGGUAGUAAGUUGUAUUGUU ») and 2 downregulated miRNAs (« UCCUCCUCCUCUCCACCA » and « ACCACCUCCCCUGCAAACGUCC » named oar-03746).

Among them, three are newly identified miRNA sequences, and one (oar-03746) was identified as a previously reported ovine sequence (Table 2). Of these four miRNAs, two were also differentially regulated in the BPS-treated condition versus the CM EV control, and another one was downregulated in the BPA condition compared the BPS condition (Figure 6B).

### 3.12. MiRNA target gene prediction

Target gene prediction was assessed for each of the 6 differentially expressed miRNA sequences. Between 27,957 and 37,416 targets per miRNA were detected (Additional Table 11). Between 111,029 and 190,537 alignments per miRNA sequence were made as each miRNA can align to several regions of the same target. As some GO Terms were missing, the function of targeted mRNA was replaced by the name of the protein generated by targeted mRNA. The top 30 most abundant predicted targeted protein and the specific targeted proteins for each differential miRNA are presented in Additional Figure 4 and Additional Figure 5, respectively.

## 4. Discussion

Small EVs and miRNA are known to play a crucial role in intercellular communication between ovarian follicular cells (Fazeli et al., 2025; Smith and Russell, 2022) and in follicle development (Alexandri et al., 2020) respectively. In that context, our study aimed to highlight the endocrine disruptor BPA and BPS mechanisms of action on ovine GC especially through EVs and their miRNA content. A further objective was to decipher the potential role of miRNAs in ovine follicle development, as we investigated miRNA profiles in FF EVs, granulosa cells and their conditioned EVs (CM EVs). First, we confirmed BPA and BPS both disrupt ovine GC steroidogenesis after 48h in vitro exposure. We also showed that EV size distribution was similar between FF EVs and 48h CM EVs. No effect of BPA or BPS was observed on 48h CM EV size distribution and concentration. In total, our current study identified 533 miRNA sequences including 129 newly identified sequences. In GC, 354 microRNAs were detected while 110 identified from their EVs. No change in GC miRNA expression was observed following 10 µM BPA exposure for 48h, while 10 µM BPS exposure decreased significantly a cellular miRNA named oar-24b. The expression of 2 CM EV associated miRNAs was altered by both BPA and BPS exposure, BPA also altering 2 additional CM EV associated miRNAs. These modifications could potentially be implicated in the regulation of BPA or BPS steroidogenic effect or in folliculogenesis.

### 4.1. MiRNA expression analysis in different matrices

Our study is the first to examine in ovine species, the differential abundance of miRNAs in FF EVs, granulosa cells and their corresponding CM EVs. In total, 533 miRNA sequences were identified using small RNAseq approach. Among them 103 sequences were exact matches of the 104 unique ovine annotated miRNAs, therefore 99 % of annotated ovine miRNAs were identified in our study. Another 301 sequences represented 99.3 % of the 303 identified additional miRNA precursor loci among reported ovine miRNA sequences or homology identified. Interestingly, 129 sequences of the 533 miRNAs (24.2%) were predicted and considered as novel ovine miRNAs, contributing to significant enrichment in ovine miRNA identification. Our study thus provided new miRNA candidates that could be relevant with the follicular development process and potentially exhibiting effects on oocyte, cumulus cells or early embryo development. MiRNAs lead to negative regulation of gene expression by base-pairing with complementary mRNA sequences, usually at the 3′UTR of mRNAs. Interaction of miRNAs with their targets may inhibit translation and/or induce mRNA degradation (Brieño-Enríquez et al., 2015). The high level of conservation of each functional miRNA across species points out how important miRNAs are as ancient components of genetic regulation (Bentwich et al., 2005; Griffiths-Jones et al., 2007).

Ovine miRNA depth annotation is less comprehensive than in human, rodent, or bovine (1,064 miRNA sequences annotated in bovine species). Moreover, next generation sequencing associated with bioinformatic analysis can predict hundreds of new miRNAs but most of them are not annotated in international databases yet. Also, in sheep, fewer tissues, such as heart, muscle, liver, intestine, blood, mammary gland, ovaries (Galio et al., 2013; Hou et al., 2018; Laganà et al., 2015; Puttabyatappa et al., 2022; Song et al., 2021) and fewer conditions have been explored until now and many miRNAs could be specific to a tissue, and could escape identification methods based on simple inter-species comparison.

### 4.2. Ovine follicular fluid as a rich source of EV-associated miRNAs

FF EVs and concomitant GC (0h) shared 68 % of the analyzed sequences after filtering. This corroborates the data reported in bovine (Andrade et al., 2017), or in human (Rooda et al., 2020) suggesting that FF EV miRNA cargo is qualitatively similar to those of GC, and that FF EVs are likely mostly originated from granulosa or cumulus cells rather than from blood supply. Moreover, this could support the hypothesis that their differential expression could matter in EV mediated cell communication in the follicular environment.

Qualitatively, 88 (97%) of the miRNA sequences detected in 48h CM EV control, corresponded to 22.5% of those in FF EV and 26.4% of those in 48h GC control. These low percentages could be due to the small amount of RNA material collected from 48h CM EV samples which were isolated from 12 mL of media and required a higher amplification. Indeed, this observation agrees with what has been described in bovine GC medium (Andrade et al., 2017), as CM EVs were isolated only after 48h of GC secretion, whereas FF EVs are accumulating in FF, which serves as a reservoir during follicular development (Revelli et al., 2009).

Comparison of miRNA sequences within FF EVs versus 0h GC showed the greatest number of differentially expressed miRNAs, 75.5 % of the analyzed sequences are differentially expressed, some of which have already been described in ovarian and GC functions (Alexandri et al., 2020; Chen et al., 2026).

Known ovine miRNAs represented respectively 25.3 % of the differentially expressed sequences between FF EVs and corresponding 0h GC. Several known ovine miRNAs were more abundant in GC before culture (0h), such as miR-181a-1, miR150, miR-107, miR-21, miR-23b, miR-181d, miR-10b, miR-146, miR-26, and the members of let-7 family.

In 0h GC, the most upregulated sequence was the unknown one « GCGCGCGCCGCGGCUGGA » (LFC = 37 with corresponding FF EV mean read =0). Of note, the following homology identified oar-223 sequence, « GUGUCAGUUUGUCAAAUACCCC », was considered most differential and present in both conditions (LFC = 6.05 with corresponding 0h GC and FF EV mean read being 13,125 and 196, respectively). Interestingly, in buffaloes, a study reported an association between reduced level of three miRNAs including miR-223 in saliva and the presence of dominant ovarian follicles (Singh et al., 2017). Another study showed that miR-223 significantly inhibits both the steroid hormone synthesis and the proliferation of GCs in chicken (Xiang et al., 2024), highlighting its importance in regulating granulosa cell function.

Among the known miRNAs overabundant in 0h GC, miR-150, let-7 and miR-21 have been related to ovine GC apoptosis and proliferation by targeting DNA-binding protein inhibitor and steroidogenic acute regulatory genes (Song et al., 2023; Zhang et al., 2022; Zhou et al., 2019). In pig, inhibition of miR-21 during meiotic maturation of the oocyte alters Programmed Cell Death Protein 4 (PDCD4) abundance that could possibly impact subsequent embryonic development (Wright et al., 2016). MiR-107 was showed to suppress porcine GC proliferation and estradiol synthesis (Liu et al., 2025) and miR-181a was involved in GC apoptosis induction (Zhang et al., 2020). MiR-10b was related to inhibition of GC proliferation in goat through targeting Brain-derived neurotropic factor (Peng et al., 2016).

In FF EVs, the most upregulated sequence was « CAGGAUUAACCAGAGGACAGUGU » (LFC = - 44.5 with corresponding 0h GC mean read =0). The following sequence « GGUCCAGUUUUCCCAGGAAUCCC », named oar-145, was considered most differential and expressed in both conditions (LFC = -5.96 with corresponding 0h GC and FF EV mean read being 672 and 38293). MiR-145 was described to regulate steroidogenesis in mice primary GC and proliferation in KGN cell line (Lingling et al., 2024; Ma et al., 2023).

Among the known ovine upregulated miRNAs, miR-143 (LFC = -4.53) was reported to regulate mouse GC estradiol production and proliferation by targeting the Kirsten rat sarcoma 2 viral oncogen homolog (KRAS) (Zhang et al., 2017). Overexpression of this miRNA decreased both progesterone and estradiol secretion and promoted GC apoptosis in bovine GC (Z. Zhang et al., 2019). MiR-22 inhibits mouse granulosa cell apoptosis by targeting SIRT1 (Xiong et al., 2016)and in bovine follicular cells, miR-494 regulates PTEN levels (Andrade et al., 2017). Thus, our study identified novel miRNAs, some exhibiting huge variation of expression between follicular compartments. Further studies are required to understand their role in the follicular development.”

### 4.3. Alteration in granulosa cell miRNA expression following 48 h in vitro culture

Analysis revealed that 220 (63.6 %) miRNAs were differentially expressed in GC before (0h) and after 48h (GC control), suggesting that 48h in vitro cell culture deeply modified cellular miRNA regulation. Indeed, the GC removal from their native in vivo environment to in vitro culture environment could induce cell stress and activate factors such as miRNAs that would affect cell signaling and function (Yenuganti and Vanselow, 2017).

The most upregulated sequence in 0h GC compared to 48h GC control « GCGCGCGCCGCGGCUGGA » as well as the most upregulated sequence in 48h GC control compared to 0h GC « AGAGUUCAUUCAGGUUUUUCU » have not been described yet. MiR-27 and miR-221 were the ovine known most upregulated miRNAs in 48h GC control condition and could be involved in GC function in vitro. Indeed, exosomal miR-27, downregulated in FF of patients with ovarian hyperstimulation syndrome increase production of ROS and apoptosis in KGN cell line (Liu et al., 2021), whereas miR-221 has been described as a modulator of ovarian function and was related to inhibition of cattle GC steroidogenesis (Robinson et al., 2018). Some other differential miRNAs identified as oar-15f (mir-503), oar-202 (mir-202), oar-155 (mir-155) could be regulators of important functions related to oocyte maturation and female fecundity as reported in human, canine, fish species and KGN cell line (Cao et al., 2022; Dettleff et al., 2025; Gay et al., 2018; Xia and Zhao, 2020). Among the differential miRNA between in vivo and in vitro ovine GCs, effects of oar-miR-30c, oar-miR-19b and oar-miR-150 are not identified yet, and therefore further studies would be required to highlight their functional roles in GCs.

### 4.4. Impact of bisphenol exposure on cellular and EV-associated miRNAs in granulosa cells

Our study has investigated the association between 10 µM BPA or BPS exposure and EV-enriched miRNAs from ovine primary granulosa cells. In this study, miRNA expression in ovine GC did not change following 48h exposure to 10µM BPA. However, 10 µM BPS exposure significantly decreased the expression of a cellular miRNA identified by homology and named oar-24b, compared to control condition. This sequence is homologous to the mir-24b precursor (MiRBase 22.1), miR-24 have been implicated in follicular cell steroidogenesis and metabolism (Gong et al., 2024; Ma et al., 2024; Wei et al., 2024). To our knowledge, this study is the first to report changes in miRNA expression in GC following acute BPS exposure. Notably, compared to BPA, BPS decreased the expression of oar-01635 sequence, which is homologous to miR-874 in several species.

Our results regarding BPA are not in line with the literature, as BPA exposure altered miRNA expression in human breast cancer cell line MCF-7, ovarian cell lines, placenta, and GC (Tilghman et al., 2012; Li et al., 2014; Deng et al., 2021; Chen et al., 2023; Hui et al., 2018; Márton et al., 2023; De Felice et al., 2015; Rodosthenous et al., 2019) which could be related to cell migration and proliferation, cell growth differentiation and tumorigenesis. In bovine, in vitro BPA exposure also leads to miRNA expression variation in granulosa cells, cumulus cells and enclosed oocytes (Sabry et al., 2022, 2021). Discrepancies between literature and our results could be in part related to culture conditions such as BPA concentration, lower in our study compared to those used in human or bovine GC studies, cell or tissue species and exposure duration.

In CM EVs, differential analysis identified 4 miRNAs altered by BPA, and two altered by BPS compared to control. BPA decreased the expression of oar-03746 and « UCCUCCUCCUCUCCACCA » sequences and increased that of « GUCCAGCAGGGGGAGCCA » and « UGAGGUAGUAAGUUGUAUUGUU » novel sequences. BPS also decreased the expression of oar-03746 and « GUCCAGCAGGGGGAGCCA » sequence. Oar-03746 sequence identified using miRDeep2 and already reported in RumimiR database could be related to miR-1306-5p (MiRBase 22.1). In ovine GC, miR-1306 promoted cell apoptosis by reducing BMPR1B mRNA and protein levels (Abdurahman et al., 2022). BMPR1B is the major gene of litter size identified in sheep and a core member of Bone morphogenetic protein (BMP)/Smad signaling pathway (including BMP15 and Smad4) closely related to fecundity in sheep. In porcine GCs, miR-1306 was identified as a functional miRNA that targets TGFBR2 receptor, promoting apoptosis of GCs as well as attenuating the TGF-β/SMAD signaling pathway (Yang et al., 2019).

In vitro BPA exposure of human GC altered expression of 9 EV-associated miRNAs, increasing levels of three of them and decreasing levels of six of them including miR-27b-3p which level was also decreased in corresponding GC (Rodosthenous et al., 2019). In addition to the above mentionned differences such as cell species, culture conditions and bisphenol concentrations, differences in miRNA expression (amplification score, sample detection, criteria selection) could explain discrepancies between studies. In our study, BPA and BPS exposure was low and could explain why no differential miRNA was found in GC after BPA exposure and only a small number of differential miRNAs in CM EV after BPA or BPS exposure. Taking together, these results indicated that although BPA and BPS share structural similarity, their functional and molecular effects on GC miRNA expression may be different and would require further studies, i.e. after several concentration or duration of exposure, to highlight their underlying mechanisms of action.

### 4.5. Characterization of non-coding small RNA cargo in FF EVs, GC EVs and CM EVs

We performed RNA sequencing of short non-coding RNAs (ncRNAs) extracted from FF EVs, and also from GC and corresponding CM EVs after 48h treatment with BPA or BPS, using read length of 50 nucleotides. In our study, different counting profile for ncRNA biotypes were observed according to matrices. Thus, FF EV samples mainly contained rRNA, followed by protein coding RNA, snoRNA, snRNA and lncRNA. GC contained more balanced representations of each biotype, whereas 48h CM EV contained the lowest proportion of miRNAs with most of of protein coding RNA, rRNA, snRNA and lncRNA. In fact, ncRNAs are divided into housekeeping and regulatory ncRNAs, which regulate expression of coding RNA through transcriptional and post transcriptional control. Housekeeping ncRNAs are expressed constitutively and ubiquitously, and play crucial roles in routine cell maintenance; they include transfer (t)RNAs, ribosomal (r)RNAs, and small nuclear (sn)RNAs. Regulatory ncRNAs are expressed in specific cell types and functions, in response to developmental cues, internal conditions, and environmental stimuli. They include miRNAs, short-interfering (si)RNAs, PIWI-interacting (pi)RNAs, long noncoding (lnc)RNAs, natural antisense RNAs, and circular (circ)RNAs (Idda et al., 2018; Robles et al., 2019).

Although our present study was especially focused on miRNA expression, regulatory ncRNAs like lncRNAs, snoRNAs or snRNAs and circRNAs could also be relevant to our inquiry, particularly since some of them have been associated with bisphenol related human diseases (He et al., 2024). Expression of lncRNA, snoRNA and snRNA was also correlated with coding RNA in liver of prenatal BPA-exposed female sheep, whereas this was not observed regarding the modulated miRNAs (Puttabyatappa et al., 2022).

### 4.6. Extracellular vesicle characterization

Our study focused on the EV-related molecular mechanisms of granulosa cell response to BPS and BPA. In vivo, GC line the inner walls of the follicle and are bathed with follicular fluid containing cell secreted extracellular vesicles. Whether FF EVs originate from different follicular cells and from blood circulation too, they are likely different from the EVs secreted to medium by GC in vitro. Here, EVs were isolated from ovine FF (FF EVs) and from the spent condionned medium of ovine GC after 48h cell culture (CM EVs) as it has already been reported for FF from other species such as bovine, human and equine (Da Silveira et al., 2012; Navakanitworakul et al., 2016; Santonocito et al., 2014; X. Wang et al., 2024), or in bovine GC (Menjivar et al., 2023) or in human granulosa primary or cell lines (Li et al., 2024). EV concentration in ovine FF was similar to that in bovine (Hailay et al., 2019) or rhinoceros (Gad et al., 2024). NTA and electron microscopy indicated similarity of distribution size of small EVs (< 200 nm) isolated from FF or 48h conditioned media, with peak EV size of 135 nm and 125 nm in diameter, respectively, which correspond to small EVs.

Using sequential centrifugations and double ultracentifugation, we have obtained heterogeneous population of ovine FF EVs, that was enriched in small EVs, although a low percentage of larger EVs as microvesicles (> 200nM) were also detectable. In fact, small EVs and microvesicles, although differed in size, shared several molecular markers (Allelein et al., 2021; J. Wang et al., 2024; Willms et al., 2018). As expected, CM EVs from BPA-treated or BPS-treated GC showed no difference in particle distribution, as it was previously reported for BPA in human female placental explants after a 24h exposure (Sheller-Miller et al., 2020). In accordance with the NTA analysis, EV protein marker analysis revealed in both FF EVs and CM EVs the presence of CD63, Alix, and CD81 known to be particularly enriched in the small EVs (Welsh et al., 2024). In this study, detection of both constitutive HSP73 and inducible HSP72 isoforms of HSPA1A in EV samples could indicate a possible isolation of both EVs and non-vesicular extracellular particles (NVEPs) in our FF EV and CM EV preparations (Yu et al., 2025). Just like, the detection in EV samples of calnexin, an endoplasmic reticulum membrane protein (Welsh et al., 2024), could indicate the presence of early endosomes which can interact with these structures (Doyle and Wang, 2019). However, it should be noted that FF EVs are heterogeneous in size, cargo composition, and biogenesis pathway (Fazeli et al., 2025), especially, different subpopulations of FF EVs were identified in human (Neyroud et al., 2022) and bovine (Wang et al., 2021).

### 4.7. Strength and limitations of the study

Our study is the first to explore BPA and BPS action mechanisms through the characterization of cellular and EV-associated miRNAs from ovine granulosa cell and FF, using small RNA sequencing. The high number of replicates in each group were sufficient to ensure robustness of the results and avoiding false positive differential expressed miRNA. Our study provides a great number of novel miRNAs in the ovine species, which could be further studied regarding targeting GC functions, follicular development or ovine female fertility. In addition, several of these miRNAs could be promising candidates for assessing the impact of bisphenol exposure on female fertility.

The confirmed effects of BPA and BPS on ovine GC steroidogenesis and in our study validated the use of this model to further investigate BPA and BPS mechanisms of action especially through EVs and their miRNA content. BPA and BPS both disrupt ovine granulosa cell steroidogenesis after 48h in vitro exposure with partially different effect: 10 µM BPA decreased progesterone and increased estradiol ovine GC secretion, whereas 10 µM BPS only increased estradiol secretion which is in line with our previous published data (Téteau et al., 2020, 2023). Of note, these concentrations do not affect GC cell viability and proliferation (Téteau et al., 2020). Despite an important number of ovaries and cell batches used in our study, heterogeneity of age, health and metabolic status of slaughtered lambs and adult sheep can affect reproductive physiology and contribute to variability of GC steroidogenesis in response to bisphenols.

In vitro steroidogenesis disruption induced by BPA or BPS was also reported from several granulosa cell species such as porcine (Wu et al., 2018; Song et al., 2019; Bujnakova Mlynarcikova and Scsukova, 2021), bovine (Campen et al., 2018) human (Amar et al., 2020; Lebachelier De La Riviere et al., 2023) and rodent (Samardzija et al., 2018; Zhou et al., 2008). Discrepancies regarding the effects of BPA or BPS were also reported in some of these species. In conclusion, in our study, 10 µM BPA consistently reduced progesterone and increased estradiol secretion in ovine GC in vitro, whereas BPS had more variable stimulatory effect on estradiol alone. This variability may have influenced the effects of bisphenol exposure on miRNA expression and could explain the moderate effects observed. Moreover, the possibility of EV or miRNA degradation related to cellular metabolism after 48h of culture cannot be ruled out. It is possible that focusing on earlier time points such as 12 or 24h of bisphenol exposure, might have provided more pronounced changes in expression.

In this study, the observed changes in miRNA expression could not be directly associated with changes in the expression of their cellular target genes. Therefore, additional experiments are needed to decipher the regulatory mechanisms, identify the targets and define the functional roles of these differential miRNAs. Furthermore, since ultrafiltration was not performed and only centrifugations were used, the possibility of EV co-precipitation with other components that carry miRNAs in extracellular space cannot be excluded. Finally, because amplification in the 48 h CM EV samples differed from that in the other conditions, quantitative comparison between the different 48h GC conditions and their respective 48-hour culture medium EV conditions was not possible. This limitation deprived us from obtaining additional information about the regulation of cellular and EV-associated miRNAs by BPA and BPS. In addition, transcriptomic analysis of GC treated by bisphenols for 48 h may be useful to elucidate the targets of differentially expressed miRNAs reported in this study

## 5. Conclusion

In conclusion, this study identified 533 miRNAs in ovine FF EVs, GC and CM EVs, including 129 novel sequences of putative miRNA. Our data showed that acute 48h BPA or BPS exposure altered the expression of specific cellular and EV-associated miRNAs from primary ovine granulosa cells and culture media. This suggested that bisphenols can modulate intercellular communication via extracellular vesicles, and that BPA and BPS may act through distinct molecular mechanisms. Further studies are required to elucidate the biological roles of these differentially expressed cellular and EV-enriched miRNAs in the context of bisphenol exposure effects on female fertility.

## Availability of data and materials

The datasets generated and analysed during the current study are available in the European Nucleotide Archive repository from EMBL’s European Bioinformatics Institute through accession number PRJEB109058 (https://www.ebi.ac.uk/ena/browser/view/PRJEB109058).

## Acknowledgements

The authors would like to thank Albert Arnould and Benoît for ovine ovary collection and the Endocrinology and Phenotyping Laboratory (INRAE Centre Val de Loire, UMR PRC, INRAE Nouzilly) for progesterone assay.

RNA sequencing was performed at GeT Genopole Toulouse Midi-Pyrénées facility (https://doi.org/10.15454/1.5572370921303193E12). We are very thankful Claire Naylies and Gaëlle Payros for their contribution to the RNAseq libraries preparation at GeT-TRiX, and Zoé Compagnie and Prescillia Alves-Gomes for the sequencing of libraries at INRAE PGTB sequencing facility (doi:10.15454/1.5572396583599417E12). We are very grateful to the genotoul bioinformatics platform Toulouse Midi-Pyrenees (Bioinfo Genotoul) for providing computing resources for preliminary RNAseq bioinformatics and biostatistics analyses performed by Yannick Lippi (GeT-TRiX).

## Additional Figure legends

**Additional Figure 1. Immunoblotting analysis for EV specific protein marker (CD63, CD81, Alix) and non-vesicular extracellular particles (Calnexin, HSPA1A).**

Five FF EVs samples (10 µg protein / lane) and six CM EVs samples (as 2 batches per condition, 2 µg protein / lane) were tested. No quantification was made.

FF = Follicular fluid; CM = Conditioned Media; EVs = Extracellular vesicles; BPA = Bisphenol A; BPS = Bisphenol S; CTRL = Control

**Additional Figure 2. Percentage Total counts per biotype using ARS-UI_Ramb_V3 genome and completed annotated genome.**

BPA = Bisphenol A; BPS = Bisphenol S; CTRL = Control; FF = Follicular fluid; GC = Granulosa cells; CM = Conditioned media; EVs = Extracellular vesicles

**Additional Figure 3. Sequence Length distribution**

A) MirTrace sequence length distribution of all 80 samples. B) MirTrace Sequence length distribution according conditions, n= 10 per condition.

**Additional Figure 4. Top 30 of the differential miRNA most abundant functions**

**Additional Figure 5. Differential miRNA most specific functions**

## Additional Table legends

**Additional Table 1: Primary anti-bodies used for western blotting.**

**Additional Table 2: Purity of total RNA isolated from granulosa cell and extracellular vesicle samples.**

**Additional Table 3: Mean percentages of Total count per RNA biotype according to conditions.**

**Additional Table 4: MiRTrace count per RNA biotype in all 80 samples.**

**Additional Table 5: Filtering statistics of sampled reads**

**Additional Table 6: Mapping rates in samples**

**Additional Table 7: Distribution of upregulated miRNA sequences in FF, 0h GC, 48h GC according to their identification**

**Additional Table 8: List of upregulated miRNA sequences in 0h GC compared to FF EVs.**

**Additional Table 9: List of downregulated miRNA sequences in 0h GC compared to FF EVs.**

**Additional Table 10: List of downregulated miRNA in 48h GC compared to 0h GC.**

**Additional Table 11: List of upregulated miRNA in 48h GC compared to 0h GC.**

**Additional Table 12: Target gene IntaRNA prediction of the 6 differential miRNA sequences**

