## Supplementary figures and images for "Characterization of ovine follicular fluid and granulosa cell-derived extracellular vesicles and their miRNA cargo following in vitro exposure to bisphenols A and S"

### Supplemental Figure1

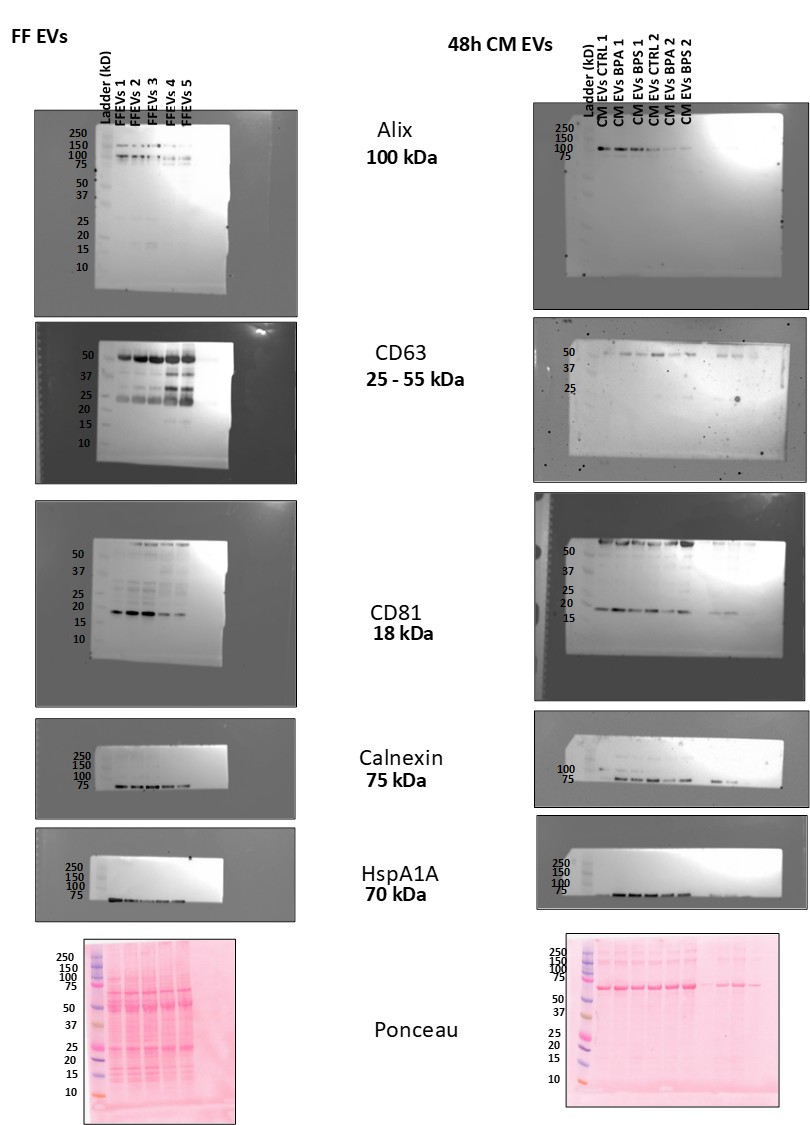

### Supplemental Figure2

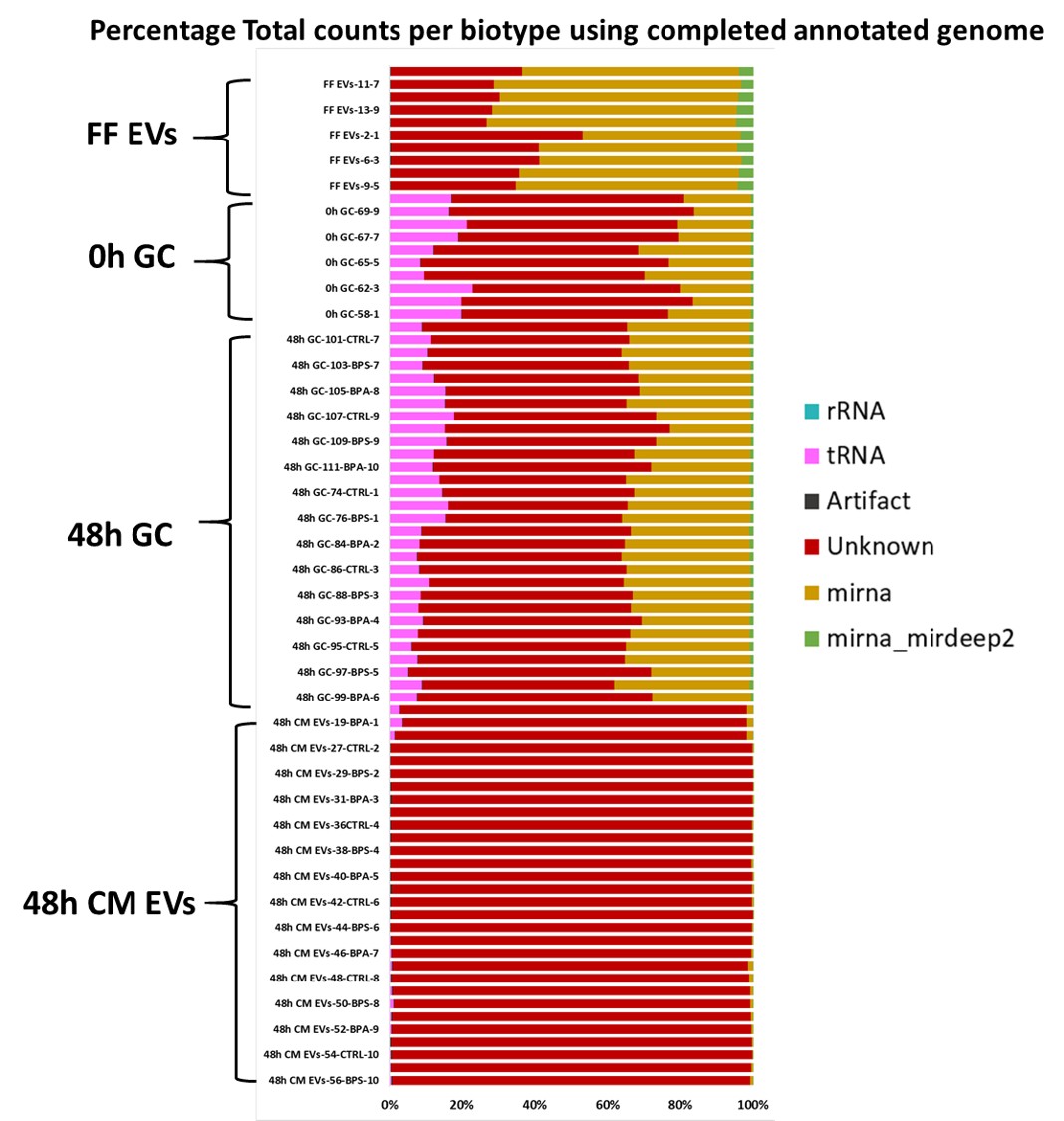

### Supplemental Figure3

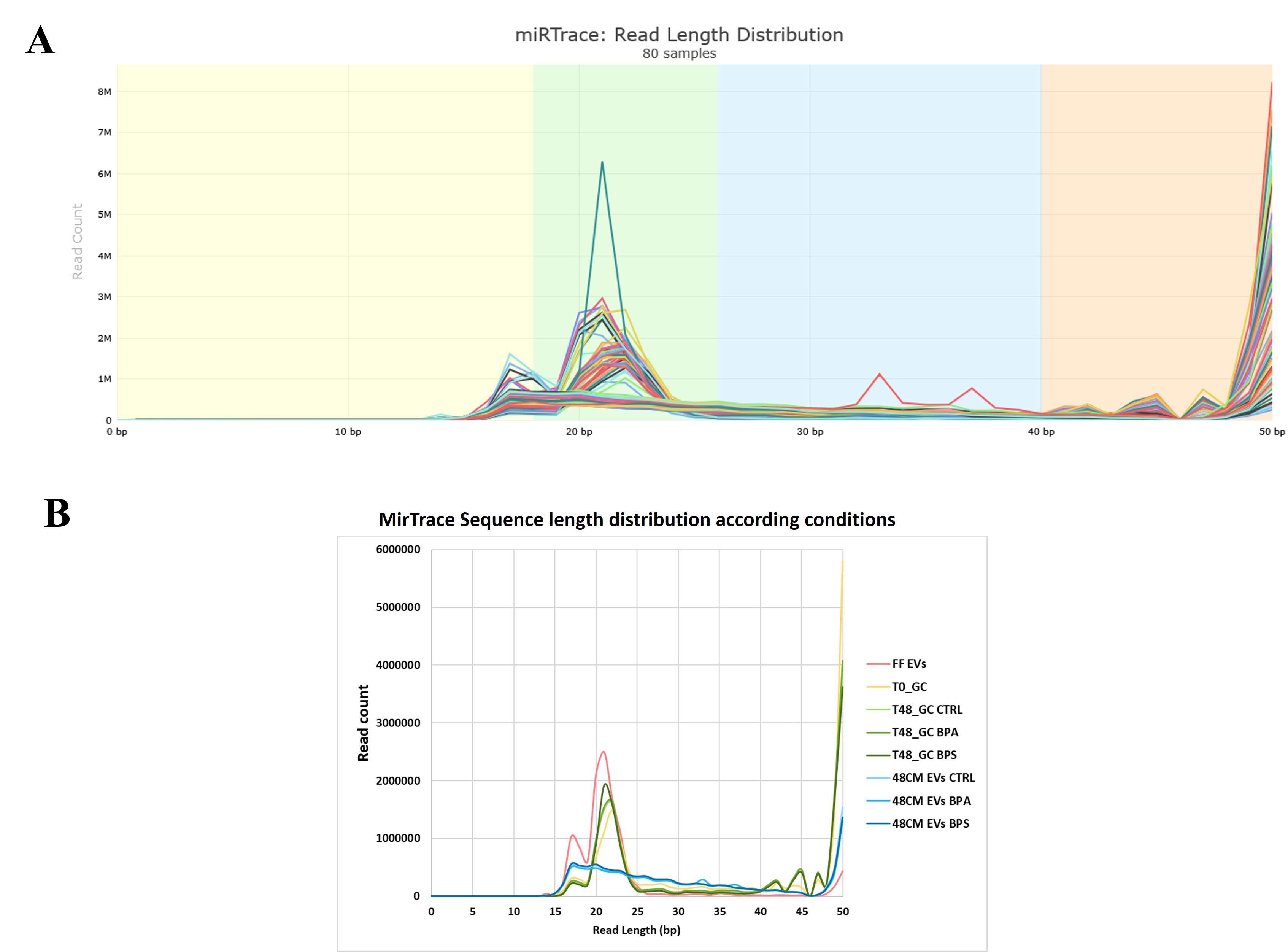

### Supplemental Figure4

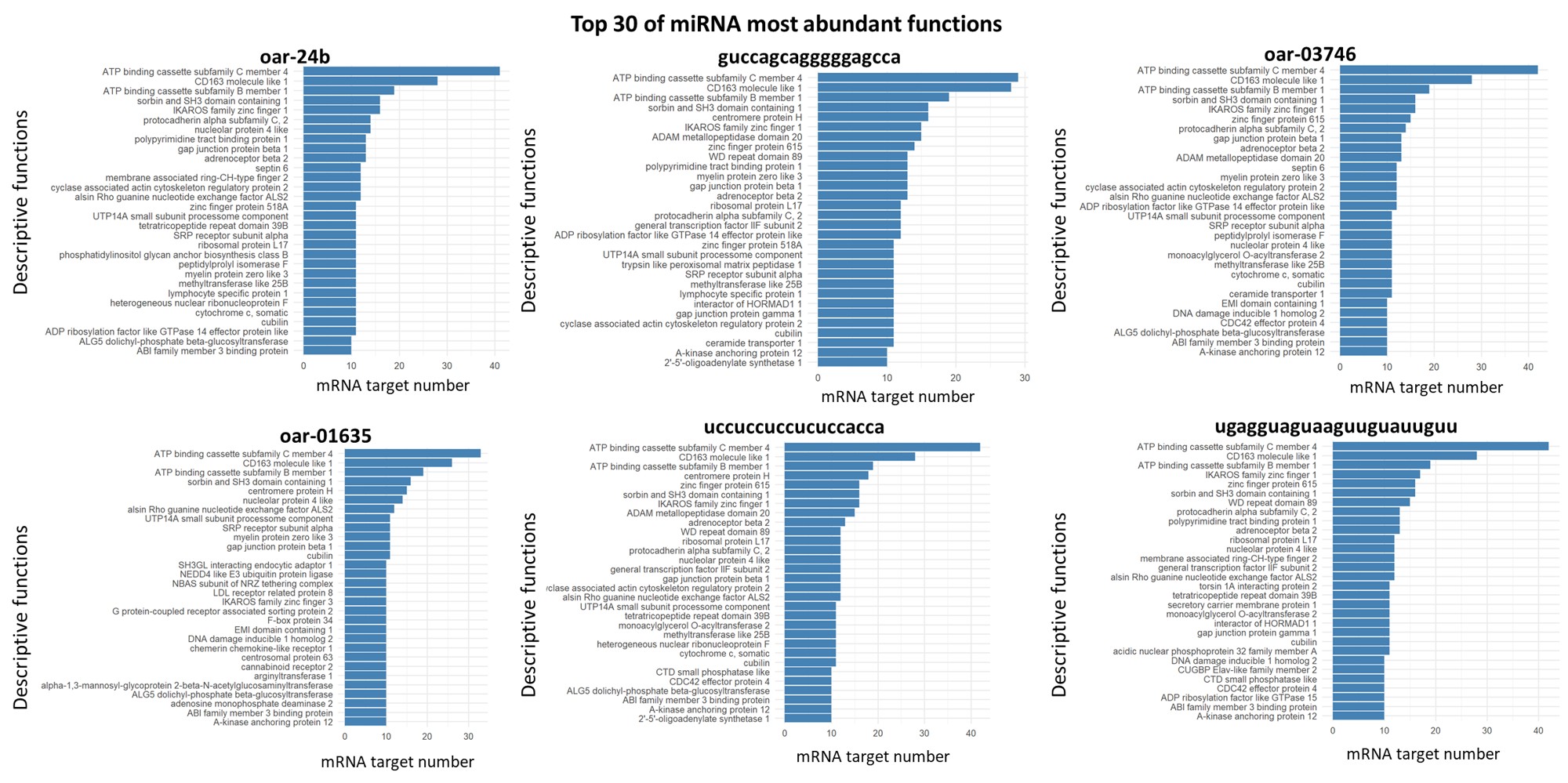

### Supplemental Figure5

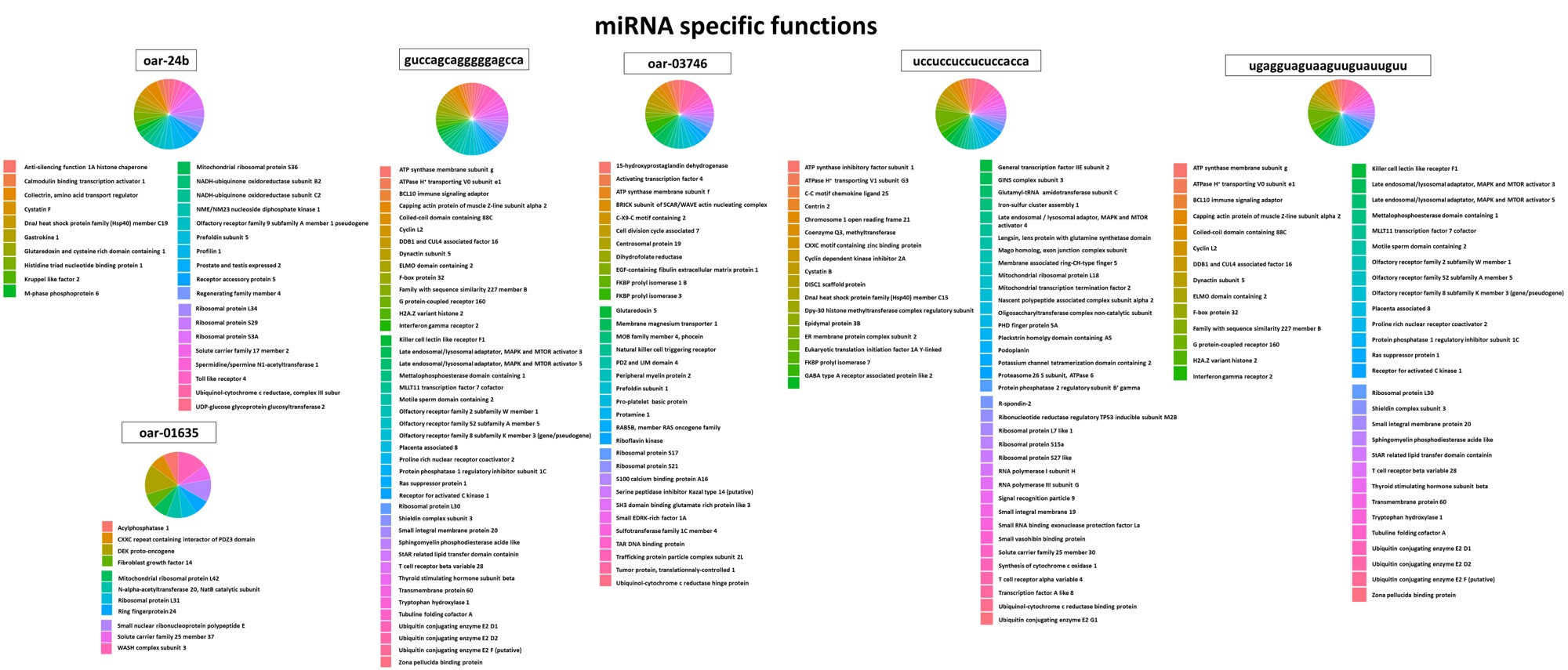
