## Supplemental Table1 for "Characterization of ovine follicular fluid and granulosa cell-derived extracellular vesicles and their miRNA cargo following in vitro exposure to bisphenols A and S"

| **Target protein** | **Expected molecular weight (kDa)** | **Référence** | **Fournisseur** | **Host species** | **Type** | **Dilution** |
|---|---|---|---|---|---|---|
| CD63 | 30 – 60 | sc-31214 | a Cruz Biotechno | Mouse | Plyclonal | 1 : 1000 |
| CD81 | 21-26 | HPA007234 | Sigma-Aldrich | Rabbit | Polyclonal | 1 : 1000 |
| ALIX | 100 | ZRB1947 | Sigma-Aldrich | Rabbit | Monoclonal | 1 : 1000 |
| HSPA1A (HSP70, both HSP73 and HSP72 forms) | 70 | SAB4200714 | Sigma-Aldrich | Mouse | Monoclonal | 1 : 1000 |
| A-Tubulin | 55 | T6074 | Sigma-Aldrich | Mouse | Monoclonal | 1 : 1000 |
| Calnexin | 70 | AB2301 | Sigma-Aldrich | Rabbit | Polyclonal | 1 : 1000 |

| Identity % between the host species protein and ovine protein |
|---|
| 92% |
| 90% |
| 100% |
| 99% |
| 95% |
| 97% |
