## Supplemental Table2 for "Characterization of ovine follicular fluid and granulosa cell-derived extracellular vesicles and their miRNA cargo following in vitro exposure to bisphenols A and S"

| Number | Sample name | Sequencing Sample Name | Nanodrop RNA Concentration after extraction (ng/µl) | Nanodrop Ratio A260/280 nm | Nanodrop Ratio A260/230 nm | RIN |
|---|---|---|---|---|---|---|
| 1 | FF EVs-1 | RS2201-FF-CG-2 | 54 | 1.95 | 1.7 | 3.8 |
| 2 | FF EVs-2 | RS2201-FF-CG-5 | 45.6 | 1.94 | 1.96 | 3.1 |
| 3 | FF EVs-3 | RS2201-FF-CG-6 | 64.4 | 1.97 | 1.89 | 2.8 |
| 4 | FF EVs-4 | RS2201-FF-CG-8 | 98.8 | 1.96 | 1.54 | 2.5 |
| 5 | FF EVs-5 | RS2201-FF-CG-9 | 118.4 | 2.05 | 1.68 | 2.5 |
| 6 | FF EVs-6 | RS2201-FF-CG-10 | 93.6 | 2.1 | 2.3 | 2.5 |
| 7 | FF EVs-7 | RS2201-FF-CG-11 | 106.5 | 1.96 | 1.82 | 2.6 |
| 8 | FF EVs-8 | RS2201-FF-CG-12 | 128.4 | 1.95 | 1.43 | 1.8 |
| 9 | FF EVs-9 | RS2201-FF-CG-13 | 87.7 | 1.87 | 1.18 | 2.4 |
| 10 | FF EVs-10 | RS2201-FF-CG-14 | 79 | 2.01 | 1.84 | 2.2 |
| 11 | CM EVs-CTRL-1 | RS2201-MC-18 | 21 | 1.5 | 0.89 | 0 |
| 12 | CM EVs-BPA-1 | RS2201-MC-19 | 14.6 | 1.54 | 0.78 | 0 |
| 13 | CM EVs-BPS-1 | RS2201-MC-20 | 12.6 | 1.17 | 0.74 | 0 |
| 14 | CM EVs-CTRL-2 | RS2201-MC-27 | 29.3 | 1.52 | 0.98 | 0 |
| 15 | CM EVs-BPA-2 | RS2201-MC-28 | 21 | 1.43 | 1.13 | 0 |
| 16 | CM EVs-BPS-2 | RS2201-MC-29 | 34 | 1.5 | 1 | 0 |
| 17 | CM EVs-CTRL-3 | RS2201-MC-30 | 37.3 | 1.43 | 0.99 | 0 |
| 18 | CM EVs-BPA-3 | RS2201-MC-31 | 103.8 | 1.35 | 0.68 | 0 |
| 19 | CM EVs-BPS-3 | RS2201-MC-32 | 24.9 | 1.49 | 0.88 | 0 |
| 20 | CM EVs-CTRL-4 | RS2201-MC-36 | 33.9 | 1.42 | 0.94 | 0 |
| 21 | CM EVs-BPA-4 | RS2201-MC-37 | 34.9 | 1.43 | 0.84 | 0 |
| 22 | CM EVs-BPS-4 | RS2201-MC-38 | 27.6 | 1.41 | 0.89 | 0 |
| 23 | CM EVs-CTRL-5 | RS2201-MC-39 | 22.2 | 1.5 | 1.06 | 0 |
| 24 | CM EVs-BPA-5 | RS2201-MC-40 | 24.7 | 1.5 | 1.23 | 0 |
| 25 | CM EVs-BPS-5 | RS2201-MC-41 | 68.5 | 1.35 | 0.66 | 0 |
| 26 | CM EVs-CTRL-6 | RS2201-MC-42 | 36.2 | 1.46 | 0.54 | 0 |
| 27 | CM EVs-BPA-6 | RS2201-MC-43 | 76.3 | 1.28 | 0.57 | 0 |
| 28 | CM EVs-BPS-6 | RS2201-MC-44 | 31.6 | 1.42 | 0.85 | 0 |
| 29 | CM EVs-CTRL-7 | RS2201-MC-45 | 36.9 | 1.45 | 0.77 | 0 |
| 30 | CM EVs-BPA-7 | RS2201-MC-46 | 32.9 | 1.48 | 0.81 | 0 |
| 31 | CM EVs-BPS-7 | RS2201-MC-47 | 33.2 | 1.5 | 0.71 | 0 |
| 32 | CM EVs-CTRL-8 | RS2201-MC-48 | 14.8 | 1.56 | 0.91 | 0 |
| 33 | CM EVs-BPA-8 | RS2201-MC-49 | 22.5 | 1.47 | 0.92 | 0 |
| 34 | CM EVs-BPS-8 | RS2201-MC-50 | 18.6 | 1.57 | 0.82 | 0 |
| 35 | CM EVs-CTRL-9 | RS2201-MC-51 | 15.3 | 1.56 | 0.9 | 0 |
| 36 | CM EVs-BPA-9 | RS2201-MC-52 | 21.4 | 1.58 | 0.79 | 0 |
| 37 | CM EVs-BPS-9 | RS2201-MC-53 | 23.1 | 1.53 | 0.95 | 0 |
| 38 | CM EVs-CTRL-10 | RS2201-MC-54 | 12.3 | 1.58 | 0.84 | 0 |
| 39 | CM EVs-BPA-10 | RS2201-MC-55 | 25.5 | 1.58 | 0.72 | 0 |
| 40 | CM EVs-BPS-10 | RS2201-MC-56 | 20.7 | 1.56 | 0.76 | 0 |
| 41 | 0h GC-1 | RS2201-FF-CG-58 | 296.9 | 2.09 | 2.23 | 9.1 |
| 42 | 0h GC-2 | RS2201-FF-CG-61 | 34.5 | 1.78 | 1.35 | 9.4 |
| 43 | 0h GC-3 | RS2201-FF-CG-62 | 39.2 | 1.86 | 1.63 | 9.2 |
| 44 | 0h GC-4 | RS2201-FF-CG-64 | 104.6 | 1.92 | 1.64 | 9.2 |

| | | | | | | |
|---|---|---|---|---|---|---|
| 45 | 0h GC-5 | RS2201-FF-CG-65 | 62.3 | 1.92 | 1.63 | 9.3 |
| 46 | 0h GC-6 | RS2201-FF-CG-66 | 83.2 | 1.92 | 1.77 | 8.9 |
| 47 | 0h GC-7 | RS2201-FF-CG-67 | 42.5 | 1.85 | 1.65 | 9.3 |
| 48 | 0h GC-8 | RS2201-FF-CG-68 | 57.7 | 1.85 | 1.54 | 8.8 |
| 49 | 0h GC-9 | RS2201-FF-CG-69 | 39.9 | 1.7 | 1.15 | 8.9 |
| 50 | 0h GC-10 | RS2201-FF-CG-70 | 35 | 1.92 | 2.24 | 9.3 |
| 51 | 48h GC-CTRL-1 | RS2201-FF-CG-74 | 110.3 | 2 | 1.78 | 10 |
| 52 | 48h GC-BPA-1 | RS2201-FF-CG-75 | 107.1 | 1.94 | 1.67 | 10 |
| 53 | 48h GC-BPS-1 | RS2201-FF-CG-76 | 116.3 | 1.97 | 1.72 | 10 |
| 54 | 48h GC-CTRL-2 | RS2201-FF-CG-83 | 50.08 | 1.799 | 1.669 | 10 |
| 55 | 48h GC-BPA-2 | RS2201-FF-CG-84 | 49.36 | 1.781 | 1.677 | 10 |
| 56 | 48h GC-BPS-2 | RS2201-FF-CG-85 | 56.32 | 1.76 | 1.703 | 10 |
| 57 | 48h GC-CTRL-3 | RS2201-FF-CG-86 | 69.48 | 1.825 | 1.69 | 10 |
| 58 | 48h GC-BPA-3 | RS2201-FF-CG-87 | 70.2 | 1.796 | 1.738 | 9.9 |
| 59 | 48h GC-BPS-3 | RS2201-FF-CG-88 | 69.6 | 1.82 | 1.759 | 10 |
| 60 | 48h GC-CTRL-4 | RS2201-FF-CG-92 | 49.84 | 1.731 | 1.612 | 9.7 |
| 61 | 48h GC-BPA-4 | RS2201-FF-CG-93 | 96 | 1.335 | 0.794 | 9.2 |
| 62 | 48h GC-BPS-4 | RS2201-FF-CG-94 | 59.28 | 1.625 | 1.636 | 9.3 |
| 63 | 48h GC-CTRL-5 | RS2201-FF-CG-95 | 79.68 | 1.862 | 1.914 | 9.7 |
| 64 | 48h GC-BPA-5 | RS2201-FF-CG-96 | 82.48 | 1.82 | 1.799 | 9.2 |
| 65 | 48h GC-BPS-5 | RS2201-FF-CG-97 | 79.96 | 1.822 | 1.771 | 9.5 |
| 66 | 48h GC-CTRL-6 | RS2201-FF-CG-2-98 | 72.6 | 1.762 | 1.813 | 9.6 |
| 67 | 48h GC-BPA-6 | RS2201-FF-CG-99 | 62.24 | 1.748 | 1.662 | 9.4 |
| 68 | 48h GC-BPS-6 | RS2201-FF-CG-100 | 71.56 | 1.739 | 1.756 | 9.4 |
| 69 | 48h GC-CTRL-7 | RS2201-FF-CG-101 | 69.8 | 1.81 | 1.66 | 10 |
| 70 | 48h GC-BPA-7 | RS2201-FF-CG-102 | 57.04 | 1.859 | 1.884 | 10 |
| 71 | 48h GC-BPS-7 | RS2201-FF-CG-103 | 71.16 | 1.851 | 1.746 | 10 |
| 72 | 48h GC-CTRL-8 | RS2201-FF-CG-104 | 84.72 | 1.856 | 1.912 | 10 |
| 73 | 48h GC-BPA-8 | RS2201-FF-CG-105 | 77.68 | 1.841 | 1.927 | 10 |
| 74 | 48h GC-BPS-8 | RS2201-FF-CG-106 | 77.64 | 1.897 | 1.814 | 10 |
| 75 | 48h GC-CTRL-9 | RS2201-FF-CG-107 | 29.72 | 1.568 | 1.532 | 9.9 |
| 76 | 48h GC-BPA-9 | RS2201-FF-CG-108 | 28.32 | 1.516 | 1.351 | 9.8 |
| 77 | 48h GC-BPS-9 | RS2201-FF-CG-109 | 23.84 | 1.585 | 1.447 | 9.9 |
| 78 | 48h GC-CTRL-10 | RS2201-FF-CG-110 | 38.96 | 1.752 | 2.059 | 10 |
| 79 | 48h GC-BPA-10 | RS2201-FF-CG-111 | 44.88 | 1.662 | 1.228 | 10 |
| 80 | 48h GC-BPS-10 | RS2201-FF-CG-112 | 40.16 | 1.761 | 1.965 | 10 |

BPA = Bisphenol A, BPS = Bisphenol S ; CTRL = Control ; FF = Follicular fluid ; GC = Granulosa cells ; CM = Conditioned media

| DropSense16 Concentration 260nm (ng/µl) | cDrop RNA (ng/µl) | cDrop ratio A260/230nm | cDrop ratio A260/280nm |
|---|---|---|---|
| 52.49 | 35.79 | 1.6 | 1.84 |
| 42.74 | 3.84 | 2.46 | 1.79 |
| 64.46 | 37.15 | 1.94 | 1.87 |
| 91.3 | 52.93 | 1.82 | 1.91 |
| 122.72 | 93.78 | 1.73 | 1.93 |
| 99.19 | 79.76 | 1.99 | 1.97 |
| 105.2 | 71.8 | 2.02 | 1.93 |
| 131.35 | 67.59 | 1.52 | 1.88 |
| 89.14 | 24.37 | 1.28 | 1.78 |
| 75.32 | 49.75 | 1.86 | 1.89 |
| 14.95 | 0 | 1.08 | 1.44 |
| 13.64 | 0 | 0.92 | 1.39 |
| 15.91 | 0 | 1.38 | 1.44 |
| 19.88 | 0 | 1.38 | 1.38 |
| 19.26 | 0 | 1.68 | 1.4 |
| 21.14 | 0 | 1.39 | 1.38 |
| 31.26 | 0 | 1.15 | 1.38 |
| 71.23 | 0 | 0.68 | 1.23 |
| 18.35 | 0 | 1.02 | 1.42 |
| 25.53 | 0 | 1.49 | 1.31 |
| 23.16 | 0 | 1.28 | 1.31 |
| 16.43 | 0 | 2.03 | 1.25 |
| 20.72 | 0 | 1.31 | 1.35 |
| 19.22 | 0 | 1.61 | 1.4 |
| 45.08 | 0 | 0.69 | 1.22 |
| 16.79 | 0 | 1.38 | 1.39 |
| 45.03 | 0 | 0.63 | 1.23 |
| 18.48 | 0 | 1.26 | 1.44 |
| 15.21 | 0 | 1.21 | 1.35 |
| 12.2 | 0 | 1.62 | 1.38 |
| 15.43 | 0 | 0.87 | 1.38 |
| 6.81 | 0 | 6.13 | 1.24 |
| 14.19 | 0 | 1.24 | 1.37 |
| 9.58 | 0 | 1.01 | 1.36 |
| 10.45 | 0 | 1.05 | 1.4 |
| 11.82 | 0 | 1 | 1.39 |
| 11.54 | 0 | 3.85 | 1.24 |
| 3.39 | 0 | -1.22 | 0.97 |
| 13.75 | 0 | 0.82 | 1.39 |
| 9.01 | 0.25 | 1 | 1.41 |
| 260.38 | 228.7 | 2.14 | 2.02 |
| 26.01 | 5.28 | 2.86 | 1.69 |
| 27.13 | 13.92 | 1.66 | 1.74 |
| 69.03 | 31.06 | 2.09 | 1.88 |

| | | | |
|---|---|---|---|
| 48.74 | 21.08 | 1.94 | 1.88 |
| 61.86 | 43.54 | 2.11 | 1.76 |
| 35.13 | 16.08 | 1.88 | 1.81 |
| 49.71 | 32.88 | 1.87 | 1.85 |
| 25.45 | 0 | 2.47 | 1.63 |
| 32.95 | 0.85 | 1.88 | 1.76 |
| 90.76 | 62.3 | 2 | 1.97 |
| 83.43 | 42.84 | 2.22 | 1.92 |
| 94.41 | 59.53 | 2.37 | 1.98 |
| 49.07 | 25 | 1.78 | 1.79 |
| 45.84 | 29.66 | 1.86 | 1.77 |
| 54.73 | 27.46 | 2 | 1.79 |
| 68.11 | 33.49 | 1.77 | 1.81 |
| 65.73 | 34.11 | 1.94 | 1.79 |
| 68.46 | 42.82 | 1.9 | 1.84 |
| 49.78 | 27.72 | 1.73 | 1.76 |
| 85.77 | 9.38 | 0.87 | 1.38 |
| 59.49 | 25.33 | 1.74 | 1.71 |
| 78.48 | 59.09 | 2.02 | 1.83 |
| 77.5 | 45.64 | 2.18 | 1.86 |
| 72.2 | 38.58 | 2.37 | 1.84 |
| 71.57 | 42.83 | 1.98 | 1.81 |
| 55.7 | 23.75 | 1.95 | 1.78 |
| 65.12 | 30.81 | 2.37 | 1.82 |
| 66.34 | 35.38 | 1.93 | 1.9 |
| 56.71 | 29.71 | 2 | 1.95 |
| 72.94 | 46.55 | 1.93 | 1.91 |
| 83.85 | 52.27 | 2.27 | 1.9 |
| 87.35 | 31.55 | 1.54 | 1.86 |
| 76.57 | 58.74 | 2 | 1.84 |
| 31.15 | 24.51 | 0.43 | 1.78 |
| 31.26 | 9.6 | 1.8 | 1.75 |
| 21.81 | 5.2 | 4.31 | 1.82 |
| 37.72 | 29.79 | 2.09 | 1.89 |
| 40.47 | 22.79 | 1.47 | 1.78 |
| 44.39 | 33.5 | 1.97 | 1.84 |

; EVs = Extracellular vesicles
