## Supplemental Table3 for "Characterization of ovine follicular fluid and granulosa cell-derived extracellular vesicles and their miRNA cargo following in vitro exposure to bisphenols A and S"

| | | | RNA Biotype (%) | | | |
|---|---|---|---|---|---|---|
| Samples | Condition | Number of replicates | miRNA | Unknown | Artifact | tRNA |
| FF-EVs | - | 10 | 38.47 | 61.23 | 0.27 | 0.03 |
| 0h GC | - | 10 | 14.46 | 68.64 | 0.08 | 16.82 |
| 48h GC | Control | 10 | 22.57 | 66.39 | 0.1 | 10.93 |
| 48h GC | BPA | 10 | 21.65 | 66.82 | 0.09 | 11.44 |
| 48h GC | BPS | 10 | 22.61 | 66.67 | 0.09 | 10.64 |
| 48h CM-EVs | Control | 10 | 0.34 | 98.87 | 0.28 | 0.5 |
| 48h CM-EVs | BPA | 10 | 0.31 | 98.78 | 0.26 | 0.65 |
| 48h CM-EVs | BPS | 10 | 0.28 | 99.2 | 0.26 | 0.25 |

BPA = Bisphenol A, BPS = Bisphenol S ; CTRL = Control ; FF = Follicular fluid ; GC = Granulosa cells ; CM = Conditioned

| |
|---|
| rRNA |
| 0.0007 |
| 0.0005 |
| 0.0005 |
| 0.0006 |
| 0.0004 |
| 0.003 |
| 0.0024 |
| 0.0035 |

media ; EVs = Extracellular vesicles
