## Supplemental Table4 for "Characterization of ovine follicular fluid and granulosa cell-derived extracellular vesicles and their miRNA cargo following in vitro exposure to bisphenols A and S"

| Sample Number | Sample name | Sequencing Identification | Analysis identification | time |
|---|---|---|---|---|
| 41 | 0h GC-1 | RS2201-FF-CG-58 | GC_0_NA_58 | 0 |
| 42 | 0h GC-2 | RS2201-FF-CG-61 | GC_0_NA_61 | 0 |
| 43 | 0h GC-3 | RS2201-FF-CG-62 | GC_0_NA_62 | 0 |
| 44 | 0h GC-4 | RS2201-FF-CG-64 | GC_0_NA_64 | 0 |
| 45 | 0h GC-5 | RS2201-FF-CG-65 | GC_0_NA_65 | 0 |
| 46 | 0h GC-6 | RS2201-FF-CG-66 | GC_0_NA_66 | 0 |
| 47 | 0h GC-7 | RS2201-FF-CG-67 | GC_0_NA_67 | 0 |
| 48 | 0h GC-8 | RS2201-FF-CG-68 | GC_0_NA_68 | 0 |
| 49 | 0h GC-9 | RS2201-FF-CG-69 | GC_0_NA_69 | 0 |
| 50 | 0h GC-10 | RS2201-FF-CG-70 | GC_0_NA_70 | 0 |
| 11 | 48h CM EVs-CTRL-1 | RS2201-MC-18 | CM_48_CTRL_18 | 48 |
| 12 | 48h CM EVs-BPA-1 | RS2201-MC-19 | CM_48_BPA_19 | 48 |
| 13 | 48h CM EVs-BPS-1 | RS2201-MC-20 | CM_48_BPS_20 | 48 |
| 14 | 48h CM EVs-CTRL-2 | RS2201-MC-27 | CM_48_CTRL_27 | 48 |
| 15 | 48h CM EVs-BPA-2 | RS2201-MC-28 | CM_48_BPA_28 | 48 |
| 16 | 48h CM EVs-BPS-2 | RS2201-MC-29 | CM_48_BPS_29 | 48 |
| 17 | 48h CM EVs-CTRL-3 | RS2201-MC-30 | CM_48_CTRL_30 | 48 |
| 18 | 48h CM EVs-BPA-3 | RS2201-MC-31 | CM_48_BPA_31 | 48 |
| 19 | 48h CM EVs-BPS-3 | RS2201-MC-32 | CM_48_BPS_32 | 48 |
| 20 | 48h CM EVs-CTRL-4 | RS2201-MC-36 | CM_48_CTRL_36 | 48 |
| 21 | 48h CM EVs-BPA-4 | RS2201-MC-37 | CM_48_BPA_37 | 48 |
| 22 | 48h CM EVs-BPS-4 | RS2201-MC-38 | CM_48_BPS_38 | 48 |
| 23 | 48h CM EVs-CTRL-5 | RS2201-MC-39 | CM_48_CTRL_39 | 48 |
| 24 | 48h CM EVs-BPA-5 | RS2201-MC-40 | CM_48_BPA_40 | 48 |
| 25 | 48h CM EVs-BPS-5 | RS2201-MC-41 | CM_48_BPS_41 | 48 |
| 26 | 48h CM EVs-CTRL-6 | RS2201-MC-42 | CM_48_CTRL_42 | 48 |
| 27 | 48h CM EVs-BPA-6 | RS2201-MC-43 | CM_48_BPA_43 | 48 |
| 28 | 48h CM EVs-BPS-6 | RS2201-MC-44 | CM_48_BPS_44 | 48 |
| 29 | 48h CM EVs-CTRL-7 | RS2201-MC-45 | CM_48_CTRL_45 | 48 |
| 30 | 48h CM EVs-BPA-7 | RS2201-MC-46 | CM_48_BPA_46 | 48 |
| 31 | 48h CM EVs-BPS-7 | RS2201-MC-47 | CM_48_BPS_47 | 48 |
| 32 | 48h CM EVs-CTRL-8 | RS2201-MC-48 | CM_48_CTRL_48 | 48 |
| 33 | 48h CM EVs-BPA-8 | RS2201-MC-49 | CM_48_BPA_49 | 48 |
| 34 | 48h CM EVs-BPS-8 | RS2201-MC-50 | CM_48_BPS_50 | 48 |
| 35 | 48h CM EVs-CTRL-9 | RS2201-MC-51 | CM_48_CTRL_51 | 48 |
| 36 | 48h CM EVs-BPA-9 | RS2201-MC-52 | CM_48_BPA_52 | 48 |
| 37 | 48h CM EVs-BPS-9 | RS2201-MC-53 | CM_48_BPS_53 | 48 |
| 38 | 48h CM EVs-CTRL-10 | RS2201-MC-54 | CM_48_CTRL_54 | 48 |
| 39 | 48h CM EVs-BPA-10 | RS2201-MC-55 | CM_48_BPA_55 | 48 |
| 40 | 48h CM EVs-BPS-10 | RS2201-MC-56 | CM_48_BPS_56 | 48 |
| 51 | 48h GC-CTRL-1 | RS2201-FF-CG-83 | GC_48_CTRL_74 | 48 |
| 52 | 48h GC-BPA-1 | RS2201-FF-CG-75 | GC_48_BPA_75 | 48 |
| 53 | 48h GC-BPS-1 | RS2201-FF-CG-76 | GC_48_BPS_76 | 48 |
| 54 | 48h GC-CTRL-2 | RS2201-FF-CG-86 | GC_48_CTRL_83 | 48 |
| 55 | 48h GC-BPA-2 | RS2201-FF-CG-84 | GC_48_BPA_84 | 48 |
| 56 | 48h GC-BPS-2 | RS2201-FF-CG-85 | GC_48_BPS_85 | 48 |
| 57 | 48h GC-CTRL-3 | RS2201-FF-CG-92 | GC_48_CTRL_86 | 48 |
| 58 | 48h GC-BPA-3 | RS2201-FF-CG-87 | GC_48_BPA_87 | 48 |
| 59 | 48h GC-BPS-3 | RS2201-FF-CG-88 | GC_48_BPS_88 | 48 |

| | | | | |
|---|---|---|---|---|
| 60 | 48h GC-CTRL-4 | RS2201-FF-CG-95 | GC_48_CTRL_92 | 48 |
| 61 | 48h GC-BPA-4 | RS2201-FF-CG-93 | GC_48_BPA_93 | 48 |
| 62 | 48h GC-BPS-4 | RS2201-FF-CG-94 | GC_48_BPS_94 | 48 |
| 63 | 48h GC-CTRL-5 | RS2201-FF-CG-2-98 | GC_48_CTRL_95 | 48 |
| 64 | 48h GC-BPA-5 | RS2201-FF-CG-96 | GC_48_BPA_96 | 48 |
| 65 | 48h GC-BPS-5 | RS2201-FF-CG-97 | GC_48_BPS_97 | 48 |
| 66 | 48h GC-CTRL-6 | RS2201-FF-CG-74 | GC_48_CTRL_98 | 48 |
| 67 | 48h GC-BPA-6 | RS2201-FF-CG-99 | GC_48_BPA_99 | 48 |
| 68 | 48h GC-BPS-6 | RS2201-FF-CG-100 | GC_48_BPS_100 | 48 |
| 69 | 48h GC-CTRL-7 | RS2201-FF-CG-101 | GC_48_CTRL_101 | 48 |
| 70 | 48h GC-BPA-7 | RS2201-FF-CG-102 | GC_48_BPA_102 | 48 |
| 71 | 48h GC-BPS-7 | RS2201-FF-CG-103 | GC_48_BPS_103 | 48 |
| 72 | 48h GC-CTRL-8 | RS2201-FF-CG-104 | GC_48_CTRL_104 | 48 |
| 73 | 48h GC-BPA-8 | RS2201-FF-CG-105 | GC_48_BPA_105 | 48 |
| 74 | 48h GC-BPS-8 | RS2201-FF-CG-106 | GC_48_BPS_106 | 48 |
| 75 | 48h GC-CTRL-9 | RS2201-FF-CG-107 | GC_48_CTRL_107 | 48 |
| 76 | 48h GC-BPA-9 | RS2201-FF-CG-108 | GC_48_BPA_108 | 48 |
| 77 | 48h GC-BPS-9 | RS2201-FF-CG-109 | GC_48_BPS_109 | 48 |
| 78 | 48h GC-CTRL-10 | RS2201-FF-CG-110 | GC_48_CTRL_110 | 48 |
| 79 | 48h GC-BPA-10 | RS2201-FF-CG-111 | GC_48_BPA_111 | 48 |
| 80 | 48h GC-BPS-10 | RS2201-FF-CG-112 | GC_48_BPS_112 | 48 |
| 6 | FF EVs-6 | RS2201-FF-CG-10 | FF_0_NA_10 | 0 |
| 7 | FF EVs-7 | RS2201-FF-CG-11 | FF_0_NA_11 | 0 |
| 8 | FF EVs-8 | RS2201-FF-CG-12 | FF_0_NA_12 | 0 |
| 9 | FF EVs-9 | RS2201-FF-CG-13 | FF_0_NA_13 | 0 |
| 10 | FF EVs-10 | RS2201-FF-CG-14 | FF_0_NA_14 | 0 |
| 1 | FF EVs-1 | RS2201-FF-CG-2 | FF_0_NA_2 | 0 |
| 2 | FF EVs-2 | RS2201-FF-CG-5 | FF_0_NA_5 | 0 |
| 3 | FF EVs-3 | RS2201-FF-CG-6 | FF_0_NA_6 | 0 |
| 4 | FF EVs-4 | RS2201-FF-CG-8 | FF_0_NA_8 | 0 |
| 5 | FF EVs-5 | RS2201-FF-CG-9 | FF_0_NA_9 | 0 |

BPA = Bisphenol A, BPS = Bisphenol S ; CTRL = Control ; FF = Follicular fluid ; GC = Granulosa cells ; CM = Co

| tissue-time | condition | tissue | rRNA | tRNA | Artifact | Unknown |
|---|---|---|---|---|---|---|
| GC_T0 | NA | GC | 93 | 2622635 | 11069 | 7492509 |
| GC_T0 | NA | GC | 73 | 2794324 | 8911 | 8959366 |
| GC_T0 | NA | GC | 100 | 4126248 | 8037 | 10316368 |
| GC_T0 | NA | GC | 71 | 1370605 | 13300 | 8642805 |
| GC_T0 | NA | GC | 94 | 1307809 | 14652 | 10321088 |
| GC_T0 | NA | GC | 72 | 2196557 | 22204 | 10097359 |
| GC_T0 | NA | GC | 72 | 3161678 | 10973 | 10133018 |
| GC_T0 | NA | GC | 86 | 4132589 | 11971 | 11227257 |
| GC_T0 | NA | GC | 68 | 2671702 | 11734 | 10880507 |
| GC_T0 | NA | GC | 116 | 2647429 | 11618 | 9908661 |
| CM_T48 | CTRL | CM | 193 | 334427 | 23200 | 11008423 |
| CM_T48 | BPA | CM | 214 | 405760 | 15778 | 10517392 |
| CM_T48 | BPS | CM | 256 | 159000 | 26488 | 11893944 |
| CM_T48 | CTRL | CM | 212 | 1613 | 29558 | 9804074 |
| CM_T48 | BPA | CM | 65 | 1558 | 27986 | 9110661 |
| CM_T48 | BPS | CM | 1190 | 437 | 17956 | 8465424 |
| CM_T48 | CTRL | CM | 249 | 1567 | 29247 | 7881555 |
| CM_T48 | BPA | CM | 239 | 995 | 31984 | 5716345 |
| CM_T48 | BPS | CM | 272 | 714 | 21939 | 10854683 |
| CM_T48 | CTRL | CM | 427 | 8393 | 23702 | 9414661 |
| CM_T48 | BPA | CM | 388 | 2212 | 25801 | 8177517 |
| CM_T48 | BPS | CM | 556 | 1622 | 21856 | 8531074 |
| CM_T48 | CTRL | CM | 1070 | 9622 | 21145 | 10165948 |
| CM_T48 | BPA | CM | 671 | 937 | 22837 | 7014004 |
| CM_T48 | BPS | CM | 93 | 4701 | 26980 | 5910836 |
| CM_T48 | CTRL | CM | 506 | 3725 | 38649 | 9717743 |
| CM_T48 | BPA | CM | 388 | 2 | 24903 | 7407398 |
| CM_T48 | BPS | CM | 557 | 3059 | 23468 | 8756299 |
| CM_T48 | CTRL | CM | 0 | 24381 | 29867 | 9135632 |
| CM_T48 | BPA | CM | 35 | 35718 | 24553 | 9709986 |
| CM_T48 | BPS | CM | 39 | 51677 | 22274 | 9047104 |
| CM_T48 | CTRL | CM | 60 | 16060 | 23053 | 7576924 |
| CM_T48 | BPA | CM | 29 | 46403 | 25917 | 8700040 |
| CM_T48 | BPS | CM | 0 | 94097 | 19683 | 9210182 |
| CM_T48 | CTRL | CM | 58 | 39688 | 16610 | 8387088 |
| CM_T48 | BPA | CM | 36 | 35226 | 16506 | 8384183 |
| CM_T48 | BPS | CM | 0 | 14758 | 26328 | 9130604 |
| CM_T48 | CTRL | CM | 0 | 20597 | 24617 | 7449261 |
| CM_T48 | BPA | CM | 31 | 35509 | 13406 | 11356833 |
| CM_T48 | BPS | CM | 28 | 28377 | 27307 | 8441870 |
| GC_T48 | CTRL | GC | 52 | 2307203 | 13877 | 8306723 |
| GC_T48 | BPA | GC | 58 | 2657755 | 10157 | 8001740 |
| GC_T48 | BPS | GC | 51 | 2284112 | 13672 | 7072368 |
| GC_T48 | CTRL | GC | 79 | 1251098 | 17828 | 8050203 |
| GC_T48 | BPA | GC | 64 | 1136160 | 12319 | 7649863 |
| GC_T48 | BPS | GC | 55 | 1061992 | 11452 | 7767249 |
| GC_T48 | CTRL | GC | 81 | 1211278 | 16026 | 8342828 |
| GC_T48 | BPA | GC | 46 | 1254912 | 14107 | 6080675 |
| GC_T48 | BPS | GC | 71 | 1371666 | 14899 | 9176532 |

| | | | | | | |
|---|---|---|---|---|---|---|
| GC_T48 | CTRL | GC | 83 | 997714 | 14260 | 7223920 |
| GC_T48 | BPA | GC | 122 | 1224193 | 17243 | 7871735 |
| GC_T48 | BPS | GC | 59 | 967416 | 11094 | 7146735 |
| GC_T48 | CTRL | GC | 83 | 954211 | 11837 | 9167269 |
| GC_T48 | BPA | GC | 109 | 1782750 | 20914 | 12905168 |
| GC_T48 | BPS | GC | 77 | 1065571 | 14770 | 13528768 |
| GC_T48 | CTRL | GC | 60 | 1122812 | 10128 | 6541835 |
| GC_T48 | BPA | GC | 146 | 993593 | 17114 | 8403254 |
| GC_T48 | BPS | GC | 18 | 731999 | 7300 | 4555602 |
| GC_T48 | CTRL | GC | 51 | 1760149 | 16023 | 8359428 |
| GC_T48 | BPA | GC | 48 | 1507801 | 12639 | 7509886 |
| GC_T48 | BPS | GC | 47 | 1312770 | 13201 | 8057342 |
| GC_T48 | CTRL | GC | 108 | 2020671 | 27444 | 9285709 |
| GC_T48 | BPA | GC | 51 | 2459738 | 12481 | 8411612 |
| GC_T48 | BPS | GC | 26 | 2124957 | 11974 | 6852671 |
| GC_T48 | CTRL | GC | 84 | 2609882 | 9452 | 8161939 |
| GC_T48 | BPA | GC | 119 | 2699824 | 11663 | 10904922 |
| GC_T48 | BPS | GC | 48 | 1887956 | 7922 | 6828839 |
| GC_T48 | CTRL | GC | 66 | 1525743 | 8094 | 6877226 |
| GC_T48 | BPA | GC | 80 | 1628133 | 13487 | 8127179 |
| GC_T48 | BPS | GC | 66 | 2187373 | 16373 | 8123386 |
| FF | NA | FF | 138 | 2483 | 32017 | 4526799 |
| FF | NA | FF | 66 | 1910 | 22152 | 3154653 |
| FF | NA | FF | 53 | 5255 | 25703 | 3339574 |
| FF | NA | FF | 88 | 6113 | 18901 | 2875400 |
| FF | NA | FF | 73 | 3287 | 26096 | 2426593 |
| FF | NA | FF | 28 | 1645 | 24534 | 4766383 |
| FF | NA | FF | 105 | 3469 | 39866 | 4632721 |
| FF | NA | FF | 51 | 2072 | 33637 | 4215226 |
| FF | NA | FF | 67 | 1989 | 30181 | 3921809 |
| FF | NA | FF | 77 | 1268 | 32312 | 3705578 |

nditioned media ; EVs = Extracellular vesicles

| mirna | mirna_mirdeep2 |
|---|---|
| 2996795 | 85797 |
| 2264990 | 81901 |
| 3467070 | 130861 |
| 4175713 | 102188 |
| 3390350 | 115537 |
| 5586465 | 133163 |
| 3289000 | 113104 |
| 3886327 | 136898 |
| 2543939 | 96463 |
| 2825629 | 119998 |
| 191788 | 8437 |
| 190425 | 11792 |
| 212184 | 7101 |
| 21829 | 932 |
| 21338 | 1062 |
| 8054 | 68 |
| 11399 | 98 |
| 13522 | 141 |
| 10369 | 311 |
| 35961 | 1213 |
| 19425 | 1344 |
| 19642 | 637 |
| 51205 | 1803 |
| 14433 | 80 |
| 23631 | 565 |
| 31977 | 581 |
| 9327 | 160 |
| 28063 | 1401 |
| 39497 | 3042 |
| 51309 | 2462 |
| 126460 | 6495 |
| 85448 | 5131 |
| 63500 | 5649 |
| 79838 | 6985 |
| 56031 | 3721 |
| 39310 | 2895 |
| 27498 | 1914 |
| 16302 | 884 |
| 54938 | 4111 |
| 71915 | 4296 |
| 5042633 | 119269 |
| 5521996 | 137686 |
| 5174985 | 123114 |
| 4566763 | 156483 |
| 4700059 | 129352 |
| 4913475 | 132905 |
| 5000788 | 131150 |
| 3969806 | 106267 |
| 5105473 | 136382 |

| | |
|---|---|
| 4067256 | 109621 |
| 3930888 | 128633 |
| 4033765 | 122157 |
| 5323113 | 147803 |
| 7825012 | 211648 |
| 5558642 | 145233 |
| 4624470 | 132827 |
| 3518231 | 95886 |
| 2745208 | 78047 |
| 5107514 | 150103 |
| 5007834 | 125507 |
| 4765084 | 122785 |
| 5118212 | 127056 |
| 4843341 | 122425 |
| 4709143 | 117437 |
| 3806020 | 117439 |
| 3939344 | 116509 |
| 3080839 | 91618 |
| 3977004 | 106699 |
| 3705808 | 102139 |
| 5423272 | 149767 |
| 7473158 | 488801 |
| 7522285 | 368675 |
| 7272651 | 461933 |
| 6884141 | 473788 |
| 6320351 | 430849 |
| 3923766 | 313834 |
| 6218490 | 492658 |
| 5717670 | 328971 |
| 6698303 | 433484 |
| 6585440 | 450839 |
