## Supplemental Table5 for "Characterization of ovine follicular fluid and granulosa cell-derived extracellular vesicles and their miRNA cargo following in vitro exposure to bisphenols A and S"

| Samples | Condition | Number of replicates | Mean of Total Filtered reads | Mean of Passed Filter reads | Mean of Low Quality reads | Mean of Too Short reads |
|---|---|---|---|---|---|---|
| FF-EVs | - | 10 | 14900394 | 12062558 | 15620 | 2822216 |
| 0h GC | - | 10 | 18346728 | 16243202 | 12017 | 1686223 |
| 48h GC | Control | 10 | 15920819 | 14859691 | 9492 | 1051637 |
| 48h GC | BPA | 10 | 16802138 | 15634997 | 10390 | 1156751 |
| 48h GC | BPS | 10 | 15486735 | 14505400 | 9444 | 971891 |
| 48h CM-EVs | Control | 10 | 15058095 | 10078466 | 20020 | 4959610 |
| 48h CM-EVs | BPA | 10 | 14315706 | 9561012 | 20981 | 4733714 |
| 48h CM-EVs | BPS | 10 | 15110216 | 10055767 | 22227 | 5032222 |

BPA = Bisphenol A, BPS = Bisphenol S ; CTRL = Control ; FF = Follicular fluid ; GC = Granulosa cells ; CM = Conditioned

| % Passed Filter reads | % Low Quality reads | % Too Short reads |
|---|---|---|
| 80.97 | 0.10 | 18.93 |
| 90.68 | 0.07 | 9.25 |
| 93.32 | 0.06 | 6.62 |
| 92.96 | 0.06 | 6.98 |
| 93.59 | 0.06 | 6.35 |
| 66.80 | 0.13 | 33.06 |
| 66.37 | 0.15 | 33.48 |
| 66.26 | 0.15 | 33.59 |

media ; EVs = Extracellular vesicles
