## Supplemental Table6 for "Characterization of ovine follicular fluid and granulosa cell-derived extracellular vesicles and their miRNA cargo following in vitro exposure to bisphenols A and S"

| Sequencing Identification | Analysis identification | Identification | time |
|---|---|---|---|
| RS2201-FF-CG-2 | FF_0_NA_2 | FF-CG-2_S16_m | 0 |
| RS2201-FF-CG-5 | FF_0_NA_5 | FF-CG-5_S1_ma | 0 |
| RS2201-FF-CG-6 | FF_0_NA_6 | FF-CG-6_S44_m | 0 |
| RS2201-FF-CG-8 | FF_0_NA_8 | FF-CG-8_S29_m | 0 |
| RS2201-FF-CG-9 | FF_0_NA_9 | FF-CG-9_S35_m | 0 |
| RS2201-FF-CG-10 | FF_0_NA_10 | FF-CG-10_S37_ | 0 |
| RS2201-FF-CG-11 | FF_0_NA_11 | FF-CG-11_S12_ | 0 |
| RS2201-FF-CG-12 | FF_0_NA_12 | FF-CG-12_S48_ | 0 |
| RS2201-FF-CG-13 | FF_0_NA_13 | FF-CG-13_S8_m | 0 |
| RS2201-FF-CG-14 | FF_0_NA_14 | FF-CG-14_S23_ | 0 |
| RS2201-MC-19 | MC_48_BPA_19 | MC-19_S51_mat | 48 |
| RS2201-MC-28 | MC_48_BPA_28 | MC-28_S54_mat | 48 |
| RS2201-MC-31 | MC_48_BPA_31 | MC-31_S57_mat | 48 |
| RS2201-MC-37 | MC_48_BPA_37 | MC-37_S60_mat | 48 |
| RS2201-MC-40 | MC_48_BPA_40 | MC-40_S63_mat | 48 |
| RS2201-MC-43 | MC_48_BPA_43 | MC-43_S66_mat | 48 |
| RS2201-MC-46 | MC_48_BPA_46 | MC-46_S69_mat | 48 |
| RS2201-MC-49 | MC_48_BPA_49 | MC-49_S72_mat | 48 |
| RS2201-MC-52 | MC_48_BPA_52 | MC-52_S75_mat | 48 |
| RS2201-MC-55 | MC_48_BPA_55 | MC-55_S78_mat | 48 |
| RS2201-MC-20 | MC_48_BPS_20 | MC-20_S52_mat | 48 |
| RS2201-MC-29 | MC_48_BPS_29 | MC-29_S55_mat | 48 |
| RS2201-MC-32 | MC_48_BPS_32 | MC-32_S58_mat | 48 |
| RS2201-MC-38 | MC_48_BPS_38 | MC-38_S61_mat | 48 |
| RS2201-MC-41 | MC_48_BPS_41 | MC-41_S64_mat | 48 |
| RS2201-MC-44 | MC_48_BPS_44 | MC-44_S67_mat | 48 |
| RS2201-MC-47 | MC_48_BPS_47 | MC-47_S70_mat | 48 |
| RS2201-MC-50 | MC_48_BPS_50 | MC-50_S73_mat | 48 |
| RS2201-MC-53 | MC_48_BPS_53 | MC-53_S76_mat | 48 |
| RS2201-MC-56 | MC_48_BPS_56 | MC-56_S79_mat | 48 |
| RS2201-MC-18 | MC_48_CTRL_18 | MC-18_S50_mat | 48 |
| RS2201-MC-27 | MC_48_CTRL_27 | MC-27_S53_mat | 48 |
| RS2201-MC-30 | MC_48_CTRL_30 | MC-30_S56_mat | 48 |
| RS2201-MC-36 | MC_48_CTRL_36 | MC-36_S59_mat | 48 |
| RS2201-MC-39 | MC_48_CTRL_39 | MC-39_S62_mat | 48 |
| RS2201-MC-42 | MC_48_CTRL_42 | MC-42_S65_mat | 48 |
| RS2201-MC-45 | MC_48_CTRL_45 | MC-45_S68_mat | 48 |
| RS2201-MC-48 | MC_48_CTRL_48 | MC-48_S71_mat | 48 |
| RS2201-MC-51 | MC_48_CTRL_51 | MC-51_S74_mat | 48 |
| RS2201-MC-54 | MC_48_CTRL_54 | MC-54_S77_mat | 48 |
| RS2201-FF-CG-58 | CG_0_NA_58 | FF-CG-58_S47_ | 0 |
| RS2201-FF-CG-61 | CG_0_NA_61 | FF-CG-61_S2_m | 0 |
| RS2201-FF-CG-62 | CG_0_NA_62 | FF-CG-62_S33_ | 0 |
| RS2201-FF-CG-64 | CG_0_NA_64 | FF-CG-64_S14_ | 0 |
| RS2201-FF-CG-65 | CG_0_NA_65 | FF-CG-65_S27_ | 0 |
| RS2201-FF-CG-66 | CG_0_NA_66 | FF-CG-66_S15_ | 0 |
| RS2201-FF-CG-67 | CG_0_NA_67 | FF-CG-67_S6_m | 0 |
| RS2201-FF-CG-68 | CG_0_NA_68 | FF-CG-68_S36_ | 0 |

| | | | |
|---|---|---|---|
| RS2201-FF-CG-69 | CG_0_NA_69 | FF-CG-69_S43_ | 0 |
| RS2201-FF-CG-70 | CG_0_NA_70 | FF-CG-70_S21_ | 0 |
| RS2201-FF-CG-102 | CG_48_BPA_102 | FF-CG-102_S40 | 48 |
| RS2201-FF-CG-105 | CG_48_BPA_105 | FF-CG-105_S25 | 48 |
| RS2201-FF-CG-108 | CG_48_BPA_108 | FF-CG-108_S41 | 48 |
| RS2201-FF-CG-111 | CG_48_BPA_111 | FF-CG-111_S19 | 48 |
| RS2201-FF-CG-75 | CG_48_BPA_75 | FF-CG-75_S18_ | 48 |
| RS2201-FF-CG-84 | CG_48_BPA_84 | FF-CG-84_S30_ | 48 |
| RS2201-FF-CG-87 | CG_48_BPA_87 | FF-CG-87_S39_ | 48 |
| RS2201-FF-CG-93 | CG_48_BPA_93 | FF-CG-93_S5_m | 48 |
| RS2201-FF-CG-96 | CG_48_BPA_96 | FF-CG-96_S11_ | 48 |
| RS2201-FF-CG-99 | CG_48_BPA_99 | FF-CG-99_S49_ | 48 |
| RS2201-FF-CG-100 | CG_48_BPS_100 | FF-CG-100_S22 | 48 |
| RS2201-FF-CG-103 | CG_48_BPS_103 | FF-CG-103_S20 | 48 |
| RS2201-FF-CG-106 | CG_48_BPS_106 | FF-CG-106_S32 | 48 |
| RS2201-FF-CG-109 | CG_48_BPS_109 | FF-CG-109_S7_ | 48 |
| RS2201-FF-CG-112 | CG_48_BPS_112 | FF-CG-112_S9_ | 48 |
| RS2201-FF-CG-76 | CG_48_BPS_76 | FF-CG-76_S3_m | 48 |
| RS2201-FF-CG-85 | CG_48_BPS_85 | FF-CG-85_S13_ | 48 |
| RS2201-FF-CG-88 | CG_48_BPS_88 | FF-CG-88_S26_ | 48 |
| RS2201-FF-CG-94 | CG_48_BPS_94 | FF-CG-94_S46_ | 48 |
| RS2201-FF-CG-97 | CG_48_BPS_97 | FF-CG-97_S34_ | 48 |
| RS2201-FF-CG-101 | CG_48_CTRL_101 | FF-CG-101_S24 | 48 |
| RS2201-FF-CG-104 | CG_48_CTRL_104 | FF-CG-104_S4_ | 48 |
| RS2201-FF-CG-107 | CG_48_CTRL_107 | FF-CG-107_S17 | 48 |
| RS2201-FF-CG-110 | CG_48_CTRL_110 | FF-CG-110_S45 | 48 |
| RS2201-FF-CG-74 | CG_48_CTRL_74 | FF-CG-2-98_S8 | 48 |
| RS2201-FF-CG-83 | CG_48_CTRL_83 | FF-CG-74_S28_ | 48 |
| RS2201-FF-CG-86 | CG_48_CTRL_86 | FF-CG-83_S38_ | 48 |
| RS2201-FF-CG-92 | CG_48_CTRL_92 | FF-CG-86_S42_ | 48 |
| RS2201-FF-CG-95 | CG_48_CTRL_95 | FF-CG-92_S31_ | 48 |
| RS2201-FF-CG-2-98 | CG_48_CTRL_98 | FF-CG-95_S10_ | 48 |

BPA = Bisphenol A, BPS = Bisphenol S ; CTRL = Control ; FF = Follicular fluid ; GC = Granulosa cells ; CM = C

| | | | mature | | mature_hai |
|---|---|---|---|---|---|
| tissue | tissue-time | condition | Mapped (with MQ>0) | Unmapped | Mapped (with MQ>0) |
| FF | FF | NA | 17.39058331 | 82.60941669 | 26.4865858 |
| FF | FF | NA | 22.52130812 | 77.47869188 | 37.75097979 |
| FF | FF | NA | 22.62708027 | 77.37291973 | 36.99557091 |
| FF | FF | NA | 28.54293151 | 71.45706849 | 41.93389824 |
| FF | FF | NA | 27.87174714 | 72.12825286 | 43.41604906 |
| FF | FF | NA | 27.19429158 | 72.80570842 | 42.77361098 |
| FF | FF | NA | 30.29622161 | 69.70377839 | 49.92728515 |
| FF | FF | NA | 30.18343786 | 69.81656214 | 45.09851015 |
| FF | FF | NA | 30.34149854 | 69.65850146 | 49.51454497 |
| FF | FF | NA | 30.50545285 | 69.49454715 | 51.70206904 |
| CM | CM48 | BPA | 0.899704462 | 99.10029554 | 0.835648307 |
| CM | CM48 | BPA | 0.158376701 | 99.8416233 | 0.138848944 |
| CM | CM48 | BPA | 0.158712548 | 99.84128745 | 0.170407151 |
| CM | CM48 | BPA | 0.145155367 | 99.85484463 | 0.152551053 |
| CM | CM48 | BPA | 0.113167787 | 99.88683221 | 0.156610656 |
| CM | CM48 | BPA | 0.070643706 | 99.92935629 | 0.109816867 |
| CM | CM48 | BPA | 0.256767585 | 99.74323241 | 0.296103918 |
| CM | CM48 | BPA | 0.356515602 | 99.6434844 | 0.378855213 |
| CM | CM48 | BPA | 0.214735691 | 99.78526431 | 0.291580908 |
| CM | CM48 | BPA | 0.234572185 | 99.76542781 | 0.272821934 |
| CM | CM48 | BPS | 0.873748835 | 99.12625117 | 0.878659969 |
| CM | CM48 | BPS | 0.0542997 | 99.9457003 | 0.116808077 |
| CM | CM48 | BPS | 0.05938482 | 99.94061518 | 0.07095432 |
| CM | CM48 | BPS | 0.141143792 | 99.85885621 | 0.148149344 |
| CM | CM48 | BPS | 0.254656584 | 99.74534342 | 0.238342499 |
| CM | CM48 | BPS | 0.252182828 | 99.74781717 | 0.139820378 |
| CM | CM48 | BPS | 0.689199425 | 99.31080057 | 0.703448946 |
| CM | CM48 | BPS | 0.412498871 | 99.58750113 | 0.455266789 |
| CM | CM48 | BPS | 0.138737281 | 99.86126272 | 0.190740772 |
| CM | CM48 | BPS | 0.376720797 | 99.6232792 | 0.475051258 |
| CM | CM48 | CTRL | 0.880487309 | 99.11951269 | 0.817944241 |
| CM | CM48 | CTRL | 0.143054031 | 99.85694597 | 0.140084573 |
| CM | CM48 | CTRL | 0.085154901 | 99.9148451 | 0.129645755 |
| CM | CM48 | CTRL | 0.249485948 | 99.75051405 | 0.190054362 |
| CM | CM48 | CTRL | 0.319341321 | 99.68065868 | 0.236139628 |
| CM | CM48 | CTRL | 0.217784205 | 99.78221579 | 0.161371408 |
| CM | CM48 | CTRL | 0.217450232 | 99.78254977 | 0.247811662 |
| CM | CM48 | CTRL | 0.496525967 | 99.50347403 | 0.591565823 |
| CM | CM48 | CTRL | 0.310313482 | 99.68968652 | 0.362580451 |
| CM | CM48 | CTRL | 0.10248763 | 99.89751237 | 0.165282628 |
| GC | GC_T0 | NA | 13.04840116 | 86.95159884 | 11.62770902 |
| GC | GC_T0 | NA | 8.079029091 | 91.92097091 | 9.178549604 |
| GC | GC_T0 | NA | 9.687658655 | 90.31234134 | 11.15238118 |
| GC | GC_T0 | NA | 16.35119075 | 83.64880925 | 15.96133391 |
| GC | GC_T0 | NA | 11.98051428 | 88.01948572 | 12.22693691 |
| GC | GC_T0 | NA | 17.57556911 | 82.42443089 | 16.83011317 |
| GC | GC_T0 | NA | 10.24869934 | 89.75130066 | 11.25429629 |
| GC | GC_T0 | NA | 10.68971118 | 89.31028882 | 11.20082712 |

| | | | | | |
|---|---|---|---|---|---|
| GC | GC_T0 | NA | 7.932513511 | 92.06748649 | 8.990465188 |
| GC | GC_T0 | NA | 9.448615328 | 90.55138467 | 10.26164804 |
| GC | GC_T48 | BPA | 24.47020263 | 75.52979737 | 16.14466714 |
| GC | GC_T48 | BPA | 20.29034241 | 79.70965759 | 14.33381607 |
| GC | GC_T48 | BPA | 13.05317064 | 86.94682936 | 11.81046947 |
| GC | GC_T48 | BPA | 17.16217274 | 82.83782726 | 13.24659592 |
| GC | GC_T48 | BPA | 24.14534248 | 75.85465752 | 14.24546682 |
| GC | GC_T48 | BPA | 23.21789757 | 76.78210243 | 16.53322193 |
| GC | GC_T48 | BPA | 24.05388691 | 75.94611309 | 15.79424827 |
| GC | GC_T48 | BPA | 20.29989098 | 79.70010902 | 13.40700475 |
| GC | GC_T48 | BPA | 23.91796031 | 76.08203969 | 15.24508915 |
| GC | GC_T48 | BPA | 18.92384222 | 81.07615778 | 10.94524618 |
| GC | GC_T48 | BPS | 23.60023616 | 76.39976384 | 14.947661 |
| GC | GC_T48 | BPS | 22.95344217 | 77.04655783 | 15.17651339 |
| GC | GC_T48 | BPS | 22.90795593 | 77.09204407 | 15.94311405 |
| GC | GC_T48 | BPS | 15.51474707 | 84.48525293 | 13.48211522 |
| GC | GC_T48 | BPS | 21.42972479 | 78.57027521 | 17.5212082 |
| GC | GC_T48 | BPS | 25.24187009 | 74.75812991 | 14.94653713 |
| GC | GC_T48 | BPS | 23.74491773 | 76.25508227 | 17.14707061 |
| GC | GC_T48 | BPS | 22.02553798 | 77.97446202 | 14.91033534 |
| GC | GC_T48 | BPS | 22.6297216 | 77.3702784 | 14.68378914 |
| GC | GC_T48 | BPS | 19.11285833 | 80.88714167 | 11.7654029 |
| GC | GC_T48 | CTRL | 22.95760134 | 77.04239866 | 14.91011585 |
| GC | GC_T48 | CTRL | 20.46140698 | 79.53859302 | 14.54699718 |
| GC | GC_T48 | CTRL | 15.72365433 | 84.27634567 | 13.31306281 |
| GC | GC_T48 | CTRL | 20.0532903 | 79.9467097 | 16.2347505 |
| GC | GC_T48 | CTRL | 26.33908405 | 73.66091595 | 16.17371652 |
| GC | GC_T48 | CTRL | 23.00753735 | 76.99246265 | 13.13690975 |
| GC | GC_T48 | CTRL | 21.76885448 | 78.23114552 | 15.83678862 |
| GC | GC_T48 | CTRL | 23.52325686 | 76.47674314 | 15.58957383 |
| GC | GC_T48 | CTRL | 22.96127453 | 77.03872547 | 14.26619899 |
| GC | GC_T48 | CTRL | 23.93692461 | 76.06307539 | 14.95363028 |

:onditioned media ; EVs = Extracellular vesicles

| rpin | mature_hairpin_genome | |
|---|---|---|
| Unmapped | Mapped (with MQ>0) | Unmapped |
| 73.5134142 | 99.03338651 | 0.966613494 |
| 62.24902021 | 98.86348335 | 1.136516647 |
| 63.00442909 | 99.06195352 | 0.938046478 |
| 58.06610176 | 99.09757884 | 0.902421162 |
| 56.58395094 | 99.24339205 | 0.756607951 |
| 57.22638902 | 99.10759332 | 0.892406681 |
| 50.07271485 | 99.24235214 | 0.757647857 |
| 54.90148985 | 99.37159638 | 0.628403624 |
| 50.48545503 | 99.1067037 | 0.893296303 |
| 48.29793096 | 99.08246803 | 0.917531968 |
| 99.16435169 | 87.63148271 | 12.36851729 |
| 99.86115106 | 84.46924838 | 15.53075162 |
| 99.82959285 | 90.26719204 | 9.732807964 |
| 99.84744895 | 86.36177241 | 13.63822759 |
| 99.84338934 | 84.27002115 | 15.72997885 |
| 99.89018313 | 93.301963 | 6.698036998 |
| 99.70389608 | 81.73787837 | 18.26212163 |
| 99.62114479 | 82.96365021 | 17.03634979 |
| 99.70841909 | 82.22187743 | 17.77812257 |
| 99.72717807 | 95.83682748 | 4.163172519 |
| 99.12134003 | 89.42799023 | 10.57200977 |
| 99.88319192 | 72.09794568 | 27.90205432 |
| 99.92904568 | 76.95267856 | 23.04732144 |
| 99.85185066 | 83.23307759 | 16.76692241 |
| 99.7616575 | 88.74652497 | 11.25347503 |
| 99.86017962 | 87.14430182 | 12.85569818 |
| 99.29655105 | 84.98165758 | 15.01834242 |
| 99.54473321 | 86.05680536 | 13.94319464 |
| 99.80925923 | 79.31372648 | 20.68627352 |
| 99.52494874 | 85.21750102 | 14.78249898 |
| 99.18205576 | 88.81815103 | 11.18184897 |
| 99.85991543 | 84.55266906 | 15.44733094 |
| 99.87035424 | 82.7193995 | 17.2806005 |
| 99.80994564 | 86.31511154 | 13.68488846 |
| 99.76386037 | 85.58229072 | 14.41770928 |
| 99.83862859 | 86.96930208 | 13.03069792 |
| 99.75218834 | 85.03966794 | 14.96033206 |
| 99.40843418 | 86.56450682 | 13.43549318 |
| 99.63741955 | 83.38896366 | 16.61103634 |
| 99.83471737 | 76.73712173 | 23.26287827 |
| 88.37229098 | 99.83015431 | 0.169845693 |
| 90.8214504 | 99.66843748 | 0.331562525 |
| 88.84761882 | 99.73949807 | 0.260501928 |
| 84.03866609 | 99.72548232 | 0.274517677 |
| 87.77306309 | 99.72195392 | 0.27804608 |
| 83.16988683 | 99.73318092 | 0.26681908 |
| 88.74570371 | 99.79432704 | 0.205672957 |
| 88.79917288 | 99.79921824 | 0.200781759 |

| | | |
|---|---|---|
| 91.00953481 | 99.68331976 | 0.31668024 |
| 89.73835196 | 99.7434901 | 0.256509901 |
| 83.85533286 | 99.60619373 | 0.393806275 |
| 85.66618393 | 99.76407698 | 0.235923016 |
| 88.18953053 | 99.61042116 | 0.389578842 |
| 86.75340408 | 97.52941834 | 2.470581661 |
| 85.75453318 | 99.79260264 | 0.207397361 |
| 83.46677807 | 99.46089318 | 0.539106821 |
| 84.20575173 | 99.51760734 | 0.48239266 |
| 86.59299525 | 99.52969063 | 0.470309371 |
| 84.75491085 | 99.5680561 | 0.431943905 |
| 89.05475382 | 96.63019713 | 3.369802867 |
| 85.052339 | 99.51919789 | 0.480802109 |
| 84.82348661 | 99.61894157 | 0.381058433 |
| 84.05688595 | 99.74075619 | 0.259243812 |
| 86.51788478 | 99.5748265 | 0.425173495 |
| 82.4787918 | 99.60385076 | 0.396149237 |
| 85.05346287 | 99.76694252 | 0.233057483 |
| 82.85292939 | 99.41825522 | 0.581744782 |
| 85.08966466 | 99.55519153 | 0.444808472 |
| 85.31621086 | 99.41827475 | 0.581725246 |
| 88.2345971 | 99.5821625 | 0.417837498 |
| 85.08988415 | 99.6179577 | 0.382042302 |
| 85.45300282 | 99.69553021 | 0.304469794 |
| 86.68693719 | 99.58089785 | 0.419102153 |
| 83.7652495 | 99.5903693 | 0.409630695 |
| 83.82628348 | 99.54066004 | 0.459339958 |
| 86.86309025 | 99.77978929 | 0.220210709 |
| 84.16321138 | 99.39898783 | 0.601012168 |
| 84.41042617 | 99.5431627 | 0.456837299 |
| 85.73380101 | 99.49636805 | 0.503631947 |
| 85.04636972 | 99.59802466 | 0.401975336 |
