## Supplemental Table7 for "Characterization of ovine follicular fluid and granulosa cell-derived extracellular vesicles and their miRNA cargo following in vitro exposure to bisphenols A and S"

| **Comparison** | **0h GC vs FF EVs** | | | **0h GC vs 48h GC C** | |
|---|---|---|---|---|---|
| Sequence annotation | Increased in 0h GC | Increased in FF EVs | Total | Increased in 0h GC | Increased in 48h GC |
| Ovine annotated | 27 | 52 | 79 | 24 | 12 |
| miRMachine | 42 | 38 | 80 | 34 | 33 |
| RumimiR | 47 | 66 | 113 | 42 | 48 |
| miRDeep2 | 13 | 27 | 40 | 11 | 16 |
| Total | 129 | 183 | 312 | 111 | 109 |

CTRL = Control ; FF = Follicular fluid ; GC = Granulosa cells ; EVs = Extracellular vesicles

| TRL |
| --- |
| Total |
| 36 |
| 67 |
| 90 |
| 27 |
| 220 |
