## Supplemental Table8 for "Characterization of ovine follicular fluid and granulosa cell-derived extracellular vesicles and their miRNA cargo following in vitro exposure to bisphenols A and S"

| miR identification or sequence | log2 Fold Change (lfc) | lfc standard error | padj |
|---|---|---|---|
| oar-02648a | 0.23 | 1.12E-01 | 4.77E-02 |
| gaaacuggaacgaacuuuuugg | 0.24 | 9.83E-02 | 2.22E-02 |
| oar-10a | 0.30 | 9.39E-02 | 2.38E-03 |
| oar-04969 | 0.38 | 7.47E-02 | 8.19E-07 |
| oar-mir-221 | 0.43 | 7.57E-02 | 3.57E-08 |
| oar-03613 | 0.43 | 1.41E-01 | 3.30E-03 |
| oar-02598 | 0.45 | 1.90E-01 | 2.26E-02 |
| oar-04560b | 0.46 | 1.26E-01 | 3.62E-04 |
| oar-mir-17 | 0.49 | 6.77E-02 | 7.82E-13 |
| oar-17a | 0.58 | 7.14E-02 | 1.48E-15 |
| oar-mir-191 | 0.59 | 1.02E-01 | 1.73E-08 |
| oar-04560a | 0.61 | 1.15E-01 | 2.04E-07 |
| oar-02833 | 0.62 | 2.08E-01 | 4.39E-03 |
| oar-mir-23a | 0.65 | 1.12E-01 | 1.60E-08 |
| oar-425 | 0.65 | 8.28E-02 | 1.20E-14 |
| oar-34c | 0.65 | 2.95E-01 | 3.70E-02 |
| oar-04036 | 0.69 | 1.67E-01 | 6.25E-05 |
| oar-03119 | 0.70 | 1.81E-01 | 1.68E-04 |
| oar-148a | 0.73 | 6.89E-02 | 9.84E-26 |
| oar-04777 | 0.73 | 1.55E-01 | 4.47E-06 |
| oar-01605 | 0.73 | 1.40E-01 | 3.35E-07 |
| oar-04203 | 0.73 | 1.65E-01 | 1.62E-05 |
| oar-let-7b | 0.79 | 8.26E-02 | 2.07E-21 |
| oar-129b | 0.80 | 3.30E-01 | 2.10E-02 |
| oar-mir-125b | 0.82 | 1.10E-01 | 1.50E-13 |
| oar-130e | 0.86 | 2.14E-01 | 1.11E-04 |
| oar-181a | 0.88 | 7.77E-02 | 7.90E-29 |
| oar-34a | 0.89 | 1.66E-01 | 1.89E-07 |
| oar-04145 | 0.90 | 2.36E-01 | 2.37E-04 |
| oar-04334 | 0.90 | 1.67E-01 | 1.63E-07 |
| oar-02050 | 0.93 | 3.84E-01 | 2.17E-02 |
| oar-17c | 0.93 | 6.12E-02 | 2.62E-51 |
| oar-mir-200c | 0.98 | 1.21E-01 | 1.84E-15 |
| oar-15b | 1.00 | 1.07E-01 | 3.13E-20 |
| oar-00815b | 1.01 | 2.11E-01 | 3.33E-06 |
| oar-03347 | 1.03 | 2.08E-01 | 1.44E-06 |
| cacauggaguugcuguuaca | 1.03 | 4.70E-01 | 3.77E-02 |
| oar-101a | 1.04 | 2.95E-01 | 6.40E-04 |
| oar-221 | 1.05 | 1.28E-01 | 5.11E-16 |
| oar-15a | 1.08 | 8.55E-02 | 3.89E-36 |
| oar-132a | 1.16 | 1.68E-01 | 1.27E-11 |
| oar-15c | 1.21 | 1.08E-01 | 1.64E-28 |
| oar-04754 | 1.27 | 3.30E-01 | 2.09E-04 |
| oar-let-7c | 1.28 | 9.66E-02 | 2.29E-39 |
| oar-196b | 1.31 | 2.48E-01 | 2.59E-07 |
| oar-05084b | 1.32 | 3.83E-01 | 9.02E-04 |
| oar-03457 | 1.35 | 3.02E-01 | 1.32E-05 |
| oar-455 | 1.36 | 1.11E-01 | 7.66E-34 |
| oar-03542 | 1.38 | 3.75E-01 | 3.71E-04 |

| | | | |
|---|---|---|---|
| oar-03746 | 1.43 | 2.98E-01 | 3.03E-06 |
| oar-mir-27a | 1.43 | 1.39E-01 | 2.03E-24 |
| oar-05022a | 1.53 | 3.25E-01 | 4.85E-06 |
| oar-00655 | 1.53 | 1.73E-01 | 3.89E-18 |
| oar-mir-106a | 1.56 | 1.11E-01 | 3.10E-44 |
| oar-04910 | 1.56 | 2.18E-01 | 2.17E-12 |
| oar-219 | 1.57 | 3.64E-01 | 2.85E-05 |
| oar-34e | 1.64 | 3.00E-01 | 9.88E-08 |
| oar-mir-181a-2 | 1.68 | 9.13E-02 | 5.95E-75 |
| oar-03461 | 1.69 | 1.93E-01 | 8.06E-18 |
| oar-204b | 1.70 | 3.68E-01 | 7.13E-06 |
| oar-00342 | 1.71 | 4.93E-02 | 1.72E-261 |
| oar-00496 | 1.74 | 3.92E-01 | 1.67E-05 |
| oar-01510 | 1.87 | 1.77E-01 | 1.32E-25 |
| oar-03834 | 1.90 | 1.24E-01 | 4.49E-52 |
| oar-04836 | 2.01 | 2.24E-01 | 9.24E-19 |
| oar-05078 | 2.04 | 7.81E-02 | 5.48E-149 |
| oar-02832a | 2.08 | 1.22E-01 | 1.52E-64 |
| oar-193a | 2.11 | 1.82E-01 | 1.88E-30 |
| oar-let-7i | 2.12 | 7.49E-02 | 9.27E-176 |
| oar-146b | 2.13 | 2.04E-01 | 5.73E-25 |
| oar-31 | 2.14 | 1.32E-01 | 3.96E-58 |
| oar-mir-374b | 2.14 | 1.56E-01 | 2.41E-42 |
| oar-01288 | 2.31 | 1.71E-01 | 1.71E-40 |
| oar-02778 | 2.37 | 1.13E-01 | 7.90E-97 |
| oar-01132 | 2.39 | 1.89E-01 | 6.03E-36 |
| oar-04620 | 2.53 | 9.87E-02 | 4.60E-144 |
| oar-mir-194 | 2.66 | 7.68E-02 | 7.45E-262 |
| oar-05062 | 2.68 | 1.24E-01 | 1.29E-102 |
| oar-mir-21 | 2.71 | 1.91E-01 | 7.54E-45 |
| oar-mir-23b | 2.75 | 9.33E-02 | 3.36E-190 |
| oar-00238 | 2.86 | 3.55E-01 | 2.63E-15 |
| oar-01280 | 2.88 | 1.42E-01 | 2.59E-90 |
| oar-mir-26a | 2.89 | 9.21E-02 | 1.00E-214 |
| oar-05139 | 2.92 | 3.31E-01 | 3.13E-18 |
| oar-10b | 2.93 | 1.11E-01 | 2.35E-152 |
| oar-181d | 2.97 | 1.93E-01 | 1.34E-52 |
| oar-03468 | 2.98 | 2.54E-01 | 4.54E-31 |
| oar-00070 | 3.01 | 8.47E-02 | 7.71E-275 |
| oar-mir-362 | 3.04 | 8.96E-02 | 2.57E-251 |
| oar-147 | 3.11 | 2.17E-01 | 8.51E-46 |
| oar-mir-374a | 3.12 | 2.43E-01 | 4.26E-37 |
| oar-01301 | 3.18 | 1.97E-01 | 8.40E-58 |
| oar-mir-26b | 3.23 | 9.33E-02 | 2.21E-261 |
| oar-17d | 3.30 | 1.11E-01 | 8.10E-194 |
| oar-9a | 3.44 | 2.14E-01 | 2.66E-57 |
| oar-92b | 3.48 | 2.05E-01 | 1.49E-63 |
| oar-mir-103 | 3.57 | 7.25E-02 | 0.00E+00 |
| oar-142 | 3.64 | 1.05E-01 | 1.54E-263 |
| oar-let-7d | 3.72 | 7.88E-02 | 0.00E+00 |

| | | | |
|---|---|---|---|
| oar-03172 | 3.80 | 3.34E-01 | 1.97E-29 |
| ugagguaggagguuguauaguu | 3.84 | 8.61E-02 | 0.00E+00 |
| oar-32 | 3.94 | 2.05E-01 | 2.10E-81 |
| oar-7b | 3.95 | 1.64E-01 | 3.71E-126 |
| oar-let-7a | 4.08 | 7.13E-02 | 0.00E+00 |
| oar-33 | 4.14 | 2.26E-01 | 3.21E-74 |
| oar-05057 | 4.18 | 1.01E-01 | 0.00E+00 |
| aauugagguguaguuagcaaaca | 4.46 | 8.02E-01 | 5.79E-08 |
| oar-mir-107 | 4.54 | 2.21E-01 | 1.01E-92 |
| oar-155 | 4.54 | 2.47E-01 | 8.20E-75 |
| oar-let-7f | 4.55 | 1.01E-01 | 0.00E+00 |
| oar-mir-150 | 4.59 | 1.27E-01 | 2.23E-282 |
| oar-146a | 4.65 | 1.17E-01 | 0.00E+00 |
| oar-130a | 4.68 | 1.87E-01 | 7.21E-137 |
| ccucacugaugagcaccu | 4.71 | 4.74E-01 | 1.17E-22 |
| oar-7a | 4.91 | 2.74E-01 | 1.04E-70 |
| oar-190b | 4.92 | 3.98E-01 | 1.97E-34 |
| oar-7c | 4.94 | 3.32E-01 | 2.07E-49 |
| oar-00430 | 5.05 | 3.66E-01 | 1.47E-42 |
| oar-9c | 5.89 | 5.57E-01 | 1.38E-25 |
| oar-223 | 6.05 | 1.59E-01 | 0.00E+00 |
| oar-mir-181a-1 | 6.45 | 6.02E-01 | 3.31E-26 |
| aaagaaguuuguucagguuuuu | 6.51 | 2.06E+00 | 2.40E-03 |
| caagcacccccauuuugagca | 7.24 | 1.31E+00 | 7.41E-08 |
| gcccuuugaggguuuggguuu | 8.28 | 7.39E-01 | 1.60E-28 |
| guccuggugucugaguga | 9.67 | 1.43E+00 | 3.70E-11 |
| auugguacuucuuagagugaa | 10.48 | 1.73E+00 | 2.99E-09 |
| auggcgcuggagcgucgggcc | 14.76 | 1.60E+00 | 9.63E-20 |
| ugagguaguaaguuguauuguu | 16.64 | 6.58E-01 | 5.81E-140 |
| gcgcgcgccgcggcugga | 38.05 | 2.86E+00 | 9.99E-40 |

FF = Follicular fluid ; GC = Granulosa cells ; EVs = Extracellular vesicles

| Sequence | 0h GC mean read | FF EV mean read | Annotation |
|---|---|---|---|
| ACACAGGUUUUGCUUCCAUCAC | 86.13 | 73.84 | RumimiR |
| - | 368.91 | 313.91 | MiRDeep2 |
| CCACCCGUAGAACCGACCUUG | 8256.32 | 6694.82 | MiRMachine |
| CAACUGUUUGCAGAGGAAACUGA | 1673.68 | 1291.39 | RumimiR |
| AGCUACAUUGUCUGCUGGGUUU | 12079.76 | 8931.85 | Ovine |
| UCCGAGCCUGGGUCUCCCUCU | 204.95 | 141.79 | RumimiR |
| GACCCUGCAGCCAAAGAAGCUAC | 79.45 | 59.08 | RumimiR |
| UGCGGGAUCUUUAGUUGUGGCG | 6028.51 | 4268.89 | RumimiR |
| CAAAGUGCUUACAGUGCAGGUA | 86855.28 | 61586.08 | Ovine |
| CUAAAGUGCUUAUAGUGCAGGUA | 70618.86 | 47240.97 | MiRMachine |
| CAACGGAAUCCCAAAAGCAGCU | 147060.00 | 96036.33 | Ovine |
| UUGCGGGAUCUUUAGUUGUGG | 1457.02 | 937.29 | RumimiR |
| UAUUCAUUUAUCUCCCAGCCUAC | 183.14 | 120.73 | RumimiR |
| AUCACAUUGCCAGGGAUUUCCA | 108857.38 | 69171.97 | Ovine |
| GAAUGACACGAUCACUCCCGUUGAG | 5151.21 | 3280.59 | MiRMachine |
| UAGGCAGUGUAGUUAGCUGAUUG | 593.21 | 235.57 | MiRMachine |
| AGUCCUGAGUGGUCCCUUCAG | 47.31 | 28.70 | RumimiR |
| ACUCCCGGUAGGCAACGCGCC | 50.68 | 30.59 | RumimiR |
| UCAGUGCAUCACAGAACUUUGU | 6918.51 | 4176.30 | MiRMachine |
| UUGUUGUCAGGAGUCACUGCU | 78.09 | 46.89 | RumimiR |
| GCUUGGCACCUAGUAAGUACUC | 1900.99 | 1137.78 | RumimiR |
| CAAGGAGCUUACAAUCUAGCUG | 1006.39 | 583.83 | RumimiR |
| UGAGGUAGUAGGUUGUGUGGU | 133635.00 | 76193.44 | Ovine |
| UCUUUUUGCGGUCUGGGCUU | 60.81 | 27.55 | MiRMachine |
| UCCCUGAGACCCUAACUUGUG | 34153.62 | 19029.13 | Ovine |
| CAGUGCAAUGAUAUUGUCAAAGC | 140.23 | 76.27 | MiRMachine |
| AACAUUCAUUGCUGUCGGUGGGUU | 1079.80 | 580.83 | MiRMachine |
| UUGGCAGUGUCUUAGCUGGUUGU | 2624.17 | 1401.85 | MiRMachine |
| CAUAGCCAGUUGGGGAAGAAUG | 29.83 | 17.53 | RumimiR |
| ACUCGGCGUGGCGUCGGUCGUG | 921.86 | 502.61 | RumimiR |
| UGCGAGUUCGGUGGGAUUUGCU | 12.04 | 6.50 | RumimiR |
| UUAAGGUGCAUCUAGUGCA | 15547.84 | 8130.59 | MiRMachine |
| UAAUACUGCCGGGUAAUGAUGG | 797.82 | 378.38 | Ovine |
| UAGCAGCACAUAAUGGUUUGU | 19229.10 | 9293.03 | MiRMachine |
| AAAACCCGCAUGAACUUUUUGG | 81.13 | 41.23 | RumimiR |
| GUCCCGGGCUGGAGGAGUCUGC | 273.54 | 126.04 | RumimiR |
| - | 21.89 | 10.51 | MiRDeep2 |
| GUACAGUACUGUGAUAACUGA | 158.59 | 76.76 | MiRMachine |
| AGCUACAUCUGGCUACUGGGUCU | 1140.72 | 540.87 | MiRMachine |
| UAGCAGCACAUCAUGGUUUACA | 127277.37 | 60077.34 | MiRMachine |
| UAACAGUCUACAGCCAUGGUCG | 661.62 | 275.99 | MiRMachine |
| UAGCAGCACAGAAAUGUUGGUA | 10392.72 | 4517.69 | MiRMachine |
| UCCUCUCUCGGUUAGCUCCAUA | 22.04 | 9.39 | RumimiR |
| UGAGGUAGUAGGUUGUAUGGUU | 25602.35 | 10387.53 | Ovine |
| UUAGGUAGUUUCCUGUUGUUG | 82.86 | 32.59 | MiRMachine |
| AAAAGUUCAUUUGGGUUGUUCCU | 19.73 | 8.19 | RumimiR |
| GACCCUGACGGGCGUGGAUUGU | 18.98 | 7.30 | RumimiR |
| GCAGUCCAUGGGCAUAUACA | 1930.41 | 755.24 | MiRMachine |
| GUUACAGUUGUUCAACCAGUUAC | 16.93 | 6.35 | RumimiR |

| | | | |
|---|---|---|---|
| ACCACCUCCCCUGCAAACGUCC | 1293.96 | 466.85 | RumimiR |
| UUCACAGUGGCUAAGUUCCGC | 30089.34 | 10622.91 | Ovine |
| UUUUUGCGAUGUGUUCCUAAUA | 33.93 | 11.08 | RumimiR |
| UAGUGGGGAACCCUUCCAUGAGG | 733.38 | 251.90 | RumimiR |
| AAAAGUGCUUACAGUGCAGGU | 620.11 | 202.93 | Ovine |
| AUCCACCUGGGCAAGGAUUCUGAA | 32.91 | 10.56 | RumimiR |
| GAGAGUUGAGUCUGGACGUCCC | 25.92 | 8.46 | MiRMachine |
| UUGGCAGUGUAUUGUUAGCUGG | 403.62 | 105.40 | MiRMachine |
| AACAUUCAACGCUGUCGGUGAGU | 13662.00 | 4132.17 | Ovine |
| CCUCUGGGCCCUUCCUCCAGC | 260.91 | 84.74 | RumimiR |
| UUCCCUUUGUCAUCCUUUGCCC | 51.25 | 15.46 | MiRMachine |
| AAGGAGCUCACAGUCUAUUGAG | 11010.43 | 3379.92 | RumimiR |
| AAGUUCAUUCGGGUUUUUCCA | 16.39 | 4.76 | RumimiR |
| AUUAUAAAGCAAUGAGACUGAU | 646.34 | 167.81 | RumimiR |
| UUCCCGGCCAACGCACCA | 1089.91 | 287.41 | RumimiR |
| CUUUUCCAAUGAGUCAGUUAU | 96.03 | 22.90 | RumimiR |
| AAUGGCGCCACUAGGGUUGUGC | 10451.67 | 2507.34 | RumimiR |
| CCACAUGGAGUUGCUGUUACAA | 216.36 | 51.03 | RumimiR |
| UGGGUCUUUGCGGGCGAGAUGA | 2641.27 | 637.82 | MiRMachine |
| UGAGGUAGUAGUUUGUGCUGUU | 52828.11 | 12032.36 | Ovine |
| CUGAGAACUGAAUUCCAUAGGCUG | 6626.32 | 1430.74 | MiRMachine |
| AGGCAAGAUGCUGGCAUAGCUG | 686026.67 | 154911.24 | MiRMachine |
| AUAUAAUACAACCUGCUAAGU | 18008.36 | 3999.56 | Ovine |
| CAUGUCCGCGGGUUCCCUAUC | 433.90 | 78.51 | RumimiR |
| UUUCCCGGCCAGUGCACC | 701.67 | 135.13 | RumimiR |
| GGCCCCUGGGCCUAUCCUAGA | 377.92 | 71.69 | RumimiR |
| UUGGCAUUACUGAGCAUCUAG | 1471.77 | 256.35 | RumimiR |
| UGUAACAGCAACUCCAUGUGGA | 4027.78 | 636.57 | Ovine |
| AAAACCCGAACGAGCUCUUUGG | 283.04 | 42.85 | RumimiR |
| UAGCUUAUCAGACUGAUGUUGAC | 238034.44 | 34781.40 | Ovine |
| AUCACAUUGCCAGGGAUU | 112725.42 | 16107.20 | Ovine |
| GUCAGAGCUGGGUUUAUUGCGUG | 19.79 | 2.53 | RumimiR |
| AUCCCACUCCUGACACCA | 2817.36 | 381.02 | RumimiR |
| UUCAAGUAAUCCAGGAUAGGCU | 131348.41 | 17991.82 | Ovine |
| AUAACACAUGUCAUAACGCGGGG | 27.10 | 3.87 | RumimiR |
| GUCCCUGAGACCCUUUAACCUGUG | 49099.56 | 6298.86 | MiRMachine |
| AACAUUCAUUGCUGUCGGUGGGUU | 79.43 | 10.21 | MiRMachine |
| ACUUCUGUCCGUCUUGCUACC | 33.82 | 4.23 | RumimiR |
| CUCGAGGAGCUCACAGUCUAG | 21938.45 | 2721.82 | RumimiR |
| AAUCCUUGGAACCUAGGUGUGAGU | 11584.86 | 1390.12 | Ovine |
| GUGUGCGGAAAUGCUUCUGCU | 83.52 | 9.84 | MiRMachine |
| UUAUAAUACAACCUGAUAAGUG | 2857.13 | 311.43 | Ovine |
| ACGCCCUUCCCCCCCUUCUUCA | 257.83 | 25.13 | RumimiR |
| UUCAAGUAAUUCAGGAUAGGU | 51589.50 | 5431.81 | Ovine |
| CCAAAGUGCUCACAGUGCAGGUA | 821.84 | 85.17 | MiRMachine |
| UCUUUGGUUAUCUAGCUGUAUGA | 76.67 | 6.84 | MiRMachine |
| AAAUUGCACGGUAUCCAUCUGC | 93.09 | 8.43 | MiRMachine |
| AGCAGCAUUGUACAGGGCUAUG | 54849.16 | 4610.55 | Ovine |
| CCCAUAAAGUAGAAAGCACU | 17882.03 | 1349.16 | MiRMachine |
| AGAGGUAGUAGGUUGCAUAG | 56094.39 | 4246.07 | Ovine |

| | | | |
|---|---|---|---|
| UCCCUGUCUUCAAUCCUGUAGU | 21.25 | 1.63 | RumimiR |
| - | 27279.45 | 1902.74 | MiRDeep2 |
| GCAAUUUAGUGUGUGUGAUAU | 200.26 | 12.44 | MiRMachine |
| UGGAAGACUAGUGAUUUUGUUGUU | 1654.60 | 103.60 | MiRMachine |
| UGAGGUAGUAGGUUGUAUAGUU | 394774.46 | 23477.07 | Ovine |
| GCAGUGCCUCGGCAGUGCAGC | 294.06 | 16.55 | MiRMachine |
| GAGAGGUAAAAAAUUGAUUUGAC | 8643.13 | 474.26 | RumimiR |
| - | 21.75 | 1.27 | MiRDeep2 |
| AGCAGCAUUGUACAGGGCUAUC | 4478.11 | 182.23 | Ovine |
| UUAAUGCUAAUCGUGAUAGGGGUU | 700.04 | 27.79 | MiRMachine |
| UGAGGUAGUAGAUUGUAUAGU | 198484.68 | 8448.73 | Ovine |
| UCUCCCAACCCUUGUACCAGUG | 21753.87 | 866.75 | Ovine |
| UUGAGAACUGAAUUCCAUAGGU | 3531.20 | 134.26 | MiRMachine |
| UAGUGCAAUAUUGCUUAUAGGGUU | 302.07 | 11.97 | MiRMachine |
| - | 75.09 | 3.04 | MiRDeep2 |
| GUGGAAGACUAGUGAUUUUGUUGU | 75.87 | 2.69 | MiRMachine |
| UGAUAUGUUUGAUAUAUUAGGUU | 70.38 | 2.53 | MiRMachine |
| CUGGAAGACUAGUGAUUUUGUUGU | 43.26 | 1.53 | MiRMachine |
| GUCCCCGAGUCCCUGGCGUGCAC | 39.52 | 1.13 | RumimiR |
| CUCUUUGGUUAUCUAGCUGUAUG | 22.20 | 0.36 | MiRMachine |
| GUGUCAGUUUGUCAAAUACCCC | 13125.36 | 196.41 | MiRMachine |
| AACAUUCAACGCUGUCGGUGAGU | 24.49 | 0.32 | Ovine |
| - | 14.12 | 0.00 | MiRDeep2 |
| - | 18.99 | 0.00 | MiRDeep2 |
| - | 27.28 | 0.00 | MiRDeep2 |
| - | 75.00 | 0.00 | MiRDeep2 |
| - | 84.59 | 0.00 | MiRDeep2 |
| - | 1328.57 | 6.01 | MiRDeep2 |
| - | 9184.49 | 0.00 | MiRDeep2 |
| - | 412.52 | 0.00 | MiRDeep2 |
