## Supplemental Table9 for "Characterization of ovine follicular fluid and granulosa cell-derived extracellular vesicles and their miRNA cargo following in vitro exposure to bisphenols A and S"

| miR identification or sequence | log2 Fold Change (lfc) | lfc standard error | padj |
|---|---|---|---|
| caggauuaaccagaggacagugu | -44.48 | 3.07E+00 | 6.98E-47 |
| gggcucaguucagcaggag | -36.83 | 3.60E+00 | 5.76E-24 |
| aaaaaccugagcaaacuuuuug | -12.62 | 2.42E+00 | 3.49E-07 |
| ugagguaguaaguuguauugu | -11.51 | 3.66E+00 | 2.52E-03 |
| uggcaccagcacuggcggugg | -11.45 | 2.07E+00 | 6.32E-08 |
| acaaaaaaaaaagcccaacccu | -9.85 | 1.89E+00 | 3.50E-07 |
| auugguacuucuuagaguga | -9.10 | 1.77E+00 | 5.58E-07 |
| agaucuguccugaaaccagcau | -7.17 | 2.63E+00 | 9.02E-03 |
| ccgcaggaaagugacaggaguu | -6.95 | 1.96E+00 | 6.10E-04 |
| aaaaaguuugucuagguuuuuc | -6.56 | 2.15E+00 | 3.36E-03 |
| agcugcagggaccaugcccagu | -6.46 | 2.99E+00 | 4.05E-02 |
| aaaaccagaacgaacuuuuugu | -6.44 | 1.28E+00 | 9.51E-07 |
| aauagcucagaaugucacuucu | -6.21 | 2.03E+00 | 3.30E-03 |
| cggcucugggucuguggggagc | -6.09 | 1.90E+00 | 2.04E-03 |
| oar-mir-410 | -6.07 | 8.92E-01 | 2.43E-11 |
| oar-mir-329b | -6.07 | 5.24E-01 | 2.30E-30 |
| oar-145 | -5.96 | 1.93E-01 | 5.45E-208 |
| uugcugcgggugucaggaaga | -5.89 | 1.89E+00 | 2.77E-03 |
| acgaaucagacacgaguuuga | -5.86 | 2.34E+00 | 1.73E-02 |
| aacagaaagugacuaagaugu | -5.85 | 2.14E+00 | 9.10E-03 |
| oar-mir-485 | -5.48 | 8.10E-01 | 3.25E-11 |
| oar-mir-329a | -5.27 | 1.43E+00 | 3.53E-04 |
| oar-208 | -5.15 | 9.02E-01 | 2.51E-08 |
| oar-mir-379 | -5.09 | 7.25E-01 | 5.37E-12 |
| oar-mir-655 | -4.93 | 6.60E-01 | 2.26E-13 |
| oar-192a | -4.87 | 6.18E-01 | 9.29E-15 |
| oar-mir-382 | -4.87 | 1.05E+00 | 6.31E-06 |
| oar-122 | -4.76 | 3.96E-01 | 1.38E-32 |
| oar-mir-1185 | -4.75 | 1.09E+00 | 2.49E-05 |
| oar-mir-495 | -4.71 | 3.84E-01 | 8.69E-34 |
| oar-mir-376c | -4.56 | 1.81E-01 | 2.55E-138 |
| ugggccuuucaucauggcagcu | -4.55 | 1.45E+00 | 2.53E-03 |
| ucugagaggacagcagacagc | -4.54 | 2.09E+00 | 3.93E-02 |
| oar-mir-143 | -4.53 | 1.29E-01 | 5.17E-271 |
| oar-mir-377 | -4.50 | 1.30E+00 | 8.31E-04 |
| oar-mir-380 | -4.49 | 2.65E-01 | 2.55E-63 |
| oar-mir-494 | -4.47 | 1.66E-01 | 4.13E-158 |
| oar-mir-133 | -4.38 | 3.73E-01 | 4.45E-31 |
| oar-mir-22 | -4.29 | 2.16E-01 | 7.43E-87 |
| oar-mir-323a | -4.16 | 4.71E-01 | 2.94E-18 |
| oar-mir-487a | -4.12 | 5.76E-01 | 2.14E-12 |
| oar-mir-432 | -4.12 | 3.43E-01 | 1.52E-32 |
| oar-mir-376e | -4.11 | 3.16E-01 | 5.43E-38 |
| oar-mir-3958 | -4.07 | 3.53E-01 | 3.63E-30 |
| oar-mir-381 | -4.06 | 6.90E-01 | 9.40E-09 |
| aaacccgaaugaacuucuuugc | -4.05 | 1.48E+00 | 9.04E-03 |
| oar-mir-376a | -4.02 | 7.03E-01 | 2.42E-08 |
| oar-mir-369 | -4.00 | 3.60E-01 | 4.91E-28 |
| oar-mir-487b | -3.96 | 2.87E-01 | 1.59E-42 |

| | | | |
|---|---|---|---|
| oar-214 | -3.93 | 2.56E-01 | 1.34E-52 |
| oar-mir-154a | -3.91 | 1.04E+00 | 2.85E-04 |
| oar-mir-376d | -3.87 | 3.54E-01 | 3.75E-27 |
| oar-mir-665 | -3.77 | 2.83E-01 | 9.76E-40 |
| oar-mir-543 | -3.77 | 5.16E-01 | 7.59E-13 |
| oar-mir-409 | -3.76 | 2.53E-01 | 2.61E-49 |
| oar-mir-758 | -3.49 | 1.42E+00 | 1.97E-02 |
| oar-mir-433 | -3.38 | 5.33E-01 | 5.10E-10 |
| oar-mir-493 | -3.25 | 3.48E-01 | 2.60E-20 |
| oar-mir-199a | -3.25 | 1.66E-01 | 3.85E-84 |
| aauguacuuguggaguuggaga | -3.16 | 1.10E+00 | 5.92E-03 |
| oar-mir-323c | -3.06 | 6.25E-01 | 1.85E-06 |
| oar-mir-496 | -3.00 | 1.12E+00 | 1.01E-02 |
| oar-mir-127 | -2.90 | 1.48E-01 | 2.44E-84 |
| oar-mir-376b | -2.76 | 3.78E-01 | 6.49E-13 |
| oar-04762 | -2.74 | 1.39E-01 | 8.55E-86 |
| oar-mir-411a | -2.72 | 4.30E-01 | 6.35E-10 |
| oar-29b | -2.69 | 6.46E-01 | 5.53E-05 |
| oar-mir-10b | -2.64 | 1.24E-01 | 3.06E-100 |
| oar-00466c | -2.62 | 4.30E-01 | 2.37E-09 |
| oar-04283 | -2.61 | 9.31E-01 | 7.23E-03 |
| oar-29a | -2.57 | 1.95E-01 | 7.37E-39 |
| oar-04427 | -2.55 | 3.84E-01 | 7.73E-11 |
| oar-126 | -2.41 | 1.56E-01 | 2.79E-53 |
| oar-04920 | -2.25 | 2.70E-01 | 2.45E-16 |
| oar-05025 | -2.19 | 2.12E-01 | 1.48E-24 |
| oar-04995a | -2.16 | 7.71E-01 | 7.36E-03 |
| oar-mir-370 | -1.98 | 6.97E-01 | 6.50E-03 |
| oar-96b | -1.98 | 5.43E-01 | 4.33E-04 |
| oar-02468 | -1.94 | 1.46E-01 | 2.14E-39 |
| oar-mir-10a | -1.89 | 1.64E-01 | 5.31E-30 |
| oar-04592 | -1.86 | 2.13E-01 | 8.13E-18 |
| oar-04349 | -1.85 | 3.51E-01 | 2.55E-07 |
| oar-459 | -1.83 | 5.09E-01 | 4.98E-04 |
| oar-05375 | -1.81 | 2.45E-01 | 4.78E-13 |
| oar-34b | -1.80 | 3.11E-01 | 1.52E-08 |
| oar-02825 | -1.75 | 3.65E-01 | 3.01E-06 |
| oar-130c | -1.68 | 1.48E-01 | 2.99E-29 |
| oar-192b | -1.65 | 1.26E-01 | 9.45E-39 |
| caauaugaucaaaugugccaga | -1.63 | 6.52E-01 | 1.74E-02 |
| oar-490 | -1.62 | 1.42E-01 | 2.27E-29 |
| oar-01446 | -1.62 | 1.14E-01 | 4.53E-45 |
| oar-02621 | -1.61 | 1.64E-01 | 3.82E-22 |
| oar-05052 | -1.61 | 1.23E-01 | 2.14E-38 |
| oar-03841 | -1.57 | 1.83E-01 | 2.12E-17 |
| oar-04987 | -1.57 | 1.78E-01 | 4.34E-18 |
| oar-1b | -1.51 | 2.32E-01 | 1.58E-10 |
| oar-05020 | -1.51 | 1.01E-01 | 2.53E-49 |
| oar-mir-30b | -1.48 | 2.81E-01 | 2.77E-07 |
| oar-02032 | -1.48 | 3.79E-01 | 1.59E-04 |

| | | | |
|---|---|---|---|
| oar-02195 | -1.42 | 3.18E-01 | 1.32E-05 |
| oar-mir-29b-1a | -1.42 | 4.57E-01 | 2.89E-03 |
| uacuccagagggugucauuca | -1.40 | 4.59E-01 | 3.44E-03 |
| oar-202 | -1.39 | 2.62E-01 | 2.48E-07 |
| oar-mir-29b-1b | -1.38 | 1.73E-01 | 4.94E-15 |
| oar-00466a | -1.35 | 9.81E-02 | 2.79E-42 |
| oar-00573 | -1.35 | 1.30E-01 | 9.24E-25 |
| oar-00673a | -1.35 | 1.46E-01 | 1.24E-19 |
| oar-03709 | -1.34 | 1.49E-01 | 1.06E-18 |
| oar-130d | -1.33 | 8.62E-02 | 6.41E-53 |
| oar-00826 | -1.32 | 1.37E-01 | 2.89E-21 |
| aaaauccgaacgaacuuuuugg | -1.32 | 2.57E-01 | 5.70E-07 |
| oar-mir-29a | -1.30 | 9.58E-02 | 3.10E-41 |
| oar-204a | -1.30 | 1.25E-01 | 1.41E-24 |
| oar-148b | -1.27 | 9.94E-02 | 5.60E-37 |
| oar-01933 | -1.23 | 6.27E-02 | 1.01E-84 |
| oar-03236 | -1.23 | 1.28E-01 | 2.22E-21 |
| oar-04984 | -1.21 | 2.60E-01 | 5.49E-06 |
| oar-01572 | -1.16 | 2.34E-01 | 1.27E-06 |
| oar-02698 | -1.16 | 1.92E-01 | 3.58E-09 |
| oar-04392 | -1.15 | 5.10E-01 | 3.23E-02 |
| oar-00176a | -1.15 | 6.71E-02 | 8.42E-65 |
| oar-193c | -1.15 | 1.01E-01 | 3.88E-29 |
| oar-00311 | -1.14 | 1.10E-01 | 1.74E-24 |
| oar-03664 | -1.12 | 2.65E-01 | 3.81E-05 |
| oar-02505b | -1.10 | 3.79E-01 | 5.60E-03 |
| oar-03097 | -1.09 | 1.17E-01 | 3.43E-20 |
| oar-mir-25 | -1.07 | 7.84E-02 | 6.53E-42 |
| oar-04478 | -1.06 | 2.57E-01 | 5.80E-05 |
| oar-mir-200a | -1.05 | 1.30E-01 | 1.57E-15 |
| oar-01877 | -1.05 | 1.47E-01 | 2.79E-12 |
| oar-06127b | -1.02 | 2.57E-01 | 1.18E-04 |
| oar-mir-200b | -1.01 | 1.33E-01 | 9.98E-14 |
| oar-04641 | -1.00 | 2.74E-01 | 4.01E-04 |
| oar-00945 | -0.98 | 2.05E-01 | 3.03E-06 |
| oar-04915 | -0.97 | 5.36E-02 | 1.02E-72 |
| oar-101b | -0.97 | 2.50E-01 | 1.68E-04 |
| oar-01307 | -0.96 | 3.70E-01 | 1.30E-02 |
| oar-02757 | -0.95 | 2.30E-01 | 6.37E-05 |
| oar-mir-19b | -0.94 | 2.64E-01 | 5.81E-04 |
| oar-mir-218a | -0.92 | 2.21E-01 | 5.83E-05 |
| oar-04949 | -0.91 | 8.51E-02 | 2.96E-26 |
| oar-05004 | -0.91 | 1.71E-01 | 2.08E-07 |
| oar-04913 | -0.90 | 1.15E-01 | 1.31E-14 |
| uauugcacuugucccggccugu | -0.87 | 1.15E-01 | 7.43E-14 |
| oar-19 | -0.87 | 2.58E-01 | 1.12E-03 |
| oar-04220 | -0.86 | 2.39E-01 | 5.50E-04 |
| acuggacuuggagucagaaggc | -0.84 | 1.21E-01 | 8.13E-12 |
| oar-00139b | -0.83 | 1.52E-01 | 9.76E-08 |
| oar-30a | -0.83 | 1.44E-01 | 1.74E-08 |

| | | | |
|---|---|---|---|
| oar-mir-148a | -0.83 | 1.30E-01 | 4.51E-10 |
| oar-216 | -0.81 | 2.35E-01 | 8.28E-04 |
| oar-01558 | -0.81 | 3.08E-01 | 1.19E-02 |
| oar-00061 | -0.80 | 1.70E-01 | 4.45E-06 |
| oar-139 | -0.80 | 1.04E-01 | 2.60E-14 |
| oar-04130 | -0.78 | 1.13E-01 | 1.13E-11 |
| oar-02031 | -0.76 | 1.03E-01 | 4.03E-13 |
| oar-203 | -0.75 | 2.09E-01 | 5.16E-04 |
| oar-193b | -0.74 | 1.14E-01 | 2.43E-10 |
| oar-01454 | -0.71 | 2.70E-01 | 1.27E-02 |
| oar-01635 | -0.70 | 1.39E-01 | 8.78E-07 |
| oar-1388 | -0.70 | 1.39E-01 | 1.06E-06 |
| oar-8 | -0.69 | 1.80E-01 | 2.10E-04 |
| oar-02126 | -0.69 | 2.52E-01 | 9.25E-03 |
| oar-01269 | -0.68 | 1.23E-01 | 7.71E-08 |
| oar-03788 | -0.67 | 6.91E-02 | 1.10E-21 |
| oar-17b | -0.67 | 6.82E-02 | 3.75E-22 |
| oar-03867 | -0.67 | 1.27E-01 | 3.08E-07 |
| oar-03854 | -0.65 | 1.32E-01 | 1.55E-06 |
| oar-03094 | -0.64 | 7.14E-02 | 9.55E-19 |
| oar-27 | -0.63 | 7.69E-02 | 9.51E-16 |
| oar-04905 | -0.57 | 7.69E-02 | 2.11E-13 |
| oar-04983a | -0.57 | 2.26E-01 | 1.55E-02 |
| oar-00160 | -0.49 | 1.16E-01 | 3.43E-05 |
| oar-190a | -0.48 | 9.46E-02 | 8.19E-07 |
| oar-24a | -0.44 | 8.16E-02 | 1.32E-07 |
| oar-01889 | -0.41 | 1.48E-01 | 7.67E-03 |
| oar-10c | -0.41 | 8.29E-02 | 1.88E-06 |
| oar-138b | -0.40 | 8.29E-02 | 2.14E-06 |
| oar-128a | -0.40 | 6.12E-02 | 1.16E-10 |
| oar-181c | -0.40 | 1.78E-01 | 3.54E-02 |
| oar-04974 | -0.28 | 6.60E-02 | 3.80E-05 |
| oar-138a | -0.28 | 1.29E-01 | 4.25E-02 |
| uggagagaaaggcaguuccuga | -0.20 | 9.06E-02 | 3.41E-02 |

FF = Follicular fluid ; GC = Granulosa cells ; EVs = Extracellular vesicles

| Sequence | 0h GC mean read | FF EV mean read | Annotation |
|---|---|---|---|
| - | 0.00 | 22.38 | MiRDeep2 |
| - | 0.00 | 14.58 | MiRDeep2 |
| - | 1.85 | 25.01 | MiRDeep2 |
| - | 0.00 | 748.50 | MiRDeep2 |
| - | 2.28 | 50.43 | MiRDeep2 |
| - | 10.65 | 174.54 | MiRDeep2 |
| - | 0.00 | 79.72 | MiRDeep2 |
| - | 0.00 | 34.79 | MiRDeep2 |
| - | 7.31 | 46.29 | MiRDeep2 |
| - | 3.39 | 16.60 | MiRDeep2 |
| - | 2.42 | 10.71 | MiRDeep2 |
| - | 0.00 | 23.04 | MiRDeep2 |
| - | 0.00 | 18.15 | MiRDeep2 |
| - | 0.00 | 12.85 | MiRDeep2 |
| AAUAUAACACAGAUGGCCUGU | 1.07 | 57.13 | Ovine |
| AACACACCUGGUUAACCUCUUU | 5.18 | 320.24 | Ovine |
| GGUCCAGUUUUCCCAGGAAUCCC | 672.38 | 38292.65 | MiRMachine |
| - | 6.82 | 38.88 | MiRDeep2 |
| - | 1.26 | 28.06 | MiRDeep2 |
| - | 1.18 | 5.59 | MiRDeep2 |
| GUCAUACACGGCUCUCCUCUCU | 2.16 | 58.76 | Ovine |
| AACACACCUGGUUAACCUUUUU | 0.34 | 14.84 | Ovine |
| UAUAAGACGAGCAAAAAGC | 0.12 | 10.90 | MiRMachine |
| UAUGUAACAUGGUCCACUAACU | 0.94 | 28.40 | Ovine |
| AUAAUACAUGGUUAACCUCUCU | 3.35 | 95.14 | Ovine |
| AUGACCUAUGAAUUGACAGA | 5.23 | 109.79 | MiRMachine |
| AAUCAUUCACGGACAACACUU | 0.89 | 22.67 | Ovine |
| GUGGAGUGUGACAAUGGUGUU | 6.94 | 216.36 | MiRMachine |
| AUAUACAGAGGGAGACUCUUAU | 0.14 | 8.49 | Ovine |
| AAACAAACAUGGUGCACUUCUU | 8.58 | 216.85 | Ovine |
| AACAUAGAGGAAAUUCCACGU | 73.80 | 1668.52 | Ovine |
| - | 0.00 | 8.08 | MiRDeep2 |
| - | 3.09 | 13.87 | MiRDeep2 |
| UGAGAUGAAGCACUGUAGCUC | 1276.90 | 28999.47 | Ovine |
| AUCACACAAAGGCAACUUUCGU | 0.54 | 11.61 | Ovine |
| UAUGUAAUGUGGUCCACGUCU | 34.54 | 764.41 | Ovine |
| UGAAACAUACACGGGAAACCUCU | 66.18 | 1408.36 | Ovine |
| UUGGUCCCCUUCAACCAGCUGU | 13.95 | 243.69 | Ovine |
| AAGCUGCCAGUUGAAGAACUG | 2357.43 | 44673.09 | Ovine |
| CACAUUACACGGUCGACCUCU | 3.76 | 65.72 | Ovine |
| AUCAUACAGGGACAUCCAGUUU | 3.04 | 46.14 | Ovine |
| UCUUGGAGUAGGUCAUUGGGUGG | 14.48 | 231.58 | Ovine |
| AACAUAGAGGAAAAUCCACAUU | 20.78 | 336.86 | Ovine |
| AGAUAUUGCACGGUUGAUCUCU | 13.83 | 228.14 | Ovine |
| AUAUACAAGGGCAAGCUCUCU | 2.58 | 31.72 | Ovine |
| - | 8.18 | 56.24 | MiRDeep2 |
| AUCAUAGAGGAAAAUCCACGU | 1.66 | 20.13 | Ovine |
| AAUAAUACAUGGUUGAUCUUU | 16.02 | 225.51 | Ovine |
| AAUCGUACAGGGUCAUCCACUU | 28.20 | 424.77 | Ovine |

| | | | |
|---|---|---|---|
| ACAGCAGGCACAGACAGGCAGU | 560.24 | 5725.66 | MiRMachine |
| AAUCAUACACGGUUCACCUAUU | 0.61 | 5.72 | Ovine |
| AUCAUAGAGGAAAAUCCACAU | 8.30 | 108.43 | Ovine |
| ACCAGUAGGCCGAGGCCCCUCA | 19.37 | 236.23 | Ovine |
| AAACAUUCGCGGUGCACUUCUUU | 3.78 | 44.12 | Ovine |
| CGAAUGUUGCUCGGUGAACCCCU | 31.33 | 377.65 | Ovine |
| UUUGUGACCUGGUCCACUAA | 1.03 | 6.35 | Ovine |
| AUCAUGAUGGGCUCCUCGGUGU | 6.49 | 49.44 | Ovine |
| UGAAGGUCUACUGUGUGCCAGG | 11.41 | 103.05 | Ovine |
| ACAGUAGUCUGCACAUUGGUU | 6132.01 | 46534.87 | Ovine |
| - | 3.68 | 14.72 | MiRDeep2 |
| CACAAUACACGGUCGGCCUCU | 3.68 | 26.11 | Ovine |
| UGAGUAUUACAUGGCCAAUCU | 0.85 | 5.54 | Ovine |
| AUCGGAUCCGUCUGAGCUUGGCU | 167.37 | 1213.43 | Ovine |
| AUCAUAGAGGAAAAUCCAUGU | 12.47 | 74.90 | Ovine |
| UCCUGUACUGAGCUGCCCCGA | 344.26 | 2344.50 | RumimiR |
| UAUGUAACACGGUCCACUAAC | 7.14 | 36.36 | Ovine |
| CUAGCACCAUUUGAAAUCGGUU | 2.30 | 15.16 | MiRMachine |
| ACCCUGUAGAACCGAAUUUGUG | 8115.56 | 50102.44 | Ovine |
| AAAAAGUUCCUUUGGGUUUU | 5.24 | 26.30 | RumimiR |
| UCAUCCCCGCCACAGACUCUGC | 1.19 | 5.87 | RumimiR |
| CUAGCACCAUUUGAAAUCGGUU | 343.38 | 1917.59 | MiRMachine |
| GGAAACCCGAACGAACUUUUG | 7.86 | 43.07 | RumimiR |
| CAUUAUUACUUUUGGUACGCG | 356.22 | 1781.36 | MiRMachine |
| UCUGGAGGGGCAGAAGGAGAAG | 47.72 | 222.53 | RumimiR |
| UUUUUGCAAUAUGUUCCUGAA | 330.51 | 1242.67 | RumimiR |
| GUACUGUGCCACGGAUGGGUAG | 2.39 | 7.26 | RumimiR |
| GCCUGCUGGGGUGGAACCUGGUCU | 4.33 | 12.71 | Ovine |
| UUUGGCACUAGCACAUUUUUGC | 4.66 | 15.79 | MiRMachine |
| GAAAGGUUCAUUUGGGUUUUU | 137.25 | 518.92 | RumimiR |
| UACCCUGUAGAUCCGAAUUUG | 595.66 | 2020.22 | Ovine |
| GUCCCUGUCCUCCAGGAGCUCAC | 6670.07 | 23566.88 | RumimiR |
| GUAGAGAAGCGCUGGGGGAAAG | 13.67 | 44.95 | RumimiR |
| AUCAGUAACAAAGAUUCAUCCUU | 5.86 | 17.61 | MiRMachine |
| UAAAAAGGUCUUUCGAGUUUUU | 44.02 | 150.16 | RumimiR |
| CAAUCACUAGUUCCACUGCCAU | 169.05 | 390.74 | MiRMachine |
| AAAAAACCGAGUGAACUUUUUG | 6.78 | 19.33 | RumimiR |
| GCAGUGCAAUGUUAAAAGGGC | 3370.65 | 10719.99 | MiRMachine |
| CUGACCUAUGAAUUGACAGCC | 462.43 | 1421.99 | MiRMachine |
| - | 5.54 | 12.61 | MiRDeep2 |
| ACCAUGGAUCUCCAGGUGGG | 86.92 | 260.81 | MiRMachine |
| GUCAAGAGCAAUAACGAAAAAU | 1454.35 | 4437.97 | RumimiR |
| CUGAGGGGCAGAGAGCGAGACUU | 16537.07 | 48451.25 | RumimiR |
| AAACAUCUGGUUGGUUGAGAGA | 83.30 | 256.69 | RumimiR |
| UCGCAACACAGUUCCAACCGAGA | 57.39 | 173.35 | RumimiR |
| GAUUGGCACGUCUUGGAAUG | 154.50 | 453.91 | RumimiR |
| AUGGAAUGUAAAGAAGUAUGUA | 28.87 | 80.71 | MiRMachine |
| CUCGGGGAUCAUCAUGUCACGAG | 429.75 | 1165.48 | RumimiR |
| UGUAAACAUCCUACACUCAGC | 3318.85 | 8846.44 | Ovine |
| GACUUACUAAGCACAGAGGUUC | 10.61 | 29.95 | RumimiR |

| | | | |
|---|---|---|---|
| CACAGCUCCAGGGGACGCCGUUC | 15.85 | 37.25 | RumimiR |
| UAGCACCAUUUGAAAUCAGUGU | 6.28 | 17.38 | Ovine |
| - | 89.59 | 219.75 | MiRDeep2 |
| UUCCUAUGCAUAUACUUCUUU | 20973.06 | 46066.56 | MiRMachine |
| UAGCACCAUUUGAAAUCAGUGU | 1268.40 | 2919.03 | Ovine |
| GAAAACCCGAAGGAACAUUUUG | 131.13 | 332.69 | RumimiR |
| GAAAAGCUGGGUUGAGAGGGCG | 3726.30 | 9278.02 | RumimiR |
| CAAUAAGUUCGUUUGGGUUUUU | 175.13 | 445.99 | RumimiR |
| AGAUGAGGCUCAGCGAGCCCUG | 290.70 | 737.01 | RumimiR |
| CAGUGCAAUGAUGAAAGGGCAU | 20367.68 | 51264.34 | MiRMachine |
| CCACGCUCAUGCACACACCCAC | 21942.44 | 54887.32 | RumimiR |
| - | 144.26 | 338.44 | MiRDeep2 |
| UAGCACCAUCUGAAAUCGGUU | 9463.28 | 23434.97 | Ovine |
| CUUCCCUUUGUCAUCCUAUGCC | 425.32 | 1040.34 | MiRMachine |
| GUCAGUGCAUGACAGAACUUG | 5451.45 | 13128.60 | MiRMachine |
| UGAAAAGUUCGUUCGGGUUUUU | 3051.74 | 7173.28 | RumimiR |
| CUGGCCCUCUCUGCCCUUCCGU | 249.08 | 563.39 | RumimiR |
| GAAUGGUGCUUUUUUGUGAAG | 84.81 | 183.25 | RumimiR |
| UUGUGACUGCUAGAACCGCUCCUG | 20.29 | 45.74 | RumimiR |
| UUAAUUUUUGCAAGGCUUUUCC | 81.57 | 176.66 | RumimiR |
| UCAAAAGUUUUAAAGCAGUCA | 3.64 | 8.68 | RumimiR |
| UUGAAAAUGACGUGACGGACUUC | 560.55 | 1250.74 | RumimiR |
| CAACUGGCCCACAAAGUCCCGC | 961.23 | 2107.89 | MiRMachine |
| GAGAGAUCAGAGGCGCAGAGU | 549.43 | 1193.88 | RumimiR |
| CCUCAGCCACACCCCUCACACA | 57.00 | 126.82 | RumimiR |
| AAAGGCCUGAAUGAACUUUUUG | 7.22 | 15.52 | RumimiR |
| GAGGGUUGGGCGGAGGCUUUCC | 1636.16 | 3404.26 | RumimiR |
| AUUGCACUUGUCUCGGUCUGA | 118420.00 | 249400.21 | Ovine |
| UUGGCUCUGCAAGGUCGGCUCA | 43.75 | 92.11 | RumimiR |
| AACACUGUCUGGUAACGAU | 881.04 | 1636.97 | Ovine |
| UCUGGUGGGAAGGAAGGGAC | 243.61 | 496.06 | RumimiR |
| AAAAGGUUCAUUUGGGUUUCU | 22.53 | 44.99 | RumimiR |
| UAAUACUGCCUGGUAAUGAUG | 1332.52 | 2587.93 | Ovine |
| UUAGGGCCCUGGCUCCAUCUC | 257.10 | 429.85 | RumimiR |
| AUACAGUGACCAGGUGACGAC | 70.45 | 133.94 | RumimiR |
| CAUGCCUUGAGUGUAGGACCGU | 8553.92 | 16931.14 | RumimiR |
| GGUACAGUACUGUGAUAACUG | 173.00 | 305.05 | MiRMachine |
| CUUUGUGUUGGCGUGUGCACCUG | 6.84 | 13.17 | RumimiR |
| UUGCUCCCUCCACACUUCCAGA | 22.48 | 41.13 | RumimiR |
| UGUGCAAAUCCAUGCAAAACUGA | 3666.45 | 6420.51 | Ovine |
| UUGUGCUUGAUCUAACCAUGU | 69.68 | 127.14 | Ovine |
| UUAUCAGAAUCUCCAGGGGUAC | 23598.88 | 43245.53 | RumimiR |
| CGUCAACACUUGCUGGUUUCCUC | 779.97 | 1453.25 | RumimiR |
| CAUCCCUUGCAUGGUGGAGG | 214.06 | 396.05 | RumimiR |
| - | 94378.71 | 170865.62 | MiRDeep2 |
| UUGUGCAAAUCUAUGCAAAACUG | 228.56 | 362.39 | MiRMachine |
| AAAAAGUUCGUUCAGGUUUUC | 22.09 | 39.25 | RumimiR |
| - | 8600.05 | 15507.54 | MiRDeep2 |
| CCAAAAAGUUUGUUUGGGUUUU | 111.07 | 195.54 | RumimiR |
| CUGUAAACAUCCUUGACUGGAAGC | 6468.55 | 11449.30 | MiRMachine |

| | | | |
|---|---|---|---|
| UCAGUGCACUACAGAACUUUGU | 12598.24 | 21694.14 | Ovine |
| AAAAUCUCUGCAGGCAAAUGUG | 30.45 | 50.66 | MiRMachine |
| AUUGGGGAAAACAGGCUCAGGGU | 12.94 | 21.02 | RumimiR |
| AAAACCUGAAUGAACUUUUUGG | 67.87 | 117.11 | RumimiR |
| UCUACAGUGCACGUGUCUCCAGU | 209.96 | 355.87 | MiRMachine |
| GUAGAGGAGAUGGCGCAGGGGACA | 285.58 | 485.74 | RumimiR |
| UUCUAGUAAGAGUGGCAGUCG | 193.54 | 325.52 | RumimiR |
| UGUGAAAUGUUUAGGACCACUA | 15.95 | 27.02 | MiRMachine |
| UAAUGCCCCUAAAAAUCCUUAU | 4475.05 | 7482.41 | MiRMachine |
| AAACUCGUGUCUGAUUCGUU | 28.00 | 44.79 | RumimiR |
| CUGCCCUGGCCCGAGGGACCGA | 945.71 | 1509.84 | RumimiR |
| AAUCUCAGGUUCGUCAGCCCGC | 981.31 | 1555.48 | MiRMachine |
| UAACACUGUCUGGUAAAGAUG | 250.74 | 351.62 | MiRMachine |
| UCUGAGACAGGAGGGCAGGUGG | 59.37 | 86.88 | RumimiR |
| UUGCUUCCUUGUGUCCACAGG | 155.76 | 248.37 | RumimiR |
| AAGGUAGAUAGAACAGGUCUUG | 1176.80 | 1850.94 | RumimiR |
| CCAAAGUGCUGUUCGUGCAGGUA | 32913.37 | 52611.03 | MiRMachine |
| GUCUCACACAGAAAUCGCACCCAU | 18587.84 | 29052.07 | RumimiR |
| AAACCGGCAUGAACUUUUUGG | 69.90 | 109.33 | RumimiR |
| CUGCCAAGCCCACGUUCAAAG | 626.02 | 970.08 | RumimiR |
| UUCACAGUGGCUAAGUUCUG | 55053.21 | 84770.72 | MiRMachine |
| UACCCAUUGCAUAUCGGAGCUG | 1331.82 | 2001.14 | RumimiR |
| CAAGAAGUUCGUUUGGGUUUUC | 30.81 | 44.88 | RumimiR |
| CUUCACCACCUUCUCCACCCAG | 1979.07 | 2751.35 | RumimiR |
| UGAUAUGUUUGAUAUUGGGUUG | 1515.89 | 2051.36 | MiRMachine |
| UGGCUCAGUUCAGCAGGAACAG | 78162.53 | 105712.00 | MiRMachine |
| UUAAAAGUUUGGUUGGGUUUUU | 164.61 | 222.87 | RumimiR |
| AACCCGUAGAUCCGAACUUGUG | 9114.51 | 11745.97 | MiRMachine |
| CAGCUGGUGUUGUGAAUCAGGCC | 491.32 | 655.90 | MiRMachine |
| UCACAGUGAACCGGUCUCUUU | 4640.23 | 6167.40 | MiRMachine |
| ACCAUCGACCGUUGAGUGGACC | 48.52 | 63.05 | MiRMachine |
| ACCUCAGUCAGCCUUGUGGAUG | 5108.93 | 6212.31 | RumimiR |
| CAGCUGGUGUUGUGAAUCAGGCC | 69.82 | 85.60 | MiRMachine |
| - | 8514.32 | 9779.55 | MiRDeep2 |
