## Supplemental Table10 for "Characterization of ovine follicular fluid and granulosa cell-derived extracellular vesicles and their miRNA cargo following in vitro exposure to bisphenols A and S"

| miR identification or sequence | log2 Fold Change (lfc) | c standard err | padj |
|---|---|---|---|
| gcgcgcgccgcggcugga | -12.87 | 2.97E+00 | 3.81E-05 |
| auugguacuucuuagagugaa | -11.99 | 1.75E+00 | 3.35E-11 |
| aaaaguucguuuggguuuuuc | -9.82 | 2.95E+00 | 1.79E-03 |
| caguuaccgcuuccgcuaccgc | -9.36 | 7.16E-01 | 1.11E-37 |
| uacaaagauaagcuaaaaguuc | -3.96 | 9.09E-01 | 3.58E-05 |
| guccuggugucugaguga | -3.49 | 1.37E+00 | 1.83E-02 |
| oar-05007 | -3.38 | 6.41E-01 | 4.14E-07 |
| oar-04641 | -3.26 | 2.91E-01 | 5.76E-28 |
| oar-05098a | -2.81 | 4.77E-01 | 1.43E-08 |
| oar-15f | -2.74 | 1.03E-01 | 4.43E-152 |
| oar-202 | -2.73 | 2.96E-01 | 3.53E-19 |
| oar-04592 | -2.63 | 2.67E-01 | 7.30E-22 |
| uacucaggaaggcauuauuca | -2.26 | 3.39E-01 | 1.26E-10 |
| oar-mir-30c | -2.22 | 3.04E-01 | 1.80E-12 |
| oar-15e | -2.12 | 1.54E-01 | 4.90E-42 |
| uacuccagagggugucauuca | -2.11 | 3.22E-01 | 2.53E-10 |
| oar-mir-19b | -2.10 | 3.05E-01 | 2.99E-11 |
| oar-33 | -2.07 | 2.61E-01 | 1.31E-14 |
| oar-30b | -2.06 | 2.31E-01 | 4.39E-18 |
| oar-34b | -2.04 | 3.29E-01 | 2.30E-09 |
| oar-05025 | -1.88 | 2.37E-01 | 1.45E-14 |
| oar-135b | -1.81 | 4.04E-01 | 2.06E-05 |
| oar-mir-150 | -1.81 | 1.49E-01 | 2.04E-32 |
| oar-145 | -1.72 | 2.31E-01 | 5.98E-13 |
| oar-190a | -1.61 | 1.17E-01 | 4.90E-42 |
| oar-05084b | -1.54 | 4.05E-01 | 3.29E-04 |
| oar-126 | -1.47 | 2.12E-01 | 2.23E-11 |
| oar-mir-30b | -1.44 | 3.10E-01 | 9.04E-06 |
| oar-04984 | -1.41 | 2.39E-01 | 1.32E-08 |
| oar-204b | -1.40 | 3.98E-01 | 9.05E-04 |
| oar-7a | -1.40 | 1.92E-01 | 1.57E-12 |
| oar-34c | -1.37 | 2.86E-01 | 4.55E-06 |
| oar-03709 | -1.33 | 2.23E-01 | 8.80E-09 |
| oar-15d | -1.31 | 1.38E-01 | 4.32E-20 |
| uaacuguggcgcaugggcuuca | -1.30 | 4.75E-01 | 1.08E-02 |
| oar-02698 | -1.30 | 2.95E-01 | 2.73E-05 |
| oar-01917 | -1.30 | 2.69E-01 | 4.19E-06 |
| oar-223 | -1.28 | 2.06E-01 | 2.47E-09 |
| oar-00673a | -1.24 | 1.59E-01 | 4.01E-14 |
| oar-210 | -1.24 | 1.48E-01 | 4.40E-16 |
| oar-00826 | -1.20 | 1.86E-01 | 5.33E-10 |
| oar-05022b | -1.19 | 1.49E-01 | 8.11E-15 |
| oar-204a | -1.19 | 1.78E-01 | 1.19E-10 |
| oar-01288 | -1.17 | 1.52E-01 | 7.64E-14 |
| oar-19 | -1.14 | 2.86E-01 | 1.49E-04 |
| oar-101a | -1.14 | 3.32E-01 | 1.27E-03 |
| oar-mir-200b | -1.12 | 1.83E-01 | 3.49E-09 |
| oar-04620 | -1.12 | 1.39E-01 | 5.19E-15 |
| oar-mir-25 | -1.11 | 9.99E-02 | 1.24E-27 |

| | | | |
|---|---|---|---|
| oar-01280 | -1.11 | 1.48E-01 | 3.51E-13 |
| oar-196a | -1.08 | 3.00E-01 | 6.89E-04 |
| oar-01605 | -1.08 | 1.30E-01 | 1.04E-15 |
| oar-02757 | -1.07 | 3.50E-01 | 4.03E-03 |
| oar-15c | -1.07 | 1.01E-01 | 8.38E-25 |
| oar-05020 | -1.06 | 1.09E-01 | 2.83E-21 |
| oar-00160 | -1.05 | 1.48E-01 | 8.13E-12 |
| oar-04478 | -1.03 | 2.44E-01 | 6.13E-05 |
| oar-mir-143 | -1.03 | 1.64E-01 | 1.53E-09 |
| oar-03097 | -1.01 | 1.54E-01 | 2.86E-10 |
| oar-05375 | -1.01 | 2.83E-01 | 8.19E-04 |
| oar-03664 | -0.98 | 3.09E-01 | 2.81E-03 |
| oar-30a | -0.97 | 1.11E-01 | 1.89E-17 |
| oar-02832a | -0.96 | 1.46E-01 | 2.35E-10 |
| oar-mir-17 | -0.94 | 8.42E-02 | 1.34E-27 |
| oar-mir-487b | -0.93 | 3.69E-01 | 2.03E-02 |
| oar-04754 | -0.91 | 3.75E-01 | 2.49E-02 |
| oar-03457 | -0.91 | 4.16E-01 | 4.59E-02 |
| oar-17b | -0.90 | 1.03E-01 | 2.67E-17 |
| oar-17a | -0.89 | 9.87E-02 | 2.21E-18 |
| oar-05004 | -0.87 | 2.23E-01 | 2.22E-04 |
| oar-00815b | -0.87 | 2.61E-01 | 1.83E-03 |
| oar-mir-30a | -0.86 | 1.19E-01 | 2.90E-12 |
| oar-mir-10b | -0.86 | 1.49E-01 | 3.45E-08 |
| oar-mir-374a | -0.85 | 2.55E-01 | 1.71E-03 |
| oar-190b | -0.84 | 3.27E-01 | 1.79E-02 |
| oar-mir-409 | -0.83 | 3.24E-01 | 1.79E-02 |
| oar-04987 | -0.81 | 2.45E-01 | 1.79E-03 |
| oar-03302 | -0.81 | 2.35E-01 | 1.17E-03 |
| oar-15a | -0.80 | 9.87E-02 | 3.17E-15 |
| oar-00945 | -0.79 | 2.77E-01 | 7.63E-03 |
| oar-01889 | -0.77 | 1.43E-01 | 2.32E-07 |
| oar-03431 | -0.76 | 2.30E-01 | 1.79E-03 |
| oar-mir-10a | -0.75 | 1.16E-01 | 3.80E-10 |
| oar-01437 | -0.74 | 3.00E-01 | 2.30E-02 |
| oar-00061 | -0.73 | 2.51E-01 | 6.30E-03 |
| oar-7c | -0.71 | 2.76E-01 | 1.74E-02 |
| oar-mir-374b | -0.69 | 1.36E-01 | 1.20E-06 |
| oar-mir-106b | -0.65 | 1.22E-01 | 3.01E-07 |
| uauugcacuugucccggccugu | -0.64 | 1.55E-01 | 8.31E-05 |
| oar-499 | -0.60 | 2.58E-01 | 3.49E-02 |
| oar-00655 | -0.59 | 1.55E-01 | 2.99E-04 |
| oar-mir-200c | -0.59 | 9.95E-02 | 1.11E-08 |
| oar-00466a | -0.54 | 1.11E-01 | 3.15E-06 |
| oar-mir-107 | -0.53 | 1.98E-01 | 1.26E-02 |
| oar-mir-30d | -0.50 | 9.40E-02 | 3.01E-07 |
| oar-mir-200a | -0.50 | 1.14E-01 | 2.73E-05 |
| oar-let-7b | -0.50 | 9.33E-02 | 3.01E-07 |
| oar-181c | -0.49 | 2.26E-01 | 4.53E-02 |
| oar-31 | -0.41 | 1.13E-01 | 7.53E-04 |

| | | | |
|---|---|---|---|
| oar-04974 | -0.36 | 9.35E-02 | 2.22E-04 |
| oar-04969 | -0.35 | 1.03E-01 | 1.32E-03 |
| ugagguaggagguuguauaguu | -0.34 | 8.57E-02 | 1.48E-04 |
| oar-140 | -0.34 | 1.28E-01 | 1.32E-02 |
| oar-00176a | -0.32 | 1.13E-01 | 7.25E-03 |
| oar-130d | -0.30 | 1.06E-01 | 7.68E-03 |
| oar-17c | -0.29 | 7.64E-02 | 3.98E-04 |
| oar-let-7c | -0.27 | 1.18E-01 | 3.35E-02 |
| oar-mir-16b | -0.27 | 8.38E-02 | 2.60E-03 |
| oar-425 | -0.22 | 8.72E-02 | 2.03E-02 |
| oar-let-7a | -0.22 | 7.71E-02 | 8.20E-03 |
| oar-04915 | -0.15 | 6.99E-02 | 4.74E-02 |

GC = Granulosa cells

| Sequence | 0h GC mean read | 48h GC mean read | Annotation |
|---|---|---|---|
| - | 342.09 | 10.13 | MiRDeep2 |
| - | 69.38 | 5.14 | MiRDeep2 |
| - | 331.49 | 0.00 | MiRDeep2 |
| - | 119.87 | 0.00 | MiRDeep2 |
| - | 57.12 | 14.79 | MiRDeep2 |
| - | 62.81 | 10.59 | MiRDeep2 |
| GAGACCCUGGUCUGCACUCUGUC | 21.89 | 3.16 | RumimiR |
| UUAGGGCCCUGGCUCCAUCUC | 211.00 | 30.40 | RumimiR |
| AAAGGUUCAUUUGUGUUUUUCU | 28.23 | 4.88 | RumimiR |
| CUAGCAGCGGGAACAGUACUGCA | 8267.54 | 1192.69 | MiRMachine |
| UUCCUAUGCAUAUACUUCUUU | 17361.94 | 2184.03 | MiRMachine |
| GUCCCUGUCCUCCAGGAGCUCAC | 5486.47 | 960.69 | RumimiR |
| - | 58.28 | 13.70 | MiRDeep2 |
| UGUAAACAUCCUACACUCUCA | 10487.17 | 2595.48 | Ovine |
| ACAGCAGCAAUUCAUGUUUUG | 22888.67 | 5028.81 | MiRMachine |
| - | 73.65 | 20.27 | MiRDeep2 |
| UGUGCAAAUCCAUGCAAAACUGA | 3020.96 | 611.11 | Ovine |
| GCAGUGCCUCGGCAGUGCAGC | 242.33 | 57.35 | MiRMachine |
| CUGUAAACACCCUACACUCUCAGC | 1877.82 | 493.99 | MiRMachine |
| CAAUCACUAGUUCCACUGCCAU | 141.38 | 30.13 | MiRMachine |
| UUUUUGCAAUAUGUUCCUGAA | 272.43 | 65.09 | RumimiR |
| CUAUGGCUUUUUAUUCCUAUGUG | 87.39 | 20.07 | MiRMachine |
| UCUCCCAACCCUUGUACCAGUG | 17895.08 | 5404.73 | Ovine |
| GGUCCAGUUUUCCCAGGAAUCCC | 553.53 | 177.43 | MiRMachine |
| UGAUAUGUUUGAUAUUGGGUUG | 1247.47 | 403.33 | MiRMachine |
| AAAAGUUCAUUUGGGUUGUUCCU | 16.31 | 5.41 | RumimiR |
| CAUUAUUACUUUUGGUACGCG | 293.93 | 99.49 | MiRMachine |
| UGUAAACAUCCUACACUCAGC | 2733.57 | 982.55 | Ovine |
| GAAUGGUGCUUUUUUGUGAAG | 69.54 | 29.24 | RumimiR |
| UUCCCUUUGUCAUCCUUUGCCC | 42.40 | 17.06 | MiRMachine |
| GUGGAAGACUAGUGAUUUUGUUGU | 62.55 | 23.77 | MiRMachine |
| UAGGCAGUGUAGUUAGCUGAUUG | 495.76 | 128.85 | MiRMachine |
| AGAUGAGGCUCAGCGAGCCCUG | 239.18 | 95.27 | RumimiR |
| CAGCAGCACACUGUGGUUUGU | 40063.90 | 16212.73 | MiRMachine |
| - | 298.34 | 130.12 | MiRDeep2 |
| UUAAUUUUUGCAAGGCUUUUCC | 67.07 | 25.66 | RumimiR |
| UUCAACGGGUAUUUAUUGAGCA | 52.67 | 22.40 | RumimiR |
| GUGUCAGUUUGUCAAAUACCCC | 10806.49 | 4320.93 | MiRMachine |
| CAAUAAGUUCGUUUGGGUUUUU | 144.42 | 61.00 | RumimiR |
| ACUGUGCGUGUGACAGCGGCUG | 7413.47 | 3200.25 | MiRMachine |
| CCACGCUCAUGCACACACCCAC | 18037.95 | 7931.07 | RumimiR |
| UUUUUGCGAUGUGUUCCUAAUA | 1389.81 | 571.60 | RumimiR |
| CUUCCCUUUGUCAUCCUAUGCC | 350.14 | 154.95 | MiRMachine |
| CAUGUCCGCGGGUUCCCUAUC | 356.74 | 157.86 | RumimiR |
| UUGUGCAAAUCUAUGCAAAACUG | 187.20 | 72.87 | MiRMachine |
| GUACAGUACUGUGAUAACUGA | 130.63 | 54.28 | MiRMachine |
| UAAUACUGCCUGGUAAUGAUG | 1100.26 | 515.16 | Ovine |
| UUGGCAUUACUGAGCAUCUAG | 1211.60 | 575.88 | RumimiR |
| AUUGCACUUGUCUCGGUCUGA | 97439.54 | 45459.32 | Ovine |

| | | | |
|---|---|---|---|
| AUCCCACUCCUGACACCA | 2319.47 | 1107.94 | RumimiR |
| UAGGUAGUUUCAUGUUGUUGG | 30.82 | 16.32 | MiRMachine |
| GCUUGGCACCUAGUAAGUACUC | 1565.54 | 752.49 | RumimiR |
| UUGCUCCCUCCACACUUCCAGA | 18.59 | 8.47 | RumimiR |
| UAGCAGCACAGAAAUGUUGGUA | 8568.62 | 3945.88 | MiRMachine |
| CUCGGGGAUCAUCAUGUCACGAG | 352.50 | 168.80 | RumimiR |
| CUUCACCACCUUCUCCACCCAG | 1626.01 | 769.32 | RumimiR |
| UUGGCUCUGCAAGGUCGGCUCA | 36.00 | 17.90 | RumimiR |
| UGAGAUGAAGCACUGUAGCUC | 1052.71 | 508.84 | Ovine |
| GAGGGUUGGGCGGAGGCUUUCC | 1350.15 | 691.26 | RumimiR |
| UAAAAAGGUCUUUCGAGUUUUU | 36.28 | 17.44 | RumimiR |
| CCUCAGCCACACCCCUCACACA | 46.86 | 24.67 | RumimiR |
| CUGUAAACAUCCUUGACUGGAAGC | 5327.08 | 2693.00 | MiRMachine |
| CCACAUGGAGUUGCUGUUACAA | 178.27 | 92.63 | RumimiR |
| CAAAGUGCUUACAGUGCAGGUA | 71542.46 | 37639.83 | Ovine |
| AAUCGUACAGGGUCAUCCACUU | 23.35 | 15.10 | Ovine |
| UCCUCUCUCGGUUAGCUCCAUA | 18.10 | 10.55 | RumimiR |
| GACCCUGACGGGCGUGGAUUGU | 15.59 | 8.09 | RumimiR |
| CCAAAGUGCUGUUCGUGCAGGUA | 27113.03 | 14874.23 | MiRMachine |
| CUAAAGUGCUUAUAGUGCAGGUA | 58142.63 | 31314.94 | MiRMachine |
| CGUCAACACUUGCUGGUUUCCUC | 641.41 | 378.64 | RumimiR |
| AAAACCCGCAUGAACUUUUUGG | 66.64 | 38.98 | RumimiR |
| CUUUCAGUCGGAUGUUUGCAG | 2574.86 | 1466.03 | Ovine |
| ACCCUGUAGAACCGAAUUUGUG | 6683.05 | 3773.52 | Ovine |
| UUAUAAUACAACCUGAUAAGUG | 2355.71 | 1183.70 | Ovine |
| UGAUAUGUUUGAUAUAUUAGGUU | 58.09 | 27.88 | MiRMachine |
| CGAAUGUUGCUCGGUGAACCCCU | 25.75 | 15.33 | Ovine |
| GAUUGGCACGUCUUGGAAUG | 127.21 | 78.25 | RumimiR |
| CUGAGACCUCCGGGUUCUGAGC | 103.24 | 58.23 | RumimiR |
| UAGCAGCACAUCAUGGUUUACA | 104813.20 | 60779.06 | MiRMachine |
| AUACAGUGACCAGGUGACGAC | 57.91 | 29.98 | RumimiR |
| UUAAAAGUUUGGUUGGGUUUUU | 135.30 | 80.42 | RumimiR |
| UCAGUCAUUCCGUUGUGUCCGA | 44.68 | 26.79 | RumimiR |
| UACCCUGUAGAUCCGAAUUUG | 490.97 | 285.62 | Ovine |
| UUGUGUCAAUAUGCGAUGAUGU | 42.43 | 23.11 | RumimiR |
| AAAACCUGAAUGAACUUUUUGG | 55.91 | 31.65 | RumimiR |
| CUGGAAGACUAGUGAUUUUGUUGU | 35.61 | 22.53 | MiRMachine |
| AUAUAAUACAACCUGCUAAGU | 14811.34 | 9362.43 | Ovine |
| UAAAGUGCUGACAGUGCAGAU | 30160.61 | 18591.07 | Ovine |
| - | 77648.57 | 48802.05 | MiRDeep2 |
| GUUAAGACUUGCAGUGAUGUU | 46.47 | 29.48 | MiRMachine |
| UAGUGGGGAACCCUUCCAUGAGG | 604.28 | 390.71 | RumimiR |
| UAAUACUGCCGGGUAAUGAUGG | 662.46 | 444.01 | Ovine |
| GAAAACCCGAAGGAACAUUUUG | 108.15 | 74.10 | RumimiR |
| AGCAGCAUUGUACAGGGCUAUC | 3686.95 | 2818.07 | Ovine |
| UGUAAACAUCCCCGACUGG | 14237.53 | 10055.51 | Ovine |
| AACACUGUCUGGUAACGAU | 729.16 | 486.14 | Ovine |
| UGAGGUAGUAGGUUGUGUGGU | 110193.51 | 76982.63 | Ovine |
| ACCAUCGACCGUUGAGUGGACC | 40.15 | 28.43 | MiRMachine |
| AGGCAAGAUGCUGGCAUAGCUG | 565160.15 | 423618.93 | MiRMachine |

| | | | |
|---|---|---|---|
| ACCUCAGUCAGCCUUGUGGAUG | 4205.58 | 3275.61 | RumimiR |
| CAACUGUUUGCAGAGGAAACUGA | 1378.45 | 1095.51 | RumimiR |
| - | 22484.55 | 17715.77 | MiRDeep2 |
| CCAGUGGUUUUACCCUAUGGUA | 3792.61 | 2888.03 | MiRMachine |
| UUGAAAAUGACGUGACGGACUUC | 461.89 | 367.20 | RumimiR |
| CAGUGCAAUGAUGAAAGGGCAU | 16802.98 | 13707.99 | MiRMachine |
| UUAAGGUGCAUCUAGUGCA | 12796.94 | 10412.98 | MiRMachine |
| UGAGGUAGUAGGUUGUAUGGUU | 21103.02 | 17306.51 | Ovine |
| UAGCAGCACGUAAAUAUUGG | 466943.35 | 387283.36 | Ovine |
| GAAUGACACGAUCACUCCCGUUGAG | 4245.65 | 3677.82 | MiRMachine |
| UGAGGUAGUAGGUUGUAUAGUU | 325125.26 | 279501.48 | Ovine |
| CAUGCCUUGAGUGUAGGACCGU | 7043.13 | 6379.09 | RumimiR |
