## Supplemental Table11 for "Characterization of ovine follicular fluid and granulosa cell-derived extracellular vesicles and their miRNA cargo following in vitro exposure to bisphenols A and S"

| miR identification or sequence | log2 Fold Change (lfc) | lfc standard error | padj |
|---|---|---|---|
| agaguucauucagguuuuucu | 8.184307817 | 2.208521917 | 0.00046443 |
| caguuaccgcuuccgcuaccg | 6.825698697 | 0.813904151 | 3.9428E-16 |
| ggaugccuggggguccugg | 6.663544051 | 0.835274 | 9.7347E-15 |
| aauagcucagaaugucacuucu | 6.532704805 | 2.957682521 | 0.0439676 |
| caccucaucccagcagugguc | 5.652218513 | 1.070897754 | 4.1453E-07 |
| uugcacguggaagagaaacaga | 5.204963441 | 1.006931565 | 7.3258E-07 |
| cgcuggcaguguugguggcacu | 5.115076943 | 1.15369792 | 2.4852E-05 |
| caauaugaucaaaugugccaga | 4.741951319 | 0.91763366 | 7.3258E-07 |
| ccugcuugggcucuccucuaga | 4.247547331 | 1.889074303 | 0.04006019 |
| aauugagguguaguuagcaaaca | 4.144069165 | 0.645208329 | 5.7135E-10 |
| oar-147 | 4.120890492 | 0.210672144 | 2.3153E-83 |
| oar-155 | 4.062273058 | 0.250147213 | 9.1672E-58 |
| oar-129a | 4.011216655 | 0.619542659 | 4.2751E-10 |
| ccagggcagccugugguaacagu | 3.719319755 | 1.081729534 | 0.00121278 |
| oar-04143 | 3.708684844 | 0.707811124 | 5.0604E-07 |
| oar-146b | 3.436989074 | 0.194294601 | 2.1821E-68 |
| oar-02648b | 3.371369721 | 0.730933039 | 1.0929E-05 |
| aaacaagaucacgccucucaga | 2.98746354 | 0.70124956 | 5.2347E-05 |
| oar-146a | 2.973104136 | 0.148302862 | 1.8353E-87 |
| oar-455 | 2.944375913 | 0.160755737 | 3.0581E-73 |
| oar-221 | 2.822759518 | 0.174523011 | 2.4145E-57 |
| oar-05084a | 2.472204755 | 0.497236797 | 1.9442E-06 |
| oar-00629 | 2.390065048 | 0.465924159 | 8.73E-07 |
| oar-24b | 2.321439617 | 0.179465804 | 6.1545E-37 |
| oar-02648a | 2.283412661 | 0.162811978 | 3.1674E-43 |
| oar-mir-221 | 2.179590906 | 0.108341498 | 5.9467E-88 |
| oar-mir-27a | 2.076783274 | 0.127100787 | 1.9815E-58 |
| oar-00430 | 2.02515034 | 0.247278327 | 1.9269E-15 |
| oar-06131 | 2.009955475 | 0.355710366 | 5.5341E-08 |
| oar-04920 | 1.975907969 | 0.347024264 | 4.385E-08 |
| oar-01908 | 1.974061261 | 0.193354756 | 2.4875E-23 |
| oar-128b | 1.941926444 | 0.474597323 | 0.00010433 |
| oar-00466c | 1.937088347 | 0.470089079 | 9.2696E-05 |
| oar-mir-22 | 1.925437016 | 0.223227932 | 5.1379E-17 |
| oar-04560b | 1.844597774 | 0.164552748 | 6.6497E-28 |
| oar-02825 | 1.806442603 | 0.44917585 | 0.00013884 |
| oar-04145 | 1.776398755 | 0.2772479 | 6.1763E-10 |
| oar-00805 | 1.774402071 | 0.313528188 | 5.3069E-08 |
| oar-04885 | 1.719843401 | 0.266148969 | 4.5268E-10 |
| oar-01337 | 1.710186515 | 0.568283905 | 0.00492251 |
| oar-04203 | 1.670745392 | 0.179550081 | 1.3613E-19 |
| oar-01279 | 1.59985689 | 0.315010974 | 1.1333E-06 |
| oar-04130 | 1.573449449 | 0.163281997 | 6.4732E-21 |
| oar-mir-23a | 1.409985834 | 0.131665744 | 1.4552E-25 |
| oar-203 | 1.409233846 | 0.347060461 | 0.00011849 |
| oar-03788 | 1.392194504 | 0.058871109 | 2.132E-121 |
| oar-181d | 1.382489526 | 0.14679919 | 4.841E-20 |
| oar-34a | 1.368440899 | 0.227537135 | 6.9523E-09 |
| acuggacuuggagucagaaggc | 1.364297728 | 0.144238227 | 3.482E-20 |

| | | | |
|---|---|---|---|
| oar-04560a | 1.353826476 | 0.152135664 | 5.008E-18 |
| oar-00496 | 1.302663644 | 0.30331641 | 4.5161E-05 |
| oar-03006 | 1.288379434 | 0.27193186 | 6.0753E-06 |
| oar-01301 | 1.248177884 | 0.143888917 | 3.4191E-17 |
| oar-00311 | 1.200612949 | 0.122425965 | 1.3481E-21 |
| oar-00342 | 1.19849236 | 0.061331913 | 2.8328E-83 |
| oar-01446 | 1.164282315 | 0.167148189 | 1.7411E-11 |
| oar-24a | 1.133888985 | 0.109213547 | 4.3028E-24 |
| cacauggaguugcuguuaca | 1.124328825 | 0.378377538 | 0.00548419 |
| oar-193a | 1.109491575 | 0.173058171 | 6.0965E-10 |
| oar-130c | 1.106555641 | 0.183153136 | 5.9287E-09 |
| oar-mir-21 | 1.100978395 | 0.213668299 | 7.86E-07 |
| oar-04836 | 1.049463924 | 0.265303575 | 0.00017961 |
| oar-92b | 1.0389248 | 0.184542168 | 6.1828E-08 |
| oar-00238 | 1.034541448 | 0.365404379 | 0.00827017 |
| oar-122 | 1.024286264 | 0.467485676 | 0.04530737 |
| oar-193c | 1.012277432 | 0.13559135 | 4.861E-13 |
| oar-04334 | 1.005570276 | 0.205592087 | 2.9156E-06 |
| oar-01519 | 1.004114351 | 0.38698908 | 0.01637928 |
| oar-130e | 0.989904539 | 0.256053303 | 0.0002535 |
| oar-02650 | 0.987487405 | 0.164738051 | 7.7709E-09 |
| oar-132a | 0.95856445 | 0.148229658 | 4.4417E-10 |
| oar-9c | 0.948015387 | 0.254454297 | 0.00043203 |
| oar-130a | 0.94616437 | 0.172094263 | 1.2908E-07 |
| oar-05062 | 0.929489222 | 0.114590775 | 3.534E-15 |
| oar-01491 | 0.913383931 | 0.177772419 | 8.4305E-07 |
| oar-129b | 0.896074716 | 0.318111087 | 0.00860484 |
| oar-03461 | 0.816187634 | 0.258320325 | 0.00305386 |
| oar-mir-106a | 0.774609319 | 0.13161498 | 1.431E-08 |
| uggagagaaaggcaguuccuga | 0.748115368 | 0.111751686 | 1.0856E-10 |
| oar-03296 | 0.745446946 | 0.165783691 | 1.882E-05 |
| oar-181b | 0.72572175 | 0.242606856 | 0.00516671 |
| oar-04905 | 0.721856723 | 0.078311857 | 3.0002E-19 |
| oar-03468 | 0.720620352 | 0.217091746 | 0.00179368 |
| oar-130b | 0.719956071 | 0.154662605 | 9.0351E-06 |
| oar-02968 | 0.718391661 | 0.264250225 | 0.01151426 |
| oar-mir-29b-1b | 0.715097601 | 0.199058768 | 0.00070855 |
| oar-04983b | 0.703605884 | 0.222655519 | 0.00305386 |
| oar-mir-103 | 0.680822809 | 0.069891014 | 2.4857E-21 |
| oar-181a | 0.663212484 | 0.082229607 | 5.0525E-15 |
| oar-132b | 0.654838308 | 0.22928337 | 0.00773068 |
| oar-139 | 0.644833472 | 0.103006647 | 1.5659E-09 |
| oar-04983a | 0.644733781 | 0.209592749 | 0.0040088 |
| oar-193b | 0.635628257 | 0.169762283 | 0.00040393 |
| oar-490 | 0.575838072 | 0.155821535 | 0.00048058 |
| oar-01635 | 0.546733134 | 0.174601236 | 0.00334505 |
| oar-01269 | 0.519633848 | 0.14931941 | 0.00105139 |
| oar-mir-181a-2 | 0.512065964 | 0.078418967 | 3.0367E-10 |
| oar-10a | 0.511529422 | 0.122193296 | 7.1112E-05 |
| gaaacuggaacgaacuuuuugg | 0.470877177 | 0.134630017 | 0.00099059 |

| | | | |
|---|---|---|---|
| oar-mir-194 | 0.46845512 | 0.083607418 | 7.145E-08 |
| oar-mir-191 | 0.462324289 | 0.111495081 | 8.3402E-05 |
| oar-04906 | 0.459739361 | 0.116983298 | 0.00019862 |
| oar-02091 | 0.44582674 | 0.14595924 | 0.0042629 |
| oar-01510 | 0.444021978 | 0.202478165 | 0.04530737 |
| oar-186 | 0.406563422 | 0.135350259 | 0.00498692 |
| oar-mir-29a | 0.348709027 | 0.147487532 | 0.02990296 |
| oar-142 | 0.332611412 | 0.078466149 | 5.7141E-05 |
| oar-02031 | 0.292560899 | 0.131981848 | 0.04328296 |
| oar-05078 | 0.195462161 | 0.088889729 | 0.04487218 |

GC = Granulosa cells

| Sequence | 0h GC mean read | 48h GC mean read | Identification |
|---|---|---|---|
| | 0 | 81.74907409 | MiRDeep2 |
| | 0 | 27.31912728 | MiRDeep2 |
| | 0 | 25.27762095 | MiRDeep2 |
| | 0 | 45.86522178 | MiRDeep2 |
| | 0 | 14.46311402 | MiRDeep2 |
| | 2.782237402 | 23.68954722 | MiRDeep2 |
| | 0.756277814 | 17.04831623 | MiRDeep2 |
| | 4.558217348 | 38.20954386 | MiRDeep2 |
| | 5.767041586 | 30.93854637 | MiRDeep2 |
| | 17.83214283 | 148.2532084 | MiRDeep2 |
| GUGUGCGGAAAUGCUUCUGCU | 68.62880207 | 1273.916905 | MIRMA0046 |
| UUAAUGCUAAUCGUGAUAGGGGUU | 576.4767587 | 11408.16683 | MIRMA0009 |
| UCUUUUUGCGGUCUGGGCUU | 3.563770203 | 60.06250923 | MIRMA0032a |
| | 4.316742258 | 22.11788521 | MiRDeep2 |
| UAGCCAGUUGGGGAAGAAUGC | 2.301350463 | 25.9265638 | RUM04143 |
| CUGAGAACUGAAUUCCAUAGGCUG | 5446.360402 | 61783.50929 | MIRMA0101b |
| ACACAGGUUUUGCUUCCAUCAC | 1.027594711 | 9.89204779 | RUM02648b |
| | 6.900421453 | 25.56822233 | MiRDeep2 |
| UUGAGAACUGAAUUCCAUAGGU | 2899.1944 | 21369.48119 | MIRMA0039a |
| GCAGUCCAUGGGCAUAUACA | 1589.280818 | 12081.65719 | MIRMA0016 |
| AGCUACAUCUGGCUACUGGGUCU | 938.8672034 | 6914.068572 | MIRMA0113 |
| AAAAGUUCAUUUGGGUUGUUCCU | 2.402420686 | 14.0914338 | RUM05084a |
| UAGAAAGUUUCUUUGGGGUUUU | 2.770759354 | 14.28637754 | RUM00629 |
| CUGGCUCAGUUCAGCAGGAACAG | 38.43829798 | 193.6417408 | MIRMA0037b |
| ACACAGGUUUUGCUUCCAUCAC | 70.74068162 | 337.6138041 | RUM02648a |
| AGCUACAUUGUCUGCUGGGUUU | 9947.351321 | 45501.06227 | MI0025269 |
| UUCACAGUGGCUAAGUUCCGC | 24749.68638 | 104488.5645 | MI0025275 |
| GUCCCCGAGUCCCUGGCGUGCAC | 32.70080652 | 124.846204 | RUM00430 |
| GUUCUCCUAUAGAUGCCGUCACA | 4.885172933 | 18.85288291 | RUM06131 |
| UCUGGAGGGGCAGAAGGAGAAG | 39.40296229 | 142.7171203 | RUM04920 |
| GUGGUUGAUUGGAUCCGUGGGU | 240.1802843 | 1009.526911 | RUM01908 |
| CUCACAGUGAACCGGUCUCUU | 6.254614306 | 23.52343359 | MIRMA0093b |
| AAAAAGUUCCUUUGGGUUUU | 4.378937394 | 15.19512983 | RUM00466c |
| AAGCUGCCAGUUGAAGAACUG | 1944.401306 | 8191.344476 | MI0025268 |
| UGCGGGAUCUUUAGUUGUGGCG | 4967.096025 | 16802.99629 | RUM04560b |
| AAAAAACCGAGUGAACUUUUUG | 5.599791028 | 17.53223548 | RUM02825 |
| CAUAGCCAGUUGGGGAAGAAUG | 24.76825252 | 87.50502517 | RUM04145 |
| AUUGUCCUUGCUGUUUGGAGAU | 16.12867398 | 52.05810365 | RUM00805 |
| CUCCGUUUGCCUGUUUUGCUGA | 19.94465279 | 62.53463077 | RUM04885 |
| CACUGGAGUUUUGUUUCAACAUU | 4.704113936 | 13.53067306 | RUM01337 |
| CAAGGAGCUUACAAUCUAGCUG | 828.6968214 | 2778.627781 | RUM04203 |
| GAAGGGACCCGAACGAACUUUUU | 13.82007803 | 38.64371246 | RUM01279 |
| GUAGAGGAGAUGGCGCAGGGGACA | 235.6366555 | 683.4031084 | RUM04130 |
| AUCACAUUGCCAGGGAUUUCCA | 89576.89205 | 238498.8969 | MI0025270 |
| UGUGAAAUGUUUAGGACCACUA | 13.15124735 | 33.83281062 | MIRMA0091 |
| AAGGUAGAUAGAACAGGUCUUG | 969.7337098 | 2557.969407 | RUM03788 |
| AACAUUCAUUGCUGUCGGUGGGUU | 65.24457081 | 172.8456466 | MIRMA0069d |
| UUGGCAGUGUCUUAGCUGGUUGU | 2158.283839 | 5566.373466 | MIRMA0066a |
| | 7086.628478 | 19407.68345 | MiRDeep2 |

| | | | |
|---|---|---|---|
| UUGCGGGAUCUUUAGUUGUGG | 1200.152879 | 2930.018388 | RUM04560a |
| AAGUUCAUUCGGGUUUUUCCA | 13.50362791 | 34.87190586 | RUM00496 |
| CAGAGAGUUCAUUCGGGUUUUUC | 25.97949912 | 61.59226609 | RUM03006 |
| ACGCCCUUCCCCCCCUUCUUCA | 212.2440861 | 502.3824704 | RUM01301 |
| GAGAGAUCAGAGGCGCAGAGU | 453.0945733 | 1033.10519 | RUM00311 |
| AAGGAGCUCACAGUCUAUUGAG | 9062.291297 | 20776.45628 | RUM00342 |
| GUCAAGAGCAAUAACGAAAAAU | 1197.059498 | 2592.966589 | RUM01446 |
| UGGCUCAGUUCAGCAGGAACAG | 64317.37004 | 143087.276 | MIRMA0017a |
| | 17.97085518 | 34.12036008 | MiRDeep2 |
| UGGGUCUUUGCGGGCGAGAUGA | 2176.199549 | 4408.695478 | MIRMA0052a |
| GCAGUGCAAUGUUAAAAGGGC | 2783.221792 | 6315.59714 | MIRMA0082c |
| UAGCUUAUCAGACUGAUGUUGAC | 195201.0232 | 378941.5231 | MI0014116 |
| CUUUUCCAAUGAGUCAGUUAU | 79.16901226 | 163.4652119 | RUM04836 |
| AAAUUGCACGGUAUCCAUCUGC | 76.32357574 | 153.0183652 | MIRMA0112b |
| GUCAGAGCUGGGUUUAUUGCGUG | 16.40837851 | 30.50150171 | RUM00238 |
| GUGGAGUGUGACAAUGGUGUU | 5.769367459 | 10.68146313 | MIRMA0104 |
| CAACUGGCCCACAAAGUCCCGC | 789.2722641 | 1569.150111 | MIRMA0105c |
| ACUCGGCGUGGCGUCGGUCGUG | 760.3480752 | 1502.643386 | RUM04334 |
| CGCCCUCUGCCCGUGUCGCCAGG | 6.194565319 | 12.33559256 | RUM01519 |
| CAGUGCAAUGAUAUUGUCAAAGC | 114.9373244 | 223.8778839 | MIRMA0088e |
| CGCAUCCCCUAGGGCAUUGGUGU | 1987.260957 | 3956.211269 | RUM02650 |
| UAACAGUCUACAGCCAUGGUCG | 547.6540584 | 945.8285819 | MIRMA0060a |
| CUCUUUGGUUAUCUAGCUGUAUG | 18.22965777 | 35.11403963 | MIRMA0090c |
| UAGUGCAAUAUUGCUUAUAGGGUU | 248.9446926 | 486.1110718 | MIRMA0057a |
| AAAACCCGAACGAGCUCUUUGG | 232.9411315 | 444.0033173 | RUM05062 |
| GAGGAAGCCCUGGAGGGGCUGGAG | 659.620961 | 1243.703179 | RUM01491 |
| UCUUUUUGCGGUCUGGGCUU | 50.23510454 | 114.1173384 | MIRMA0081b |
| CCUCUGGGCCCUUCCUCCAGC | 214.5797115 | 402.4696736 | RUM03461 |
| AAAAGUGCUUACAGUGCAGGU | 510.0949865 | 867.458117 | MI0025282 |
| | 7000.296376 | 11896.49529 | MiRDeep2 |
| GCAAAGCACACGGCCUGCAGAGA | 136.2158646 | 234.6536895 | RUM03296 |
| AACAUUCAUUGUUGUCGGUGGGUU | 28.41047995 | 45.11739062 | MIRMA0040b |
| UACCCAUUGCAUAUCGGAGCUG | 1096.371346 | 1818.468693 | RUM04905 |
| ACUUCUGUCCGUCUUGCUACC | 27.87346205 | 44.66693486 | RUM03468 |
| CAGUGCAAUAGUAUUGUCAAAG | 465.9133179 | 749.5468471 | MIRMA0058b |
| AAGAGGUUCAUUCAGGUUUUC | 25.81299365 | 44.42103898 | RUM02968 |
| UAGCACCAUUUGAAAUCAGUGU | 1045.602613 | 1587.787476 | MI0025276b |
| AAGAACCUGAACGAACCUUUUGG | 34.06147745 | 52.37545363 | RUM04983b |
| AGCAGCAUUGUACAGGGCUAUG | 45187.37009 | 72222.5287 | MI0025248 |
| AACAUUCAUUGCUGUCGGUGGGUU | 889.1169336 | 1429.624041 | MIRMA0027a |
| UAACAGUCUCCAGUCACGGCC | 62.20250356 | 87.10218765 | MIRMA0061b |
| UCUACAGUGCACGUGUCUCCAGU | 173.1944191 | 268.7569191 | MIRMA0084 |
| CAAGAAGUUCGUUUGGGUUUUC | 25.32297191 | 40.77848289 | RUM04983a |
| UAAUGCCCCUAAAAAUCCUUAU | 3681.4618 | 5918.431017 | MIRMA0059b |
| ACCAUGGAUCUCCAGGUGGG | 71.70488166 | 106.0896827 | MIRMA0033 |
| CUGCCCUGGCCCGAGGGACCGA | 777.0050166 | 1093.348335 | RUM01635 |
| UUGCUUCCUUGUGUCCACAGG | 128.0390873 | 181.2977198 | RUM01269 |
| AACAUUCAACGCUGUCGGUGAGU | 11253.48235 | 16156.9706 | MI0025259 |
| CCACCCGUAGAACCGACCUUG | 6792.666707 | 9461.313187 | MIRMA0076a |
| | 303.5061173 | 419.850039 | MiRDeep2 |

| | | | |
|---|---|---|---|
| UGUAACAGCAACUCCAUGUGGA | 3310.665412 | 4560.43578 | MI0025261 |
| CAACGGAAUCCCAAAAGCAGCU | 121129.1605 | 164345.0756 | MI0025260 |
| AUGCACCUGGGCAAGGAUUCUGA | 1667.665545 | 2235.557461 | RUM04906 |
| CAAAAACUGGCAGCUUCAUGUA | 147.9754015 | 204.2933184 | RUM02091 |
| AUUAUAAAGCAAUGAGACUGAU | 532.0157334 | 637.9208533 | RUM01510 |
| CAAAGAAUUCUCCUUUUGGGCU | 9227.78208 | 12221.13755 | MIRMA0004 |
| UAGCACCAUCUGAAAUCGGUU | 7781.707 | 9918.024457 | MI0014117 |
| CCCAUAAAGUAGAAAGCACU | 14715.36735 | 18833.0892 | MIRMA0056 |
| UUCUAGUAAGAGUGGCAGUCG | 159.5420415 | 194.1973767 | RUM02031 |
| AAUGGCGCCACUAGGGUUGUGC | 8612.239138 | 9835.080222 | RUM05078 |
