## Supplemental Table12 for "Characterization of ovine follicular fluid and granulosa cell-derived extracellular vesicles and their miRNA cargo following in vitro exposure to bisphenols A and S"

| miRNA identification | Target number | Alignment number |
|---|---|---|
| oar-24b | 35675 | 166931 |
| oar-01635 | 27957 | 111029 |
| guccagcagggggagcca | 36856 | 126357 |
| uccuccuccucuccacca | 36412 | 150014 |
| oar-03746 | 37416 | 190537 |
| ugagguaguaaguuguauuguu | 35125 | 139245 |
